# FAK Inhibition Remodels the Metastatic ECM and Restores CD8⁺ T Cell Trafficking and Immunosurveillance

**DOI:** 10.64898/2026.01.21.700837

**Authors:** Nicole D. Barth, Bo Peng, Michael Papanicolaou, Fatemeh Ahmadi Moughari, Camille L. Duran, Christopher S. McGinnis, John C. Dawson, Emily R. Webb, Morwenna Muir, Fraser Laing, Yookyung Jung, Xue-Yan He, Ansuman T. Satpathy, Mikala Egeblad, Syed Haider, Rachael Natrajan, Maja H. Oktay, David Entenberg, Valerie G. Brunton

## Abstract

Metastatic breast cancer remains largely incurable, driven in part by immunosuppressive microenvironments that limit CD8^+^ T cell-mediated clearance. Using a murine pulmonary metastatic breast cancer model, we show that the focal adhesion kinase (FAK) inhibitor VS-4718 promotes a CD8^+^ T cell-dependent regression of metastatic lesions by reprograming the metastatic microenvironment. VS-4718 reduced immunosuppressive myeloid and regulatory T cells while increasing CD8^+^ T cell infiltration. Cellular and secreted proteome profiling revealed that VS-4718 downregulates ECM components such as laminin α5 and collagen VIIIα1, which we show impair CD8^+^ T cell migration and activity. In human breast cancer cohorts, elevated *LAMA5/COL8A1* expression and a FAK-dependent ECM signature associate with poor outcome and prognostic for residual disease. Intravital imaging demonstrated that VS-4718 enhances CD8⁺ T cell extravasation and induces T cell-tumor cell contacts necessary for cytotoxicity. *Ex vivo* lung slice cultures recapitulated these findings, showing enhanced T cell swarming, metastatic cluster shrinkage, and apoptosis. These findings reveal how FAK inhibition remodels the metastatic ECM to potentiate coordinated CD8^+^ T cell responses. VS-4718 might aid in clearing metastases in breast cancer patients through modulating both stromal and immune components.

**STATEMENT OF SIGNIFICANCE:** Focal adhesion kinase (FAK) inhibition remodels collagen- and laminin-rich extracellular matrix barriers and alleviates physical constraints that limit CD8⁺ T cell access and activity in metastases. This enhances infiltration, migration, and tumor cell engagement, and synergizes with PD-1 blockade, supporting combined therapeutic strategies in metastatic breast cancer.

## INTRODUCTION

Despite continuous advances, metastatic breast cancer remains largely incurable, as current therapies rarely prevent disease progression and relapse ^1,2^. The metastatic tumor microenvironment (TME), composed of stromal cells and extracellular matrix (ECM) components, shelters disseminated tumor cells from immune surveillance and therapy. Beyond serving as structural scaffolds, ECM and stromal remodeling actively direct the fate of metastatic tumor cells by influencing the ability of immune cells to infiltrate, migrate, engage with, and eliminate cancer cells ^3–5^.

In metastatic breast cancer, tumor-derived signals reprogram stromal cells to remodel and stiffen the ECM, creating a fibrotic and immunosuppressive niche that facilitates metastasis and limits therapeutic efficacy ^6–9^. Thus, modulating this remodeled metastatic TME has the potential to restore both the infiltration and interaction of immune cells with cancer cells and ultimately improve clearance of metastatic breast cancer lesions. CD8⁺ T cells are key mediators of anti-tumor immunity and their engagement with tumor cells is essential for effective cytolytic activity and tumor clearance ^10^. However, they encounter significant physical and biochemical barriers within the metastatic TME that limit their capacity to patrol and eliminate cancer cells ^3^. T cell motility depends on the composition of both stromal and tumor compartments, with migration readily occurring in loose matrices in a chemokine-dependent manner ^5^, but becoming restricted and integrin-dependent in collagen-rich fibrotic regions ^5,11^. Even in the presence of high levels of chemoattractants, CD8^+^ T cell movement is dictated by the ECM, confining them to the stroma by preventing infiltration into tumor nests ^5^ and markedly reducing T cell-cancer cell interactions and subsequent cytotoxic activity ^5^. In addition to physically restricting migration, ECM components also biochemically suppress T cells through inhibitory receptors like Leukocyte-Associated Immunoglobulin-like Receptor 1 (LAIR-1), which transmits inhibitory signals in T cells upon binding to collagen ^12^, leading ultimately to impaired cytotoxicity ^13,14^.

FAK is a non-receptor tyrosine kinase that integrates signals from integrins and growth factor receptors to orchestrate matrix remodeling and tumor-stroma communication ^15^. We and others have shown that FAK actively drives ECM reorganization and desmoplasia, correlating with poor CD8^+^ T cell infiltration and immune exclusion ^16–18^. Furthermore, FAK signaling in tumor cells also promotes immunosuppression by upregulating chemokines and cytokines that attract regulatory T cells and macrophages, while excluding cytotoxic lymphocytes ^19^. Additionally, nuclear FAK also suppresses antigen presentation and promotes immune evasion ^20^. Pharmacological FAK inhibition disrupts these tumor-stroma interactions, reduces matrix stiffness, and restores anti-tumor immunity, leading to regression in multiple solid tumor models ^17,19–22^.

In breast cancer, elevated FAK expression is observed in tumors with more aggressive clinicopathologic features, such as lymphovascular invasion, triple-negative subtype, and reduced progression-free and overall survival ^23,24^. Following dissemination, metastatic breast cancer cells rely on FAK and integrin β1 signaling to engage the ECM and proliferate within distant organs such as the lung ^25,26^.

However, the specific mechanisms by which FAK regulates the metastatic immune environment, particularly regarding CD8⁺ T cell dynamics within established metastatic lesions, remain unclear. As FAK is known to be involved in ECM remodeling, which can influence T cell-tumor interactions critical for eradication of lesions, we hypothesized that FAK governs secondary site immune cell dynamics by altering ECM and suppressing immune surveillance, thereby reducing immune cell trafficking, tumor engagement, and ultimately, immune-mediated clearance of metastatic breast cancer.

Here, we demonstrate that FAK inhibition elicits CD8⁺ T cell-dependent regression of lung metastases by remodeling the lung metastatic ECM and immune landscape. FAK inhibitor treatment reduces immunosuppressive myeloid and regulatory T cell populations while downregulating key ECM components, notably laminin α5 and collagen VIIIα1, which we show directly impact CD8⁺ T cell motility and activity. Using intravital imaging, *ex vivo* lung slice co-cultures, and cytotoxicity assays, we demonstrate that FAK inhibitor treatment drives CD8⁺ T cell extravasation and sustained, antigen-specific tumor interactions, effects further amplified by PD-1 blockade. While FAK inhibition alone improves CD8⁺ T cell access and reduces motility, combination therapy drives CD8⁺ T cell arrest, directional persistence, and frequency of tumor cell engagement, resulting in reduced metastatic proliferation and increased tumor cell apoptosis. Together, these findings uncover a novel mechanism by which FAK inhibition remodels the ECM and reprograms the metastatic immune microenvironment to potentiate anti-tumor immunity.

## RESULTS

### FAK inhibition with VS-4718 elicits immune-mediated suppression of metastasis

To investigate the potential of FAK inhibition to reduce metastatic burden, we utilized VS-4718 in preclinical models of breast cancer ^27^. VS-4718 is one of the best-characterized FAK inhibitors in preclinical models and has previously been reported to alter the immune environment in primary tumors ^17,19,28^. We engineered MMTV-PyMT derived tumor cells to stably express the Cerulean2 (Cer2) fluorescent reporter and to present the chicken ovalbumin (OVA) peptide, facilitating *in vivo* tracking of tumor cells and antigen-specific T cell responses (**Supplementary Fig. 1A/B)** ^29,30^. Flow cytometry confirmed that Cer2-OVA MMTV-PyMT (Cer2-OVA MMTV-PyMT) cells presented the fragmented chicken ovalbumin peptide SIINFEKL on their surface via MHC class I molecules (**Supplementary Fig. 1B**). We further confirmed that VS-4718 inhibits phosphorylation of FAK in Cer2-OVA MMTV-PyMT cells (**Supplementary Fig. 1C**). We initially evaluated the impact of VS-4718 on pulmonary metastatic burden in mice following a published experimental metastasis protocol ^31^ (**Fig. 1A**). Quantification of metastatic burden was assessed in H&E stains and immunofluorescence stains for the Cer2 fluorophore (**Fig. 1B**). VS-4718 treatment markedly reduced metastatic burden, decreasing the percentage of lung area occupied by metastases from 0.7% in vehicle-treated mice to 0.05% in VS-4718-treated mice (**Fig. 1C**). To validate the broader relevance of our findings, we confirmed that FAK inhibition (**Supplementary Fig. 2A&B**) also reduced metastatic burden in the 4T1 model (**Supplementary Fig. 2 C-E**), a distinct and widely used model of triple-negative metastatic breast cancer^32^. Analysis of immunofluorescence in metastatic pulmonary lesions demonstrated a significant reduction in Ki-67⁺ proliferating cells in Cer2-OVA MMTV-PyMT lesions following VS-4718 treatment, suggesting that FAK inhibition suppresses tumor cell proliferation *in vivo* (**Fig. 1D&E**). As we did not observe an effect of VS-4718 treatment on the *in vitro* proliferation of breast cancer cells (**Supplementary Fig. 1D**), we conclude that the reduced *in vivo* proliferation observed in VS-4718-treated mice is due to microenvironment-mediated mechanisms.

**Figure 1:**
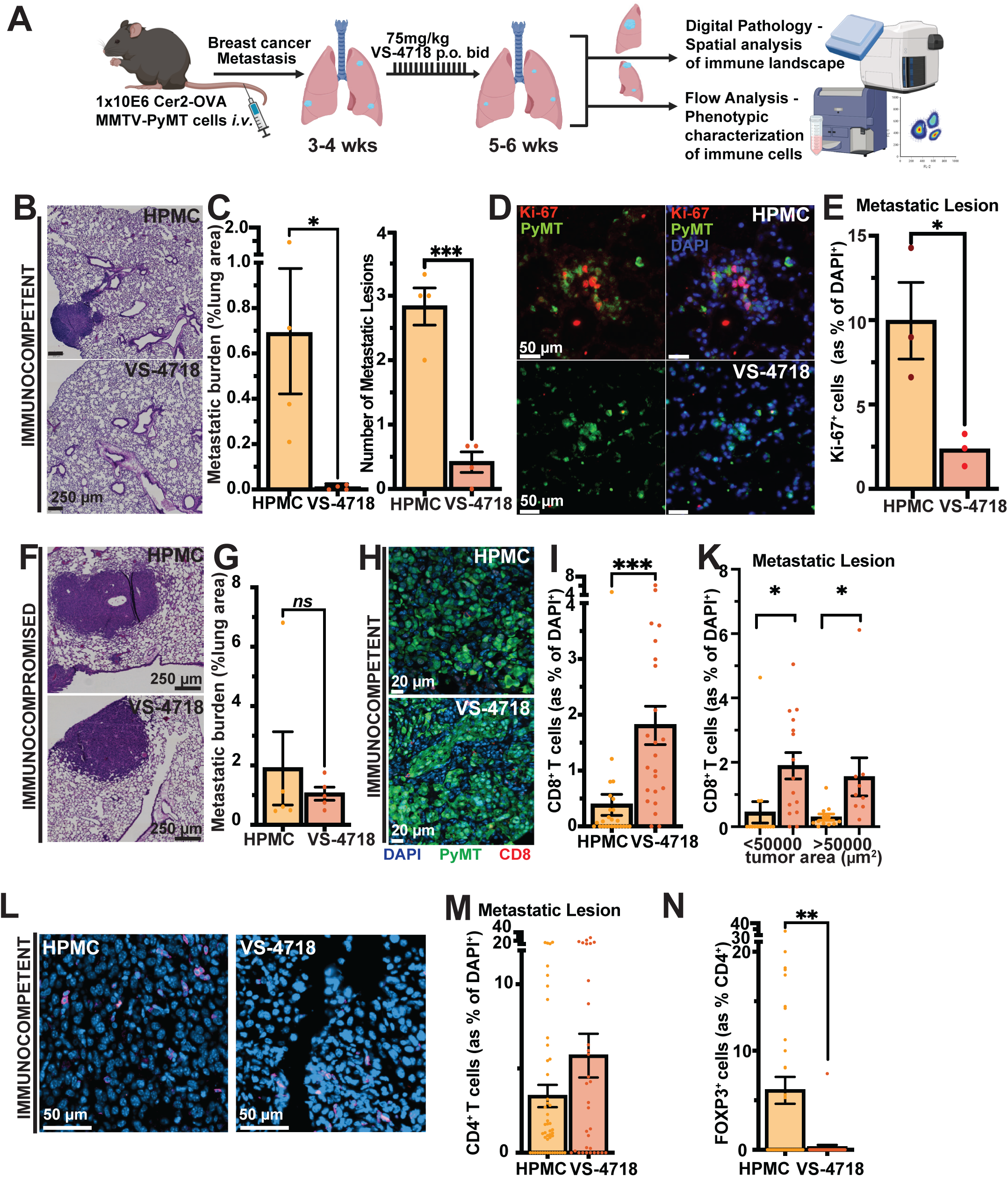
VS-4718 reduces pulmonary metastatic burden and reprograms T cell immunity. **(A)** Schematic of implantation of Cer2-OVA MMTV-PyMT cancer cells i.v. and their seeding into the lung. 3-4 weeks post implantation, mice are treated twice a day (bid) with either 0.5% HPMC (vehicle control) or 75 mg/kg VS-4718 for 2 weeks. At 5-6 weeks post implantation, lung tissue is collected for analysis. **(B)** Representative H&E images of lungs taken from control and VS-4718 treated mice at 5 weeks. Scale bar = 250 μm. **(C) Left:** Quantification of metastatic burden at 5 weeks post injection, expressed as metastatic area normalized to whole lung area. **Right:** Average number of metastases determined as the average of the number of metastatic lesions. n = 4 mice/group. (**D**) Representative images of Ki-67^+^ cells in metastatic lesions from vehicle and VS-4718 treated mice at 5 weeks post-injection. Ki-67 = red; Cer2-OVA MMTV-PyMT, anti-GFP = green; Nuclear staining, DAPI = blue. Scale bar = 50 μm. **(E)** Quantification of Ki-67^+^ cells. n = 4 mice/group. **(F)** Representative H&E images of lung metastatic burden in vehicle and VS-4718 treated CD-1 Nude mice at 5 weeks. **(G)** Quantification of lung metastatic burden in CD-1 Nude mice. n = 4 mice/group. **(H)** Representative images of CD8^+^ T cells in metastatic lesions in vehicle and VS-4718 treated mice at 6 weeks post injection. Nuclear staining, DAPI = blue); CD8^+^ T cells = red; Cer2-OVA MMTV-PyMT, anti-GFP = green. Scale bar = 20 μm. **(I)** Quantification of CD8^+^ T cells in metastatic lesions. Each dot represents CD8^+^ T cell as a percentage of all DAPI^+^ cells in metastatic lesions. n = 5 mice/group. **(K)** Percentage of CD8^+^ T cells relative to the size of metastatic lesion in mice treated with vehicle HPMC or VS-4718. Metastatic lesions were binned into two clusters according to their size <50000 or ≥50000 μm^2^. Each dot represents the percentage of CD8^+^ T cells within a lesion of the size. n = 5 mice/group. **(L)** Representative images of CD4^+^ T cells and regulatory T cells (FOXP3^+^) in metastatic lesions at 6 weeks post injection. Nuclear staining, DAPI = blue; CD4^+^ T cells = magenta; and FOXP3^+^ = red. Scale bar = 50 μm. **(M)** Quantification of CD4^+^ T cells. n = 5 mice/group. **(N)** Quantification of FOXP3^+^ regulatory T cells as a percentage of CD4^+^ T cells. Each dot represents FOXP3^+^ percentage of CD4^+^ T cells per metastasis. Mean ± SEM; * = p < 0.05, ** = p < 0.01, *** = p < 0.001. Mann-Whitney test was performed to compare groups in panels C, E, G, I, M, N.

VS-4718 treatment also led to increased infiltration of CD45⁺ leukocytes (CD8^+^ T cells in particular) in the lung, raising the possibility that FAK inhibition modulates the metastatic immune microenvironment (**Supplementary Fig. 3A-C, Supplementary Fig. 2F&G**). Interestingly, following VS-4718 treatment, we observed CD8⁺ T cells infiltrating into the core of the rare residual metastases, whereas in vehicle-treated lungs CD8⁺ T cells were largely confined to the periphery of lesions (**Supplementary Fig. 3D**). Given this observation, and the fact that FAK has previously been shown to modulate the immune environment (particularly T cells ^19^) in primary solid tumors, we next sought to investigate whether the reduction in metastatic burden following VS-4718 treatment is mediated by T cells. To test this, Cer2-OVA MMTV-PyMT cells were intravenously injected into T cell-deficient CD-1 Nude mice and metastatic burden assessed after VS-4718 treatment (**Fig. 1F**). VS-4718 treatment did not reduce metastatic burden in CD-1 Nude mice (**Fig. 1G**), indicating that the therapeutic effect of VS-4718 on metastasis is dependent on T cells. Interestingly, despite the absence of an effect on the overall metastatic burden, immunofluorescence analysis of Ki-67^+^ expression still revealed a reduction in metastatic proliferation following VS-4718 administration (**Supplementary Fig. 4**). This highlights that the VS-4718-mediated reduction in metastatic burden is T cell-dependent and is not due to direct anti-proliferative effects on tumor cells.

Because VS-4718 treatment nearly completely eradicated early metastatic lesions (**Fig. 1B&C**), immune cell infiltration within these lesions could not be reliably quantified. Therefore, to examine immune cell infiltration into the metastatic lesions in more detail, we started treatment at 4 weeks post-injection of Cer2-OVA MMTV-PyMT cells, which enabled us to characterize immune cells within the metastatic lesions in more detail. Similar to the early treatment regimen, delayed administration of VS-4718 led to a substantial regression of metastatic burden, and a significant reduction in the number of proliferating cells within metastatic lesions (**Supplementary Fig. 5A-D**). Interestingly, while we only observed a trend towards an increase in CD8⁺ T cell frequencies in tumor adjacent lung tissue (**Supplementary Fig. 5E**), CD8⁺ T cell infiltration within metastatic lesions following VS-4718 treatment was substantially increased (**Fig. 1H&I**). Stratification by size of lesions revealed that both small and large metastases in VS-4718-treated mice harbored a higher percentage of CD8⁺ T cells, indicating that FAK inhibition via VS-4718 promotes the infiltration of CD8^+^ T cells into metastatic lesions, regardless of size (**Fig. 1K**). In contrast to CD8⁺ T cells, no change in CD4⁺ T cells was observed within metastatic lesions or the adjacent lung tissue (**Fig. 1L&M, Supplementary Fig. 5F**). Meanwhile, we did observe a significant reduction in regulatory CD4⁺ T cells (Tregs, FOXP3^+^ CD4^+^ T cells) in metastatic lesions (**Fig. 1N**) and a similar trend in the tumor adjacent lung following VS-4718 treatment (**Supplementary Fig. 5G**). Since CD8⁺ T cell infiltration into tumors and metastatic lesions is a known critical predictor of response to therapy, whereas its absence is frequently associated with therapeutic resistance ^33,34^, these data indicate that FAK inhibition remodels the metastatic immune microenvironment toward a more anti-tumorigenic state. Furthermore, there is an increase of CD8^+^ T cells along with a concomitant decrease in Treg cells, and this change in CD8⁺ T cell and Treg distribution provides a more robust predictor of therapeutic outcome than either metric alone, with elevated ratios associated with better immune responses and survival in both pre-clinical and clinical studies ^35–37^.

### VS-4718 creates an anti-tumorigenic immune landscape in metastatic lungs

Given the effect of VS-4718 on T cells, we next investigated whether VS-4718 has the capacity to prime T cell immunity and prevent immune cell exhaustion by characterizing CD8⁺ T cell subpopulations in tumor bearing mice with multi-color flow cytometry. We defined CD8^+^ T cells as naïve (CD62L^hi^CD44^lo^), memory (CD62L^hi^CD44^hi^), or effector (CD62L^lo^CD44^hi^) cells (**Supplementary Fig. 6A/B**). While the proportions of memory and effector CD8⁺ T cells in the whole metastatic lung remained unchanged following VS-4718 treatment, we observed a significant increase in naïve CD62L^hi^CD44^lo^ CD8⁺ T cells (**Supplementary Fig. 6B**). The enrichment of naïve CD8⁺ T cells in tumor-bearing mice implies that VS-4718 may facilitate their recruitment or retention, thereby priming the host for anti-tumor responses. Supporting recruitment, we found an increase of CD8⁺ T cells in the peripheral blood of VS-4718-treated mice (**Supplementary Fig. 7**) ^33,34^. Upon encountering tumor antigen within metastatic lesions, these naïve cells may differentiate into effector CD8⁺ T cells, potentially contributing to the observed reduction in metastatic burden. We also found that VS-4718 treatment led to an increase in the naïve CD4⁺ T cell subpopulation, although this did not reach significance, and this was accompanied by a reduction in regulatory CD4⁺ T cells (**Supplementary Fig. 6C&D**). The effects on CD8^+^, CD4^+^, and regulatory CD4^+^ T cells in the lung and blood were not observed in the naïve (non-tumor-bearing) lung, indicating that they are dependent on the presence of metastatic lesions in the lung (**Supplementary Fig. 8**). These findings further reinforce the notion that VS-4718 remodels the immune landscape in metastatic lungs toward a more anti-tumorigenic state.

Further characterization of the CD8^+^ T cell subsets in tumor-bearing mice (**Fig. 2A**) showed that VS-4718 treatment markedly reduced the expression of the exhaustion markers PD-1 and CTLA-4 on CD8⁺ T cells, without affecting TIM-3. This suggests that VS-4718 causes a shift from an exhausted to a more functionally active T cell phenotype (**Fig. 2B**). Notably, prior studies have shown that TIM-3⁺ CD8⁺ T cells can maintain proliferative potential in the context of low PD-1 expression, thereby sustaining effective T cell responses ^38^. Reductions in specific exhausted subsets, including transitory exhausted (PD-1⁺EOMES⁺) and terminally exhausted (PD-1⁺CTLA-4⁺TIM-3⁺) CD8⁺ T cells, were also observed in tumor-bearing mice following VS-4718 treatment (**Fig. 2C-E**) and this was predominantly caused by the loss of a double-positive PD1^+^ CTLA4^+^ population (**Fig. 2D**).

**Figure 2:**
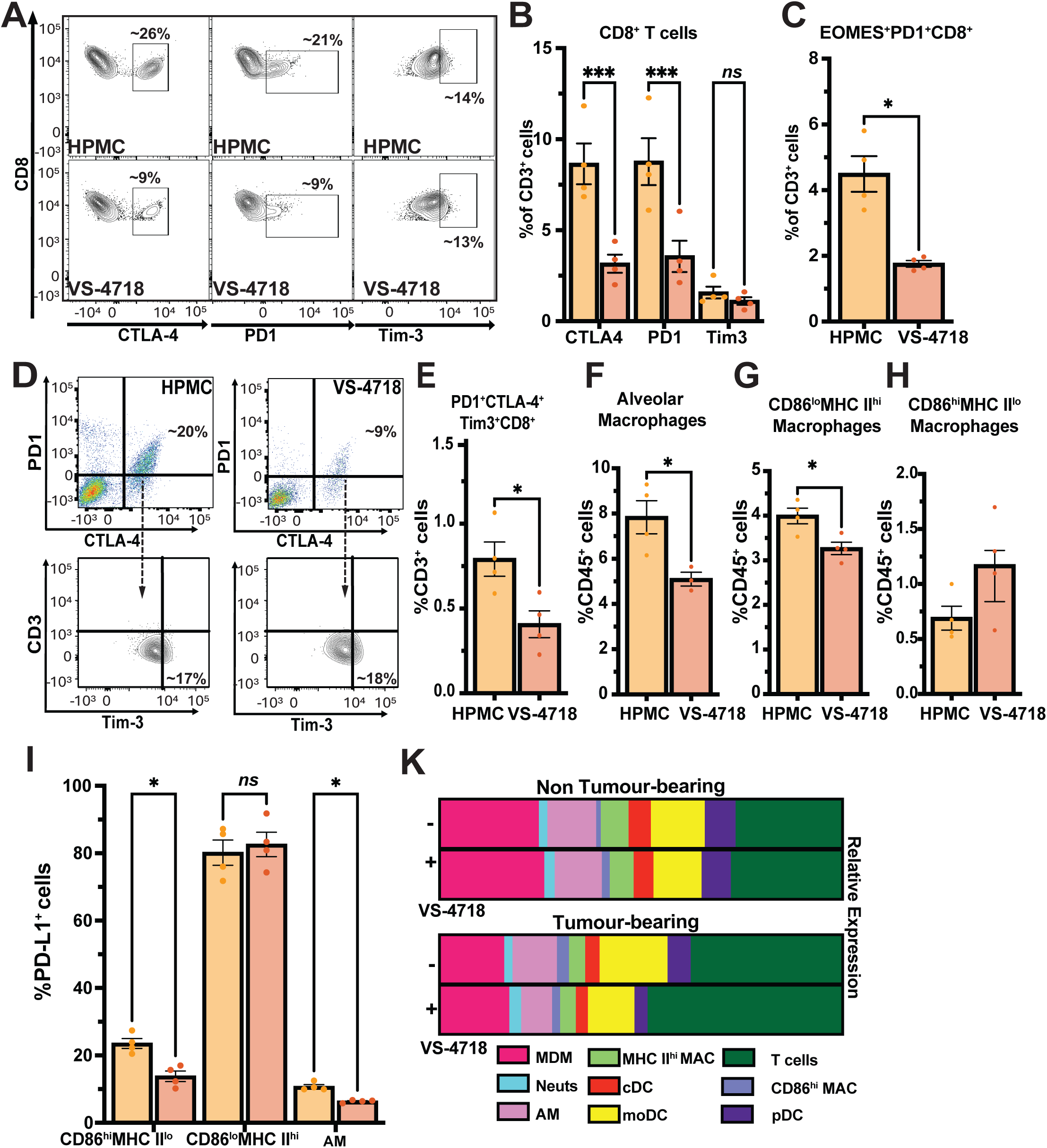
VS-4718 remodels the immune landscape and reduces T cell exhaustion in metastatic lungs. **(A)** Contour-plots of CTLA-4-, PD-1-, and Tim-3-surface expression on CD8^+^ T cells from Cer2-OVA MMTV-PyMT tumor-bearing mice treated with vehicle (0.5% HPMC, top panel) or VS-4718 (75 mg/kg, bottom panel) for two weeks. n = 4 mice/group. **(B)** Quantification of checkpoint-positive CD8^+^ T cells. Mean +/- SEM, *** = p < 0.001. **(C)** Frequency of PD-1^+^EOMES^+^CD8^+^ T cells (pre-exhausted subset) in lungs of vehicle- vs, VS-4718-treated mice. n = 4 mice/group. **(D)** Representative pseudo-color plots of PD-1 and CTLA-4 co-expression (**top**) and Tim-3 expression in PD-1^+^, CTLA-4^+^, CD8^+^ T cells (**bottom**). **(E)** Quantification of PD-1^+^CTLA-4^+^Tim-3^+^ CD8^+^ T cells (severely exhausted phenotype). n = 4 mice/group. **(F-H)** Frequency of alveolar macrophages (AM) in metastatic lung. AM (CD45^+^CD11b^+^CD11c^+^F4/80^+^CD169^+^); CD86^lo^ MHC II^hi^ macrophages (CD45^+^CD11b^+^CD11c^-^ F4/80^+^CD86^lo^MHC II^+^) and CD86^hi^MHC II^lo^ macrophages (CD45^+^CD11b^+^CD11c-F4/80^+^CD86^hi^MHC II^lo^). n = 4 mice/group. **(I)** VS-4718 reduces PD-L1 expression across myeloid subsets. Percentage of PD-L1^+^ cells in CD86^hi^ macrophages, MHC II^hi^ macrophages and AMs. n = 4 mice/group. **(K)** Stacked bar chart of myeloid and lymphoid populations in lungs of naïve and tumor-bearing mice treated with vehicle or VS-4718. Populations include neutrophils, monocyte derived macrophages (MDM), AMs, CD86^hi^ and MHC II^hi^ macrophages, classical DCs (cDCs), moDCs and plasmacytoid DC (pDC). Mean ± SEM; * = p < 0.05, ** = p < 0.01, *** = p < 0.001.

Given the predominance of myeloid cells within the lung, we next investigated whether FAK inhibition reshapes the pulmonary myeloid landscape in tumor-bearing mice. Flow cytometric profiling **(Supplementary Fig. 9A**) revealed no major alterations in myeloid subsets in non-tumor-bearing lungs following VS-4718 treatment (**Supplementary Fig. 9B**). However, in tumor-bearing lungs, VS-4718 treatment led to a significant reduction in alveolar macrophages (AMs) and CD86^lo^MHC II^hi^ but not CD86^hi^MHC II^lo^ macrophages, (**Fig. 2F-H**). These findings are consistent with prior studies reporting that AMs suppress CD8⁺ T cell responses and that their depletion can enhance CD8⁺ T cell infiltration ^39^, offering a potential mechanistic basis for the increased CD8⁺ T cell presence observed in metastatic lesions upon VS-4718 treatment. Notably, we also detected a decrease in PD-L1 expression among CD86^hi^ AMs (**Fig. 2I**), further supporting a reduction in immunosuppressive signaling within the tumor microenvironment. This diminished checkpoint expression among persisting myeloid populations is only observed in tumor bearing mice (**Fig. 2I & Supplementary Fig. 9C**) and may further relieve suppression of effector T cells and contribute to enhanced anti-tumor responses. Moreover, we observed a marked shift in the immune cell balance, with T cell populations increasing relative to myeloid populations in tumor-bearing mice receiving VS-4718 (**Fig. 2K**), indicating a reprogramming of the lung immune landscape toward T cell-mediated immunity. Interestingly, VS-4718 does noy impact myeloid and T cell populations in non-tumor bearing mice (**Fig. 2K**). The robust increase in CD8⁺ T cells, the reduction in regulatory CD4^+^ T cells, and the substantial regression of metastatic burden following FAK inhibition, suggest that the reshaping of the immune landscape is sufficient to elicit effective anti-tumor immunity.

Given previous reports that FAK can modulate cytokine release in tumor cells and thereby reshape the tumor immune microenvironment ^17,40^, we investigated whether this modulation could contribute to the immune cell changes observed in our models (**Supplementary Fig. 2G**). Because these effects appeared to be tumor-dependent, we hypothesized that tumor cell-intrinsic FAK signaling modulates cytokine secretion to influence immune composition. To test this, we performed a cytokine array analysis using conditioned media from VS-4718-treated Cer2-OVA MMTV-PyMT cells. Profiling 64 cytokines and chemokines commonly implicated in both pro- and anti-tumor immune responses revealed that VS-4718 selectively downregulated factors associated with recruitment of regulatory T cells and myeloid cells, including CCL2, CCL5, CXCL7, and LIX (**Supplementary Fig. 10**). These cytokines are known mediators of immunosuppressive infiltration and correlate with poor clinical outcomes, particularly in TNBC^41–43^. Such cytokine alterations likely contribute to the immune cell shifts observed by flow cytometry following VS-4718 treatment.

### VS-4718 enhances CD8⁺ T cell infiltration, directionality, and tumor engagement in vivo

Since we had not seen changes in cytokines linked to CD8^+^ T cell mobilization, but we did see an inability of the CD8^+^ T cells to infiltrate the core of metastatic lesions, we next sought to investigate whether VS-4718 influences CD8^+^ T cell migration dynamics *in vivo*. As PD-1 signaling is also known to promote a highly motile phenotype (by suppressing antigen-induced T cell arrest and reducing dwell time at sites of antigen presentation ^44^), we also examined the impact of combining VS-4718 with anti-PD-1 checkpoint blockade on CD8⁺ T cell behavior. To visualize CD8^+^ T cell behavior *in vivo*, we adoptively transferred CellTracker^TM^ Green-labelled OT-1 CD8^+^ T cells into mice bearing Cer2-OVA MMTV-PyMT lung metastases and implanted with a window for high-resolution time-lapse imaging of the lung (**Fig. 3A**). **Fig. 3B** shows representative 3D reconstructions of time-lapse imaging 3 h after injection of CD8^+^ T cells showing the majority of CD8^+^ T cells were associated with the vasculature and not engaged with Cer2-OVA MMTV-PyMT cells in vehicle treated mice (1^st^ row). Following treatment with VS-4718, more CD8^+^ T cells extravasated and interacted with Cer2-OVA MMTV-PyMT cells. Similar behaviors of CD8^+^ T cells could be seen upon addition of anti-PD-1 checkpoint inhibitor and in the combination treatment. Quantification of time-lapse imaging movies confirmed the increased CD8^+^ T cell extravasation following VS-4718 treatment, which was further enhanced by anti-PD-1 co-administration (**Supplementary Fig. 11**). Time-lapse intravital imaging revealed that all treatments caused decreases in the percentage of motile CD8⁺ T cells that had extravasated, though the decrease for VS-4718 treatment alone was not statistically significant (**Fig. 3C**). We also observed a marked decrease in CD8^+^ T cell velocity in treatments containing VS-4718, with combination treatment reducing T cell velocity by ∼85%, from 1.8 μm/min to 0.25 μm/min (**Fig. 3D**). Importantly, we did not observe a significant effect of VS-4718 treatment on CD8^+^ T cell velocity in the tumor-adjacent lung (**Supplementary Figure 12A**). Together these features are consistent with treatment causing increased tumor cell homing, antigen-driven arrest, and tumor engagement ^45,46^.

**Figure 3:**
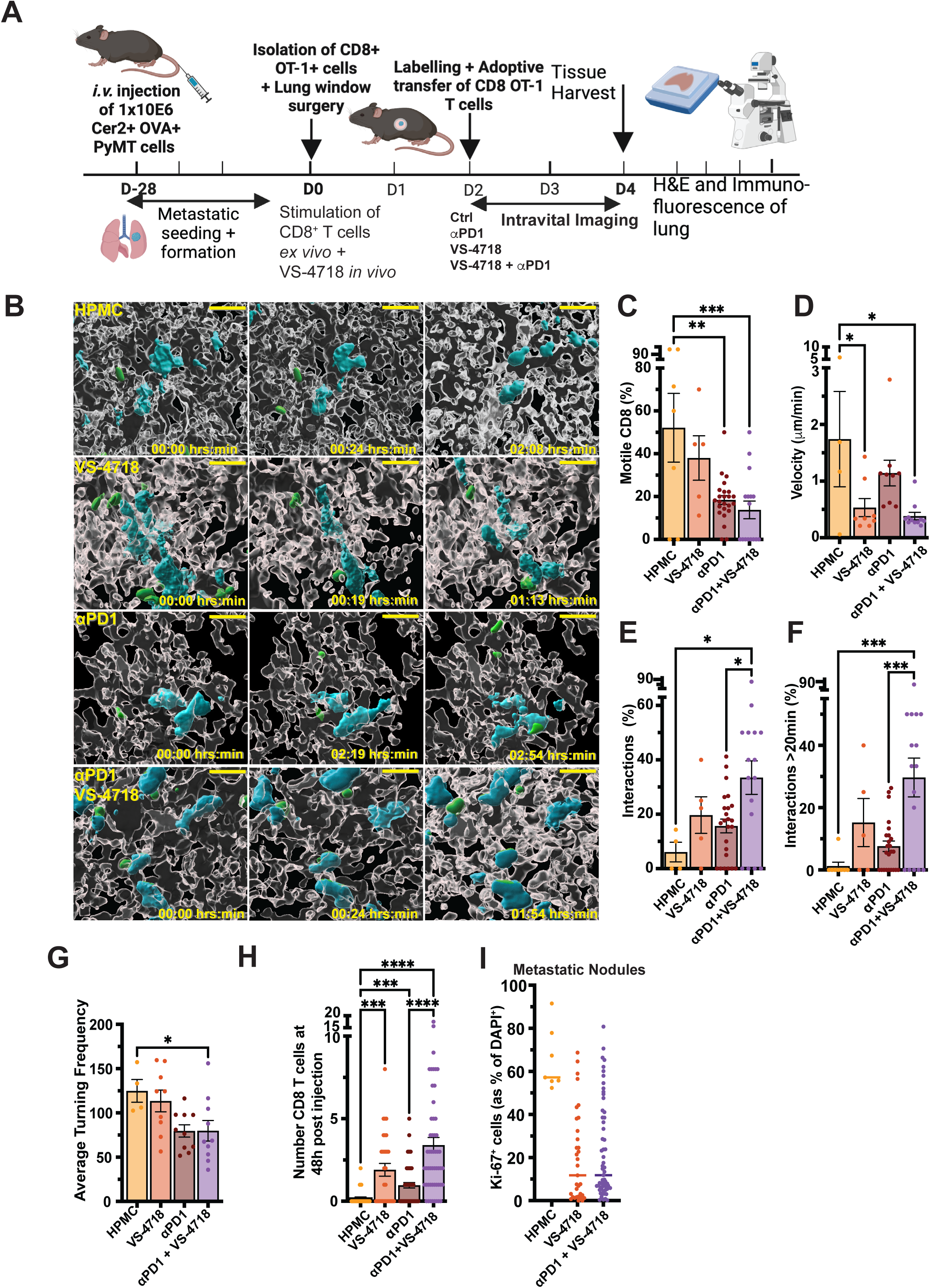
VS-4718 promotes CD8⁺ T cell infiltration and tumor cell engagement in pulmonary metastases. **(A)** Schematic of experimental setup. C57BL/6 mice were injected i.v. with 1x10^6^ Cer2-OVA MMTV-PyMT cells. Following lung window implantation, mice received CellTracker™ Green-labelled OT-1 CD8⁺ T cells i.v. 24 h post-surgery. Treatments included vehicle (0.5% HPMC), VS-4718 (75 mg/kg), anti-PD-1 (100 μg), or combination therapy, administered for 48 h prior to intravital imaging. **(B)** Representative 3D reconstructions of imaging data (3 h post-T cell transfer) showing OT-1 CD8⁺ T cells (green) interacting with Cer2-OVA MMTV-PyMT cells (cyan). Vasculature labelled with i.v. injected TMR-labelled 155 kDa dextran (white). Quantification of CD8⁺ T cell behavior from time-lapse imaging showing: **(C)** percentage of motile CD8^+^ T cells, **(D)** average velocity, **(E)** percent of all T cells interacting with tumor cells, **(F)** percent of all T cells with stable interactions (>20 min), and **(G)** average turning frequency. n ≥ 3 mice/group using 6-12 fields each. CD8⁺ T cell tracking performed with ROI Tracker plugin in Fiji. One-way ANOVA. **(H)** Number of CD8^+^ T cells 48 h post adoptive transfer. Dots represent intravital region of interests. n = 5 mice/group. **(I)** Quantification of proliferating tumor cells (Ki-67⁺) in metastatic nodules. Mean ± SEM; * = p < 0.05, ** = p < 0.01, *** = p < 0.001, **** = p < 0.0001.

To directly assess CD8⁺ T cell-tumor cell interactions, we quantified (a) the proportion of CD8⁺ T cells engaging with cancer cells including transient interactions (**Supplementary Movie 1**) and (b) the frequency of sustained, longitudinal interactions (**Supplementary Movie 2**). Quantification of CD8⁺ T cell-tumor cell contacts showed an increased proportion of CD8⁺ T cells engaging tumor cells in VS-4718- and combination-treated mice (**Fig. 3E)**. Notably, stable interactions (>20 min), which were absent in vehicle-treated animals, were observed for 18% and 30% of CD8^+^ T cells in the VS-4718 and combination groups, respectively (**Fig. 3F**). Increased stable interactions is indicative of effective immunological synapse formation and cytotoxic potential ^47,48^. Moreover, combination treatment facilitated frequent and multiple interactions between CD8⁺ T cells and individual tumor cells, a hallmark of effective cytotoxic killing ^46,47^. Prior studies have shown that 2-9 serial CD8⁺ T cell-tumor interactions are sufficient to induce tumor cell apoptosis, with as few as three distinct contacts within 50 min driving cumulative cytotoxicity and irreversible tumor cell death ^47^. Turning frequency, a metric of directional migration was decreased significantly under combination therapy, but not other treatments (**Fig. 3G**) and was not modulated in the tumor-adjacent lung (**Supplementary Fig. 12B**). A lower average turning frequency reflects increased directional persistence, indicating cells are maintaining their trajectory toward a target rather than executing frequent directional shifts ^49,50^. Notably, the combination treatment also significantly increased persistence of CD8^+^ T cells in imaged sections, as shown by the quantification of CD8^+^ T cell numbers upon re-localization of the same field of view 48 h after the first imaging session (**Fig. 3H**).

To evaluate the functional impact of these interactions, we assessed tumor cell proliferation 96 h after treatment. Proliferation was significantly reduced in GFP⁺ metastatic lesions in VS-4718- and combination-treated mice (**Fig. 3I**). This indicates that proliferation is modulated, even upon short treatment with VS-4718 or combination therapy *in vivo*. These effects are in accord with the previously reported increase of CD8^+^ T cells in the metastatic lungs (**Fig. 1H**). Consequently, we hypothesized that both VS-4718 and anti-PD-1 increase the extravasation rate of CD8^+^ T cells, and that this can be further increased with combination therapy. VS-4718 treatment alone is sufficient to decrease CD8^+^ T cell velocity, potentially linked to an increased antigen-driven arrest response. While VS-4718 treatment alone does not directly lead to an increase in CD8^+^ T cell and cancer cell interactions, it can significantly enhance anti-PD-1 driven CD8^+^ T cell interactions (**Fig. 3E&F**). Finally, anti-PD-1 is sufficient to modulate the CD8^+^ T cell turning frequency. Taken together, these findings highlight a synergistic mechanism wherein VS-4718 and anti-PD-1 cause greater CD8⁺ T cell access and retention.

### VS-4718 Enhances CD8⁺ T Cell-Tumor Interactions and Cell Killing in Lung Metastases ex vivo

Given that intravital imaging revealed enhanced CD8^+^ T cell extravasation and tumor engagement following VS-4718 and anti-PD-1 treatment, we sought to validate these effects by extending observations and dissecting the underlying mechanisms. However, because assessing T cell-mediated killing *in vivo* using intravital imaging is technically challenging, we established an *ex vivo* model to study CD8⁺ T cell-mediated cytotoxicity in lung metastases. Thus, we next examined whether FAK inhibition affects CD8⁺ T cell dynamics and cytotoxicity in *ex vivo* metastatic lung slice co-cultures that preserve the structure and matrix context of metastatic foci.

Harvested lung slices were pretreated with vehicle, anti-PD-1, VS-4718, or their combination, followed by addition of OT-1 CD8⁺ T cells (**Fig. 4A**). Within the first 3 h of co-culture, the number of interactions (but not stable interactions) were altered, comparable to *in vivo* imaging measurements across treatment groups (**Supplementary Fig. 13A&B**). However, combination therapy did increase T cell turning frequency at this timepoint (**Supplementary Fig. 13C**), though the effect was not sustained at 24 h (**Supplementary Fig. 13D**). By 24 h, the percentage of motile CD8^+^ T cells increased following VS-4718 and combination treatment (**Fig. 4B**). Consistent with our *in vivo* measurements (**Fig. 3D**), VS-4718 alone or in combination reduced CD8⁺ T cell velocity (**Fig. 4C**). At this time point, CD8⁺ T cell-tumor interactions increased markedly in combination-treated slices (**Fig. 4D&E**), recapitulating *in vivo* observations (**Fig. 3E&F**).

**Figure 4:**
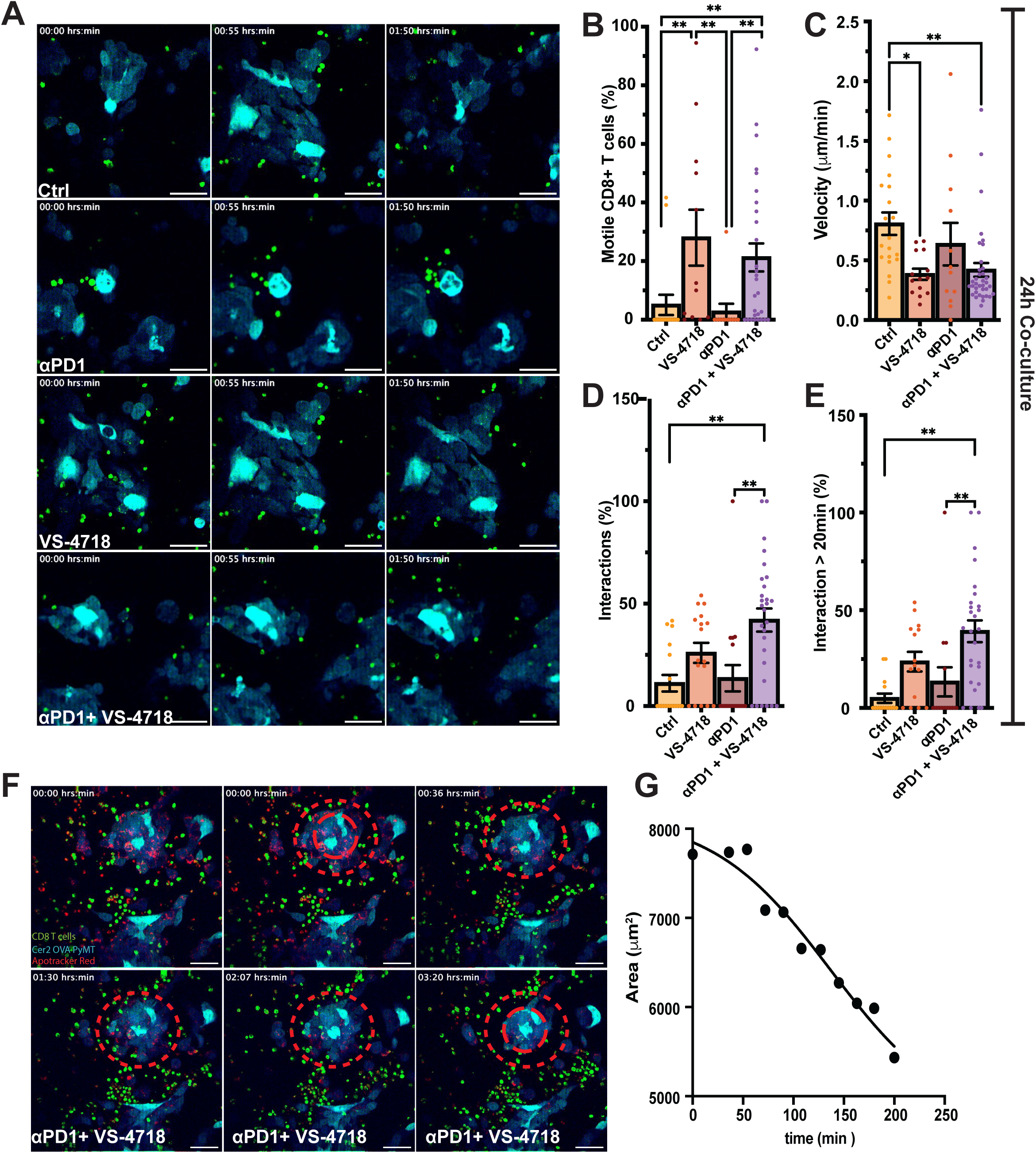
VS-4718 enhances CD8⁺ T cell-tumor cell interactions and induces cancer cell death *ex vivo*. **(A)** Representative images of CD8⁺ T cells (green) interacting with tumor cells (cyan) in lung slices cultured *ex vivo*. Scale bar = 50 μm. Quantification of CD8⁺ T cell-tumor cell dynamics after 24 hrs of treatment, showing: **(B)** percentage of motile CD8^+^ T cells, **(C)** velocity, **(D)** interaction frequency, and **(E)** stable interactions (>20 min). n ≥ 3 mice/group using ≥ 3 slices each. **(F)** Representative images of metastatic lesion area over time movie under combined treatment. Dashed circle = size of metastatic lesion at t=0. Solid circle = size of metastatic lesion at t=3:20. **(G)** Quantification of tumor lesion area over time. Mean ± SEM; * = p < 0.05, ** = p < 0.01, *** = p < 0.001, **** = p < 0.0001.

To assess whether enhanced interactions reflected increases in CD8⁺ T cell activity, we measured T cell size which has been shown to be a surrogate for activation ^51,52^. CD8⁺ T cells cultured alone or with IL-2 contracted over time, while full stimulation with anti-CD3/CD28 and IL-2 robustly increased size (**Supplementary Fig. 14A&B**). CD8⁺ T cells on metastatic lung slices showed modest (though insignificant) enlargement, consistent with tonic activation. Only the combination of VS-4718 and anti-PD-1 significantly increased T cell size beyond IL-2 alone, albeit below levels induced by full stimulation with anti-CD3/CD28 and IL-2.

We next looked at the cytotoxic potential of CD8^+^ T cells by imaging their behavior and detected CD8^+^ T cell swarming around tumor cells (**Supplementary Movie 3**), which was associated with progressive shrinkage of metastatic lesions only with the combination treatment (**Fig. 4F&G**). Furthermore, we observed that combination treatment led to widespread apoptosis across lesions, as detected by Apotracker Red ^53^ staining at day 7 (**Supplementary Fig. 14C**).

Collectively, these findings indicate that combination therapy enhances CD8⁺ T cell-tumor cell interactions and cytotoxicity. Further, VS-4718 had no direct effect on activation of CD8⁺ T cells (**Supplementary Fig. 15**), indicating that increased killing in co-cultures is not driven by direct of FAK inhibition on CD8^+^ T cells.

### VS-4718 suppresses ECM production

As FAK inhibition did not directly impact CD8⁺ T cells but influenced their behavior in tumor-bearing lungs, we reasoned that VS-4718 may be acting indirectly through tumor cell-driven remodeling of the ECM. FAK signaling is known to regulate ECM secretion, organization, and turnover in cancer cells. Thus, its inhibition could remodel the tumor ECM and, in turn, influence CD8⁺ T-cell migration and engagement. We therefore examined whether VS-4718 alters ECM composition and associated signaling pathways in tumor cells through proteomic analysis. Cer2-OVA MMTV-PyMT cells were treated with VS-4718 at a concentration that inhibited FAK activation (**Supplementary Fig. 1C**) and cells and supernatant were harvested to determine changes in the cellular proteome and secretome. Volcano plot analysis of differentially expressed proteins in the intracellular and secreted fractions revealed a broad restructuring of the tumor cell proteome and secretome (**Fig. 5A&B**). Unsupervised hierarchical clustering of log₂-transformed, median-centered, z-scored proteomic data identified three discrete clusters (C1-C3) with distinct expression dynamics (**Fig. 5C, Supplementary Table 1**). Proteins in Cluster C1 were depleted intracellularly but enriched in the secretome upon VS-4718 treatment, suggestive of increased secretion. Cluster C2 comprised proteins upregulated both within the cells and in the supernatant, while Cluster C3 contained proteins that were consistently downregulated across both compartments.

**Figure 5:**
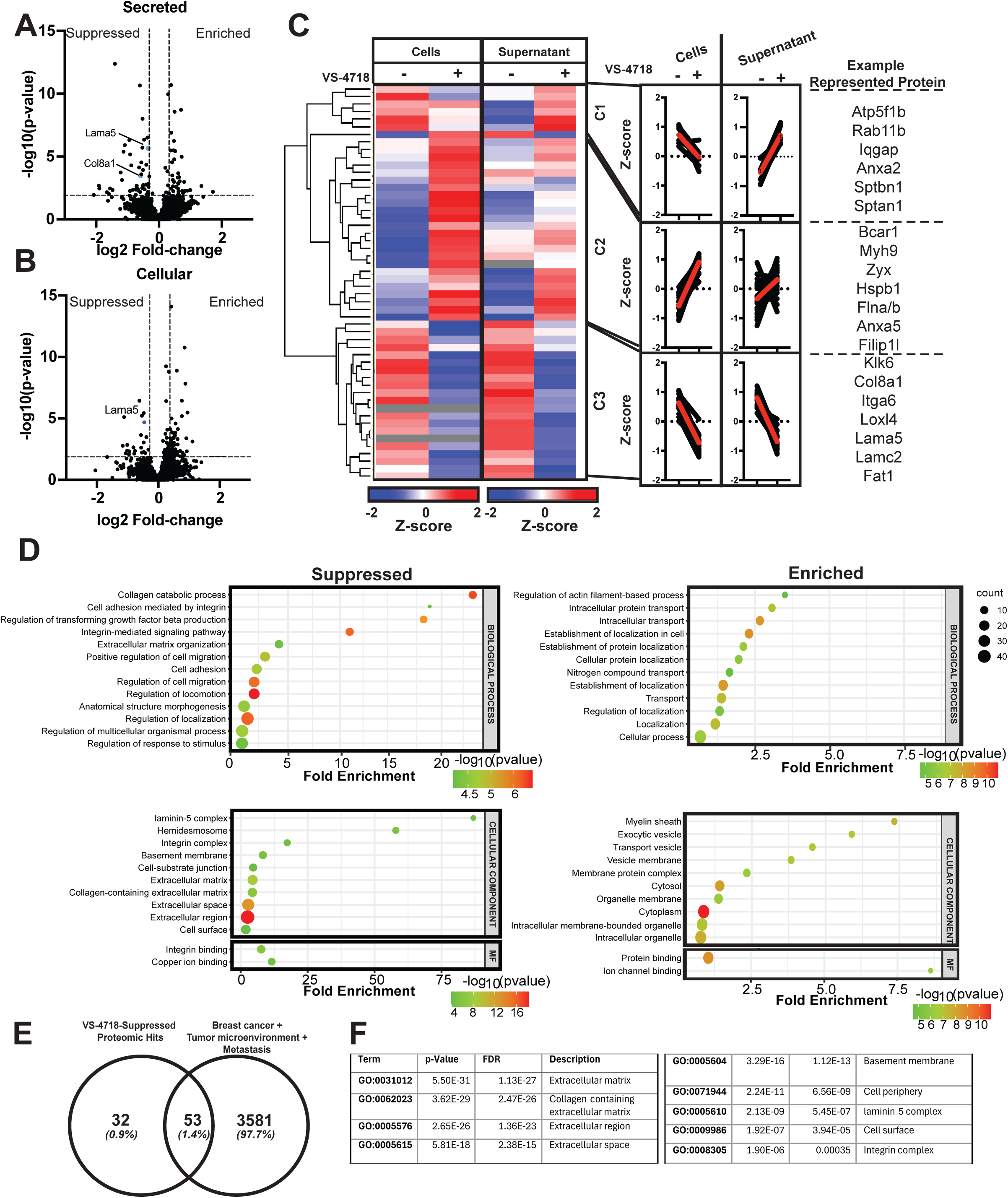
VS-4718 reprograms the tumor-intrinsic proteome to suppress ECM. **(A)** Volcano plot of differentially secreted and (**B**) cellular proteins following VS-4718 treatment. Multi-sample t-test, FDR-corrected. n = 3 independent experiments. Dashed lines in (A+B) indicate proteins considered as significantly suppressed ( top left) and enriched ( top right). **(C) Left:** Unsupervised hierarchical clustering of cellular and secreted proteins from Cer2-OVA MMTV-PyMT cells treated +/- VS-4718 for 24 h and 48 h, respectively, revealing three major clusters (C1-C3) based on Euclidean distance. **Right:** Cluster profiles and representative proteins from each cluster. Heatmap shows significantly changed proteins only. n = 3 independent experiments **(D)** Bubble plot representation of suppressed and enriched biological process (**top**) and cellular component (**bottom**) pathways in the secretome of Cer2 OVA PyMT cells treated for 48 h with VS-4718. MF = molecular functions. **(E)** Venn-Diagram of all significantly downregulated targets in secretome of cells upon VS-4718 treatment and a list of genes identified in public databases to be involved in breast cancer, tumor microenvironment and metastasis. **(F)** Gene ontology of common genes between proteomic hits and public database genes.

Gene ontology (GO) enrichment of proteins significantly modulated in both the cellular and secreted proteome indicated a pronounced suppression of biological processes associated with ECM organization, cell adhesion, migration, and integrin-mediated signaling (**Fig. 5D**). Consistently, downregulated components included hemidesmosomes, laminin-5 complexes, basement membrane constituents, and broader ECM-related structures, while the predominant molecular function suppressed was integrin binding. The main biological processes enriched predominantly include transportation pathways, hence an enrichment of vesicles and membrane components can also be observed (**Fig. 5D** bottom right). The most enriched molecular functions following FAK inhibition were associated with protein and ion binding (**Fig. 5D** bottom right), indicative of broader remodeling of trafficking and intracellular communication networks. We focused our further analysis on proteins significantly modulated in the secretome and STRING-based protein-protein interaction analysis (**Supplementary Fig. 16**) showed that suppressed protein networks were strongly associated with ECM-receptor interactions (e.g., laminin α5, laminin γ5, laminin β3, integrin α6, integrin β4), collagen fibril assembly (e.g., collagen VIII α1), matrix metalloproteinase activation and ECM degradation (e.g., MMP13, MMP10, MMP3), and components of the complement and coagulation cascades (e.g., C3, CFH). Moreover, we also see a significant reduction in basement membrane components highlighted in pale yellow (e.g., laminin α3, laminin β3). As most upregulated factors were associated with the phenotype of the cancer cells and not factors that would potentially impact the microenvironment, we decided to focus on the suppressed factors.

To assess whether VS-4718-regulated proteins identified in our *in vitro* proteomic analyses were also implicated in metastatic progression or immune modulation, we interrogated publicly available datasets from GeneCards, Comparative Toxigenomics Database (CTD), and the Human Genetics Knowledgebase. This approach yielded 3,634 genes associated with biological processes including, breast cancer, metastasis, and tumor microenvironment. Following conversion of our suppressed secreted proteomic hits to corresponding gene symbols and cross-referencing them with the metastasis-associated dataset, 53 overlapping genes were identified (**Fig. 5E, Supplementary Table 2**). Notably, gene ontology analysis of these shared genes showed an enrichment for ECM components, in particular collagen-containing ECM and basement membrane proteins including laminins, collagens, and integrins (**Fig. 5F**). This highlights that major changes mediated by VS-4718 on tumor cells impact the ECM.

### Changes in Matrix Components alter the barrier for CD8^+^ T cells and activity levels

To further investigate the impact of VS-4718 on the ECM, we chose to focus on laminin α5 (gene: *LAMA5*) and collagen VIII α1 (gene: *COL8A1*), as both of these proteins have been previously described as core components of the extracellular matrix in metastatic lesions, including pulmonary breast cancer metastases ^54^. Notably, collagen VIII α1 (a basement membrane collagen upregulated in breast^55^ and gastric cancer^56^), has not only been linked to poor survival, but high levels of *COL8A1* have also been linked to immune modulation and is associated with increases in T reg and macrophage infiltration^56^. Given this connection, and our observation that collagen VIII α1 and laminin α5 are downregulated *in vitro* following VS-4718 treatment, we hypothesized that FAK inhibition can alter the ECM metastatic niche to promote CD8^+^ T cell access and migratory behavior.

We first assessed whether VS-4718 reduced collagen VIIIα1 and laminin α5 expression in pulmonary metastatic breast cancer lesions in the Cer2-OVA MMTV-PyMT experimental metastasis model. Both proteins were markedly enriched in metastatic lesions relative to tumor adjacent lung tissue (**Fig. 6A&B**). VS-4718 treatment resulted in a pronounced reduction of collagen VIIIα1 and laminin α5 within metastatic lesions, while expression in tumor adjacent regions remained largely unaffected (**Fig. 6A&B, Supplementary Fig. 17**). Notably, expression of both proteins in the adjacent lung tissue was comparable with that in lungs from non-tumor bearing mice (**Supplementary Fig. 17**), indicating that VS-4718 may prevent their increased expression in metastatic lesions. We observed a similar increase of collagen VIIIα1 and laminin α5 in metastatic lesions compared to the tumor adjacent regions in the 4T1 model, which was again decreased upon treatment with VS-4718 (**Supplementary Fig. 18**).

**Figure 6:**
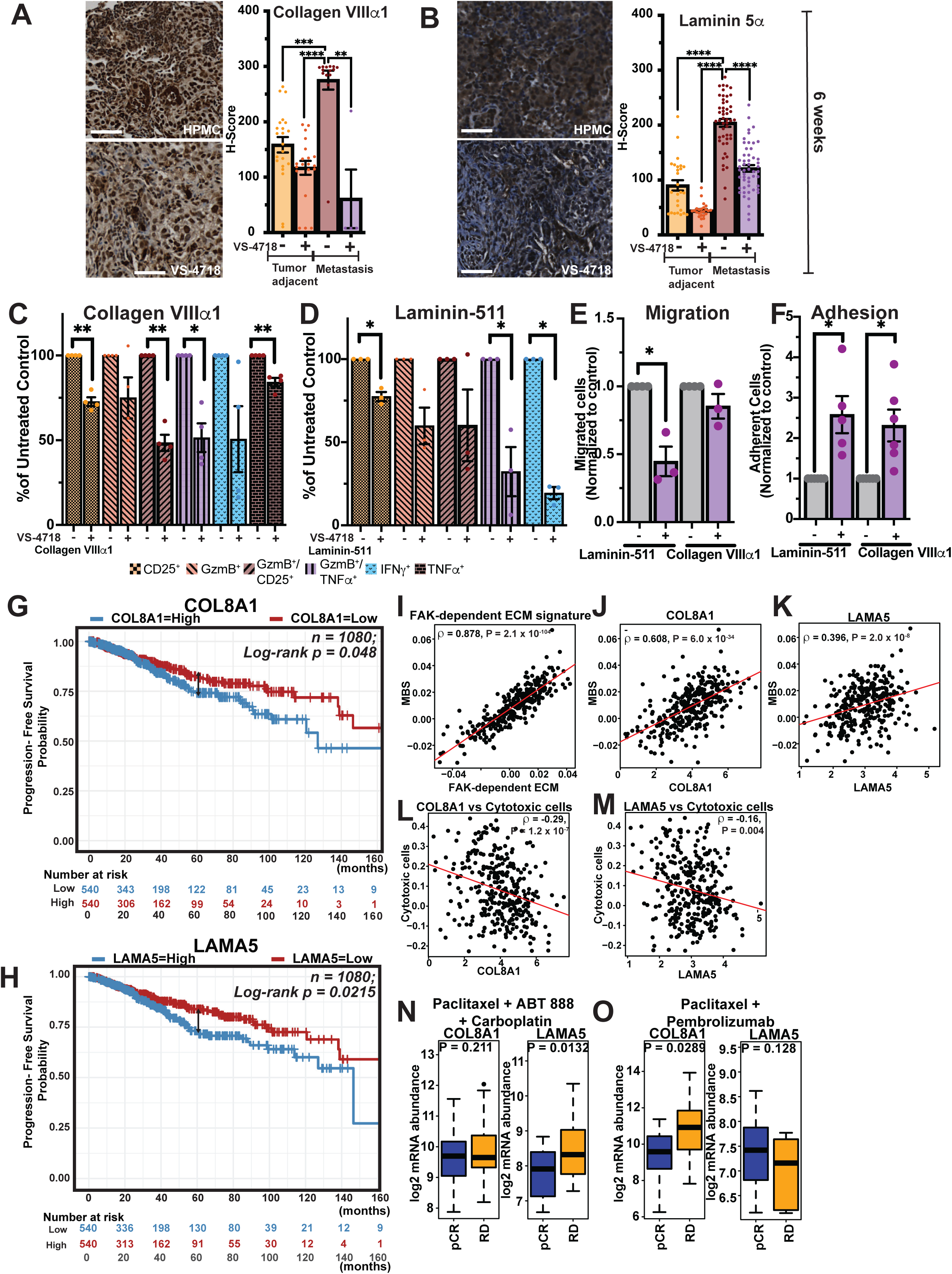
Collagen VIIIα and Laminin 5α restrict CD8⁺ T cell activation and correlate with survival in patients. Mice bearing Cer2-OVA MMTV-PyMT lung metastases were treated with vehicle (0.5% HPMC) or 75 mg/kg VS-4718 *p.o.* for 2 weeks starting 4 weeks post-implantation. (**A & B**) Representative images and quantification of collagen VIIIα1 and laminin 5α in metastatic versus non-metastatic lung regions at 6 weeks (H-score; each dot represents one lesion; n = 5 mice/group). Kruskal-Wallis test and Dunn’s multiple comparison. (**C & D**) Collagen VIIIα1 and laminin 511 suppress CD8⁺ T cell activation *in vitro*. CD8⁺ T cells were plated on 1 μg/mL ECM components, stimulated with CD3/CD28 + IL-2 for 48 h, and assessed by flow cytometry for CD25, granzyme B, IFNγ, and TNFα expression (n = 3-5). (**E**) ECM-mediated inhibition of CD8⁺ T cell migration. Transwell inserts coated with laminin 511, or collagen VIIIα were used to assess migration toward serum-rich media for 5 h (n = 3-4). (**F**) ECM-dependent CD8⁺ T cell adhesion. Adhesion to laminin 511, or collagen VIIIα-coated plates was quantified after 90 min and normalized to uncoated plates (n = 3-4). **(G & H)** Kaplan-Meier curves showing progression-free survival (PFS) of patients with invasive breast carcinoma stratified by **(G)** *COL8A1* and **(H)** *LAMA5* expression levels. Patients were divided into high and low expression groups based on the median RNA-seq expression. Point-wise z-test using Greenwood’s standard error. Data source = Breast invasive carcinoma TCGA. (**I-K**) Spearman’s Correlation of Metastatic burden signature correlated to FAK-dependent ECM score **(I**), COL8A1 (**J**) and LAMA5 (**K**) expression in triple negative breast cancer patients of the SCAN-B dataset^62^. Spearman’s correlation of cytotoxic CD8^+^ T cells with COL8A1 (**L**) and LAMA5 (**M**) expression in triple negative breast cancer patients of the SCAN-B dataset. (**L & M**). Log_2_ mRNA abundance of COL8A1 and LAMA5 in triple negative breast cancer patients with complete response (pCR) or residual disease (RD) in biopsies taken before treatment with either Palclitaxel and Pembrolizumab (**O**) or Palclitaxel, ABT888 and Carboplatin (**N**) in ISPY-2 trial.

Given prior evidence that basement-membrane components can modulate immune cell access and dampen lymphocyte activation ^57,58^, we next examined the effects of collagen VIIIα1 and laminin-511 on CD8⁺ T-cell activation *in vitro*. Laminin α5 is a key subunit of the laminin-511 complex which is commonly used to study laminin α5 effects ^59^. Splenic CD8⁺ T cells were activated on plates coated with or without either protein. Collagen VIIIα1 markedly reduced the proportion of TNFα⁺, CD25⁺ and double-positive CD25⁺/Granzyme B⁺ and Granzyme B⁺/TNFα⁺ CD8⁺ T cells. Similar trends were also observed with laminin-511 (**Fig. 6C&D**). These findings indicate that collagen VIII α1 and laminin-511 exert an inhibitory effect on CD8⁺ T-cell activation. Importantly, our finding that treatment of CD8^+^ T cells *in vitro* with VS-4718 does not directly change their activation (**Supplementary Fig. 15**) indicates that VS-4718’s immunostimulatory effects are not due to direct T cell stimulation. Instead, these results indicate that VS-4718 diminishes immunosuppression, in part, through remodeling of the tumor ECM, which reduces inhibitory ECM-immune interactions, and preserves CD8⁺ T-cell activity.

Because basement membrane components can also modulate immune cell trafficking and intratissue migration patterns ^60,61^, we next assessed their effects on CD8⁺ T-cell migration using a transwell assay. Both collagen VIIIα1 and laminin-511 suppressed CD8⁺ T-cell migration toward serum-rich medium; however, laminin-511 exhibited the most potent inhibitory effect (**Fig. 6E**). Decreased CD8⁺ T cell transwell migration might be caused by either physical restriction or increased adhesion of CD8⁺ T cells to basement membrane components. To explore the mechanism underlying ECM-mediated inhibition of CD8⁺ T-cell migration, we assessed CD8⁺ T-cell adhesion to laminin-511 and collagen VIIIα1 (**Fig. 6F**). Adhesion to both laminin-511 and collagen VIIIα1 was increased, suggesting that reduced transwell migration may result from enhanced tethering of CD8⁺ T cells to basement membrane proteins (**Fig. 6F**). In a random migration assay, we found that while CD8^+^ T cells plated on laminin-511 exhibit both reduced velocity and increased average turning frequency, collagen VIIIα1 had no effect (**Supplementary Fig. 19**). This may explain how despite similar effects on adhesion, collagen VIIIα1 does not change transwell migration, whereas laminin-511 coated transwells reduce migration. Together, these findings indicate that VS-4718 remodels the metastatic ECM, reducing immunosuppressive barriers, and enhancing CD8⁺ T-cell activation and migration.

We next investigated the clinical relevance of our findings to human breast cancer by using RNA-seq and clinical data from TCGA Breast Invasive Carcinoma cohort. Patients were stratified into “High” and “Low” expression groups based on the median expression value for each gene. As expected, elevated expression of either *COL8A1* and *LAMA5* was significantly associated with shorter progression free survival, consistent with their proposed pro-metastatic and immunosuppressive functions (**Fig. 6G&H**). Progression free survival probability at 5 years was significantly reduced from 0.82 to 0.74 for the “High” *COL8A1* group (p=0.047), and from 0.84 to 0.72 for the “High” *LAMA5* group (p = 0.0017). Further analysis using the more contemporary treated SCAN-B breast cancer dataset ^62^ showed that these matrix components are also associated with the GENIUS score, a genomic risk classifier that integrates subtype-specific multigene expression signatures, weighted by intrinsic subtype probabilities, into a single prognostic index of breast cancer recurrence risk, as well as with tumor grade and metastatic burden signatures in breast cancer patients. While only *PTK2* (FAK), but not *COL8A1* or *LAMA5*, showed a positive correlation with tumor grade across all breast cancer patients (**Supplementary Fig. 20A**), all genes showed a positive correlation with grade in patients with triple-negative breast cancer (TNBC) (**Supplementary Fig. 20B**) but not ER^+^ patients (**Supplementary Fig. 20C**). Further, we observe a positive correlation of GENIUS score with *PTK2*, the here identified FAK-dependent ECM signature, *COL8A1* in TNBC patients but not in all or ER^+^ breast cancer patients (**Supplementary Fig. 21**). Focusing on matrix-associated changes, we observed strong correlations between proteomic-identified FAK-dependent ECM components, including *COL8A1* and *LAMA5*, and metastatic burden signatures in TNBC (**Fig. 6 I-K**) compared with all breast cancer patients (**Supplementary Fig. 22A**). Furthermore, analysis of a dataset from Perou and colleagues comprising patient-matched primary and metastatic lesions ^63,64^, revealed that *COL8A1* expression remained largely unchanged, whereas *LAMA5* was significantly enriched in metastatic sites (**Supplementary Fig. 22B**). When analyzed by individual metastatic site (**Supplementary Fig. 22C-E**), this comparison held in each location except for lymph nodes, which was insignificant for *LAMA5*. These results suggest that laminin α5 accumulation is a feature of non-lymphoid metastatic niches.

As we observed a correlation between ECM component expression and CD8⁺ T-cell infiltration in our preclinical studies, we next examined whether ECM components, including *COL8A1* and *LAMA5*, were also associated with cytotoxic CD8⁺ T cells in the SCAN-B dataset (**Fig. 6L & M**). While no significant correlation was observed between cytotoxic CD8⁺ T cells and the FAK-dependent ECM score (**Supplementary Fig. 23A**), *COL8A1* and *LAMA5* expression showed a significant negative correlation with cytotoxic CD8⁺ T cells (**Fig. 6L & M**). These findings are consistent with our *in vitro* and *in vivo* data and suggest distinct mechanisms by which collagen VIII α1 and laminin α5 can influence immune regulation. When patients were stratified based on FAK-dependent ECM expression and cytotoxic CD8⁺ T-cell levels, overall survival was reduced in patients with high expression of *COL8A1*, *LAMA5*, or the FAK-dependent ECM score and low CD8⁺ T-cell expression, compared with patients exhibiting low ECM expression and high CD8⁺ T-cell levels (**Supplementary Fig. 23B–D**). However, survival was also improved in patients with high ECM expression and high CD8⁺ T-cell levels, indicating that outcomes are primarily driven by CD8⁺ T-cell abundance. Notably, patient data revealed a negative correlation between cytotoxic T cells and *COL8A1* expression, consistent with the immunosuppressive effects observed experimentally.

Finally, we investigated whether ECM remodeling might also influence therapeutic response. Analysis of a follow-up dataset from Perou and colleagues, in which lesions were classified as responders or non-responders ^65^, revealed a trend toward higher ECM scores in non-responding lesions (**Supplementary Fig. 23E-F**) compared to responding lesions (3/5 patients). These patients had received multiple prior chemotherapies, including carboplatin, gemcitabine, cyclophosphamide, and doxorubicin, yet non-responding lesions exhibited persistent ECM enrichment, supporting a link between matrix remodeling and therapy resistance. To assess the predictive value of the ECM score*, COL8A1*, and *LAMA5* for therapeutic response, we compared pre-treatment expression levels in TNBC patients achieving a complete response versus those with residual disease in the I-SPY 2 trial (**Fig. 6N&O**). Elevated *COL8A1* expression was observed in patients with residual disease that had been treated with paclitaxel and pembrolizumab, whereas elevated *LAMA5* expression was associated with residual disease in patients treated with paclitaxel, ABT-888, and carboplatin. These findings suggest that ECM components may serve as predictive markers of therapeutic response and support the potential benefit of incorporating FAK inhibition into therapy.

Together, these data indicate that elevated expression of ECM components such as *LAMA5* and *COL8A1* are associated with poor clinical outcome and metastatic progression. While the number of metastatic lesions with documented therapeutic response was limited, we observed a consistent trend toward higher ECM scores in non-responders, implying that a dense, laminin- and collagen-rich matrix may serve as a barrier to effective treatment. Integrating these clinical observations with our experimental findings suggests that FAK inhibition, by attenuating ECM deposition, may not only facilitate CD8⁺ T-cell access and function but also enhance the efficacy of standard and immune-based therapies in metastatic breast cancer. **Figure 7** provides a conceptual summary of this model, illustrating how FAK inhibition remodels the laminin- and collagen-rich metastatic ECM to enhance CD8⁺ T-cell access, activation, and tumor-cell clearance within pulmonary metastases.

**Figure 7:**
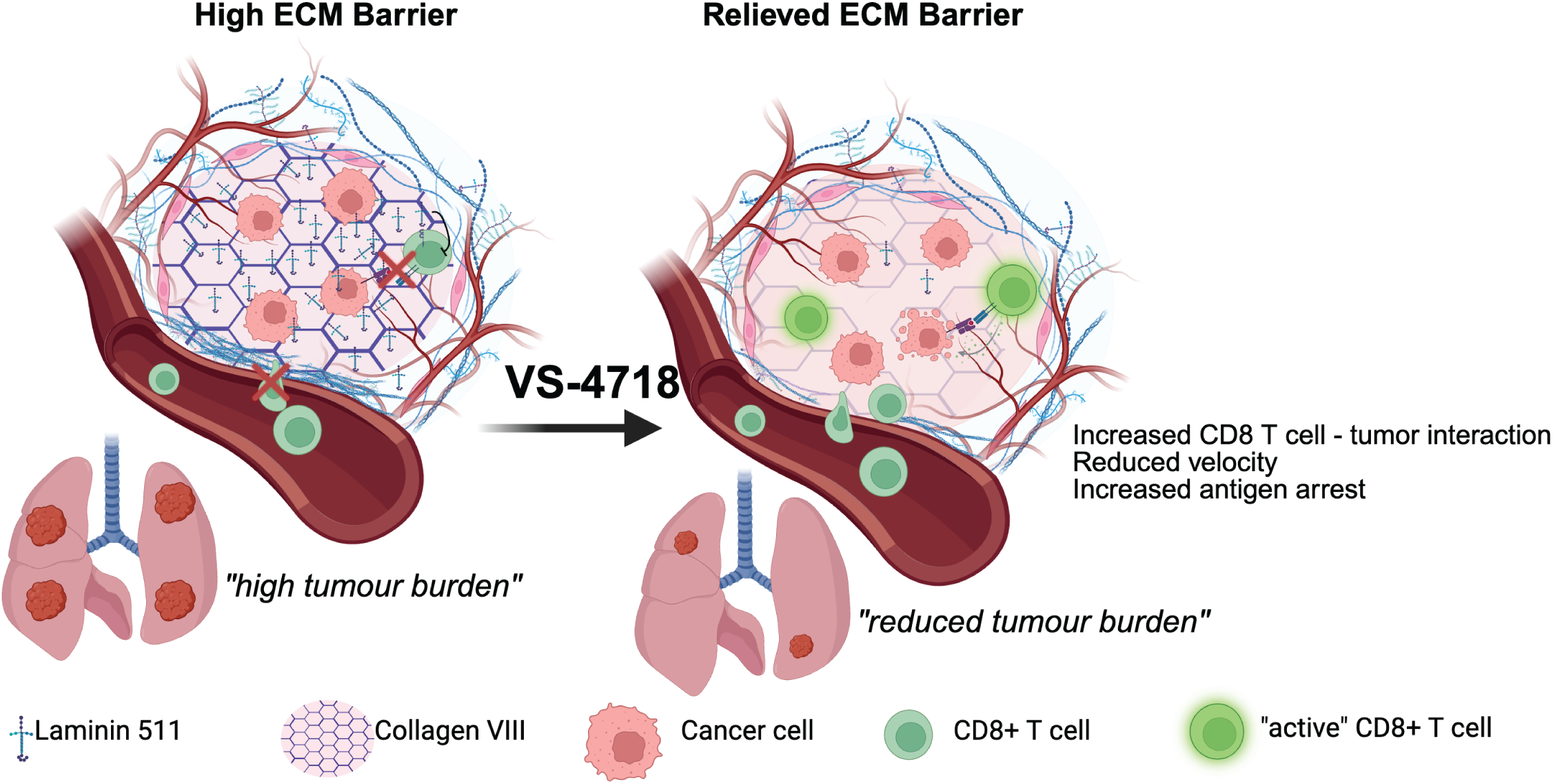
Schematic of FAK inhibition driven modulation of ECM to alter CD8^+^ T cell activity and behavior promoting clearance of pulmonary metastases. FAK inhibition reduces laminin-α5 and collagen VIIIα1 within metastatic lesions, weakening basement-membrane-derived physical and inhibitory barriers, thereby improving CD8⁺ T-cell infiltration, migration, tumor-cell engagement, and cytotoxic activity to promote metastatic regression.

## DISCUSSION

Our study uncovers novel mechanistic insights into the role of FAK within metastatic breast cancer revealing for the first time how FAK-driven ECM barriers restrict T cell access and function in established metastases. We demonstrate that FAK inhibition (via VS-4718) elicits CD8^+^ T cell-dependent regression of lung metastases by remodeling both the immune landscape and the metastatic ECM. VS-4718 treatment reduces immunosuppressive myeloid and regulatory T cell populations, while downregulating basement membrane proteins, particularly laminin α5 and collagen VIIIα1, which we show directly impair CD8^+^ T cell migration and activity.

To our knowledge, this is the first use of intravital imaging to characterize therapy-induced changes in T cell dynamics within established metastatic lesions. This showed that FAK signaling has an important role in influencing T cell dynamics in the metastatic site. Indeed, a T cell velocity reduction such as we observed is a well-known indicator of increased CD8^+^ T cell activity ^46^, and is in concordance with our finding of reduced T cell exhaustion *in vivo* and increased activity *ex vivo*. Importantly, our observation that VS-4718 had no effect on T cell velocity in tumor-adjacent tissue rules out a possible direct effect of VS-4718 on CD8^+^ T cell dynamics.

Checkpoint blockade, particularly the PD-1/PD-L1 axis, is also known to similarly influence CD8^+^ T cell dynamics. PD-1 signaling suppresses CD8^+^-triggered Ca2^+^ influx and dampens the TCR-driven “stop signal”, thereby shortening interaction times with MHC-I^+^ target cells and permitting more rapid, less confined migration ^66^. Accordingly, blockade of PD-1/PD-L1 increases antigen-induced arrest, reduces velocity, stabilizes T cell-APC/tumor cell synapses, and increases cytokine production, establishing that PD-1/PD-L1 engagement serves as a major negative-feedback mechanism restraining antigen responsiveness in primary tumors ^46,67^.

Interestingly, in our model, anti-PD-1 alone did not significantly alter the velocity of motile CD8^+^ T cells, but did decrease the percentage of motile cells, consistent with prior reports of antigen-driven arrest. However, the combination of VS-4718 and anti-PD-1 did reduce turning frequency, boost antigen-driven arrest, and, enhance stable, longitudinal interactions with tumor cells (a known characteristic of effective cytotoxic killing ^47^) and increased cytotoxicity. This is the first demonstration that *ex vivo* lung slices, that preserve the architecture and ECM composition of established metastatic lesions, can allow detailed temporal analysis of CD8+ T cell dynamics and cytotoxic activity providing a valuable adjunct to intravital imaging.

Our findings identify FAK as a key regulator of the metastatic ECM, and add to the emerging evidence that the ECM is not merely a structural scaffold, but actively regulates immune cell infiltration, dynamics, and activation in tumors ^5,14^. In primary tumors, FAK has been found to be associated with increased expression of collagen I and collagen IV ^17^. However, VS-4718 treatment did not cause significant changes in secretion of these collagens in our model. Instead, we identified collagen VIIIα1 and laminin α5 (a subunit of the basement membrane protein laminin-511) as previously unidentified targets of FAK inhibition. Both collagen VIIIα1 and laminin-511, key basement membrane components enriched around blood vessels ^59,68^, are overexpressed in various cancers, including breast cancer^55,69,70^.

Laminin-511 regulates leukocyte transmigration both directly and indirectly through stabilizing VE-Cadherin and endothelial barrier function, thereby restricting extravasation ^59,61^. Its downregulation upon VS-4718 treatment thus likely contributes to the enhanced CD8^+^ T cell extravasation we observed in metastatic lung lesions, consistent with its previously observed ability to limit immune cell entry into tissues ^71^. Moreover, *COL8A1* expression correlates with increased macrophage and Treg infiltration into tumors^72^, supporting its role as a matrix-derived barrier to effective anti-tumor immunity. While both collagen VIIIα1 and laminin α5 expression is primarily perivascular in tumor adjacent lung tissue, metastatic lesions show marked upregulation with widespread distribution throughout the tissue in the PyMT and 4T1 metastatic lesions. This intratumoral localization suggests these basement membrane proteins restrict immune cell migration through mechanisms beyond their perivascular barrier function.

We further observed that collagen VIIIα1 and laminin-511 directly reduce CD8⁺ T cell activation. Although a direct link between collagen VIIIα1 and CD8⁺ T cell activity has not previously been established, collagen VIIIα1 may contribute to CD8⁺ T cell dysfunction and exhaustion by serving as a ligand for inhibitory immune receptors. LAIR-1, for example, has been found to bind to the conserved Gly-Pro-Hyp collagen repeat found in most collagens^73^. This supports a model in which VS-4718-mediated downregulation of collagen VIIIα1 (and potentially other related collagens) alleviates receptor-mediated suppression of CD8^+^ T cells, restoring cytotoxic T cell activity. These ECM changes, combined with VS-4718-modulated cytokine secretion by breast cancer cells, likely contribute to the observed decrease in macrophage and regulatory T cell populations in the metastatic lungs and may further influence responsiveness to systemic therapies. Indeed in the I-SPY 2 trial, elevated pre-treatment expression of *COL8A1* and *LAMA5* was associated with residual disease following combination chemotherapy and immunotherapy, indicating that ECM composition may influence treatment efficacy. While these observations are correlative and limited by cohort size, they support a model in which a dense, basement membrane–rich ECM constrains both immune-mediated tumor clearance and therapeutic responsiveness. Larger prospective studies across multiple metastatic contexts will be required to validate these associations and to determine whether FAK inhibition broadly modulates ECM composition and enhances immunotherapy responses in patients.

Importantly, laminin α5 and collagen VIIIα1 represent only two components of a broader ECM signature modulated by VS-4718, several elements of which have been independently linked to metastatic progression and therapy resistance. Other downregulated proteins, including MMP3, COL6A3 and LOXL4, are similarly implicated in metastatic progression. MMP3 facilitates ECM degradation and epithelial to mesenchymal transition ^74,75^, whereas COL6A3 ^76,77^ and LOXL4 ^78^ contribute to macrophage-infiltration, pro-tumor microenvironments and matrix remodeling that supports dissemination. These findings highlight how VS-4718 reshapes the broader ECM network that regulates both tumor cell behavior and immune surveillance.

The lung is a uniquely matrix-rich organ with a highly specialized basement membrane architecture and distinctive mechanical and biochemical properties that may influence therapeutic responses differently than in primary tumors. Unlike many other tissues, the lung relies on a highly elastic, laminin-rich basement membrane to maintain alveolar structure. Our analysis of patient RNA datasets shows that *LAMA5* is elevated in many metastatic sites (**Supplementary Fig. 20**), raising the possibility that FAK-dependent ECM remodeling may provide a greater therapeutic benefit in these organs as well. This also highlights an important direction for future studies: determining whether FAK-regulated ECM components similarly contribute to immune cell trafficking in other metastatic sites.

Building on these observations, an important future direction will be the investigate the value of the FAK-dependent ECM score in a wider set of patient samples as a biomarker to potentially guide patient stratification. Our data highlight laminin α5 and collagen VIIIα1 as ECM components associated with immune exclusion, metastatic progression, and reduced therapeutic responsiveness. Quantitative assessment of these factors, either individually or as part of a composite ECM score, may help identify patients with ECM-dense metastatic niches who are less likely to respond to chemotherapy or immunotherapy alone and who may benefit from combination strategies incorporating FAK inhibition.

The recent FDA approval of the FAK inhibitor defactinib (combined with the RAF/MEK inhibitor avutometinib) for KRAS-mutant low-grade serous ovarian cancer underscores the emerging clinical relevance of FAK-targeted therapies^79^. Indeed, multiple FAK inhibitors are currently being evaluated in clinical trials across pancreatic^80^, ovarian^81^, and other solid cancers^15^. Our work lends support to this clinical advancement by providing mechanistic insight into how FAK inhibition may synergize with PD-1 blockade, particularly in metastatic settings characterized by dense ECM deposition. These findings offer a rationale for integrating clinically viable FAK inhibitors into immunotherapy regimens for metastatic breast cancer.

## METHODS

### Mouse Strains

C57BL/6JCrl and Balb/cAnNCrl mice were purchased from Charles River Laboratory. OT-1 C57BL/6 (#003831) mice were both purchased from Jackson or bred in-house. CD-1 Nude mice were purchased from Charles River Laboratory. Mice were acclimatized in the local animal facility for at least 1 week prior to experiment. Mice were housed in a conventional unit in Individually Ventilated Cages (IVCs) with controlled temperature and humidity (20-24 **°**C/45-65%) environment on a 12-hour light-dark cycle with food and water ad libitum. All female mice were age-matched (6-8 weeks) at the time of cell injection. Experiments were approved under the IACUC and the University of Edinburgh ethical approval and were carried out in accordance with the United Kingdom Animal Scientific Procedures Act (1986). Mice were randomly allocated to groups using an equal weight distribution between groups prior to treatments.

### Generation and Culture of Cell Lines

Lego Cerulean 2 (LeGO-Cer2, Plasmid #27388, Addgene) and pcDNA OVA (pcDNA2-OVA, #64599, Addgene) were purchased from Addgene. HEK293T cells were transfected with 3^rd^ Generation Helper plasmid using Lipofectamine^TM^ 3000 transfection reagent (#L3000001, ThermoFisher Scientific). Lentivirus was harvested 48 h after transfection and concentrated using a lenti-x concentrator 1:10. MMTV PyMT cells at a confluency of 60% were transduced first with Cerulean 2 and then transfected with pcDNA OVA with Lipofectamine^TM^ 3000 transfection reagent. Cer2^+^ cells were sorted using FACSaria for top 90-100% fluorescence intensity and OVA positive cells were selected with antibiotic resistance to G418 at 300 μg/mL. Cells were cultured in DMEM supplemented with 10% FBS and 1% penicillin/strep.

PTK2 was CRISPR edited using two pre-designed guide Alt-R CRISPR-Cas9 crRNA; Mm.Cas9.PTK2.1.AA (TCGAGTACTAAGACTCACCT) and Mm.Cas9.PTK2.AB (GTAAAGCACGTGGCCTGCTA). Briefly, crRNA was reconstituted at 100 µM using nuclease-free duplex buffer. For gRNA duplex formation, 1.1µL of crRNA was mixed with 1.1µL of Alt-R CRISPR-Cas9 tracrRNA (Integrated DNA Technologies, #1072533) and annealed using a thermocycler with the following program: 95°C for 5 minutes, step-cool down from 95°C to 25°C (ramp rate = 0.1°C/s) and finally cooled to 4°C (ramp rate 0.5°C/s). The duplex was incubated with 10µg of Alt-R S.p. Cas9 Nuclease V3 (Integrated DNA Technologies #: 1081059) at room temperature for 10 minutes to allow Cas9:RNP formation and then stored on ice. Control Cas9:RNP lacking a crRNA was performed in parallel.

Nucleofection was performed using the Amaxa SE Cell Line 4D-Nucleofector X Kit S (Lonza #: V4XC-1032). Cells for nucleofection were harvested using standard cell culture protocol and 400,000 cells were centrifuged and the pellet resuspended in 20µL of SE buffer. Cas9:RNP complexes for the same target gene were pooled and mixed with the cell suspension and a final concentration of 5µM of Alt-R Cas9 Electroporation enhancer (Integrated DNA Technologies, #1075916) was added. The cell suspension was transferred to the well of a 16-well microcuvette strip and nucleofection performed using a 4D Nucleofector and program EH-109. Immediately after nucleofection, cells were transferred to a well of a 6-well plate in complete growth media. Cells were cultured as normal and successful CRISPR gene editing confirmed using western blotting.

### Cell Culture

Cer2-OVA MMTV-PyMT cell line and FAK^-/-^ Cer2-OVA MMTV-PyMT cell line, which is a syngeneic breast cancer cell line to C57BL/6 mice, has been generated as described here from spontaneous MMTV-PyMT tumors. Cell lines were cultured in DMEM supplemented with 10% heat-inactivated FBS (#F2442, Sigma) and 100 U/mL penicillin and 100 μg/mL streptomycin (#P0781, Sigma) (Complete DMEM). 4T1 cells were provided by Chartinum Chutoe and Alex von Kriegsheim (purchased from ATCC, UK) and cultured in RPMI supplemented with 10% FBS and penicillin and streptomycin. Cells are passaged at a density between 20-80%. For passage, cell media is aspirated and PBS without divalent cations added. Cells are then detached with 0.05% Trypsin-EDTA (Gibco, UK) for 15 min at 37 °C in 5% CO_2_. Trypsin is neutralized with complete DMEM and cells seeded at a density of 15% in complete DMEM. Cells were used up to passage 20 for *in vivo* experiments and passage 30 for *in vitro* experiments.

### Tumor Cell Implantation

To inoculate metastasis in the mouse, Cer2-OVA MMTV-PyMT and 4T1 cells were expanded in Dulbecco’s Modified Enhanced Medium (DMEM), supplemented with 10% Fetal Bovine Serum (FBS), 100 units/mL penicillin and 100 μg/mL streptomycin at 37 °C in 5% CO_2_ (ThermoFisher Scientific, UK). To prepare cells for injection, cells were detached in 0.05% Trypsin-EDTA (Gibco, UK) for 15 min at 37 °C in 5% CO_2_, cells were transferred in sterile PBS without divalent cations (PBS^-/-^) (Gibco) and centrifuged at 300g, 5 min at room temperature (r.t.). Cells were resuspended in PBS-/- and passed through a 40 μm filter, this process was carried out twice prior to concentration of Cer2-OVA PyMT cells at 5x10^6^ cells/mL and 4T1 cells at 2.5x10^6^ cells/mL in sterile PBS^-/-^. 1x10^6^ cells and 5x10^5^ cells per mouse were injected via tail vein for Cer2-OVA MMTV-PyMT cells and 4T1 cells, respectively. Cells were routinely tested for Mycoplasma contamination by PCR and were used up to passage 20. Metastatic lungs were taken between 5-6 weeks after implantation of Cer2-OVA MMTV-PyMT cells and 19 days past implantation of 4T1 cells.

### Treatment with VS-4718 and anti-PD-1 in vitro and in vivo

FAK inhibitor (FAKi) VS-4718 was provided by Selleckchem (#S7653), it is a dual inhibitor with higher affinity to FAK1/FAK compared to FAK2/PYK2 kinases ^82^. To enhance its biostability, it is formulated at 75 mg/kg for weekdays and 150 mg/kg at the weekends in vehicle (0.5% hydrocarboxymethyl cellulose) (Sigma Aldrich) in sterile water. VS-4718 or vehicle is administered twice a day on weekdays and once a day on weekends by oral gavage. For *in vitro* studies, VS-4718 was dissolved in DMSO and used between 300 nM to 1000 nM. For immunotherapy regimen, anti-PD1 (clone RMP1-14, BioXCell) was given intraperitoneally (i.p.) every 48 h at 100 μg. Anti-PD1 was used at 10 μg/mL for *in vitro* studies and administered every 48 h.

### Isolation and Culture of CD8^+^ T Cells

Mouse OT-1 T cells were isolated from the spleen of female OT-1 C57BL/6 (8-12 weeks of age) by negative selection using a CD8 T Cell Isolation Kit (MACS, Miltenyi Biotec). In brief, spleen was passed through a 40 μm filter (ThermoFisher) using the stamp of a 5 mL syringe and washed in PBS/1% BSA. Red blood cell lysis was carried out with ACK Lysing Buffer (ThermoFisher) for 5 min on ice. Cell lysis was stopped with 1:10 PBS/1% BSA. Cells were resuspended at 30 μL MACS buffer (PBS^-/-^, 2 mM EDTA, 0.5% BSA) per 1x10^7^ and 10 μL per 1x10^7^ cells antibody cocktail was added. After 5 min on ice, 30 μL of MACS buffer and 20 μL of beads were added per 1x10^7^ cells and incubated for 10 min. LS columns were calibrated with 3 mL MACS buffer and cells were recovered with 3 mL MACS buffer. CD8^+^ T cells were then resuspended at 1x10^6^/mL in DMEM or RPMI supplemented with 10% FBS, 1% non-essential amino acids (NEAA, Gibco), 5 μM – 14 μM β-mercaptoethanol (Sigma), 1% L-glutamine, 100 units/mL penicillin and 100 μg/mL streptomycin. 80 units/mL human/mouse IL-2 (Peprotech), 5 μg/mL anti-CD3 (eBioscience) and 5 μg/mL anti-CD28 (eBioscience) were added and cells cultured at 37 °C in 5% CO_2_.

### CD8^+^ T cell Migration Assay

For migration assays, PyMT Cer2 OVA cells were seeded at 5x10^4^ cells/well in 24-well plates (1 mL/well, DMEM with 10% FBS) and cultured for 24 h at 37 °C, 5% CO₂ until 70-80% confluence. Medium was then replaced with 600 μL RPMI-1640 supplemented with 1% FBS, 1% L-glutamine, and 1% penicillin-streptomycin (migration medium) with or without VS-4718 or AMP945 (300 or 600 nM), and cells were incubated for a further 24 h. 3 μm pore size Transwell inserts (Corning, 6.5 mm diameter) were coated with Laminin-511 and Collagen VIIIα1 at the final densities of 1 µg/cm². Activated CD8⁺ T cells were harvested and resuspended in RPMI-1640 supplemented with 1% FBS at 1x10⁶ cells/mL. 100 μL of T cell suspension (1x10⁵ cells) was added to the upper chamber of Transwell inserts and 600 μL RPMI-1640 supplemented with 10% FBS was added to lower chamber. After 5 h at 37 °C, 5% CO₂, the lower chamber medium was collected, centrifuged at 300 g for 5 min, and cells were resuspended in 30-50 μL medium for counting by hemocytometer. Migrated cells were calculated as (migrated cells in experimental group - migrated cells in negative control), and normalized to non-ECM coating control. Negative controls consisted of inserts placed in 1% FBS RPMI-1640 medium to account for background migration.

### Random Migration CD8^+^ T Cell Assay

Activated CD8⁺ T cells were isolated as described previously and plated onto 24-well plates coated with Laminin-511 or Collagen VIIIα1. ECM solutions were diluted in serum-free medium to final coating densities of 1 µg/cm² Laminin-511 and Collagen VIIIα1, applied overnight at 4 °C, and washed once with PBS. T cells were labeled with 250 nM CellTracker™ Deep Red for 15-30 min at 37 °C, washed, and resuspended in phenol-free medium. A total of 500 cells/well were added to coated wells and imaged by time-lapse microscopy for 6 h at 5-min intervals. Single-cell trajectories were extracted using TrackMate, and tracks with >11 spots were analyzed to quantify mean speed, directional change rate (here termed: average turning frequency), and linearity of forward progression.

### CD8^+^ T Cell Adhesion Assay

ECM solutions were diluted in serum-free medium to final coating densities of 1 µg/cm² Laminin-511 or Collagen VIIIα1, applied overnight at 4 °C, and washed once with PBS. Activated CD8⁺ T cells were labelled with CellTracker™ Deep Red dye (250 nM, 15-30 min, 37 °C; Invitrogen, Cat #C34565, Lot#2845270) and added to the coated plates, followed by incubation for 90 min at 37 °C. Non-adherent cells were removed by gentle washing with PBS twice, and adherent cells were imaged using the Incucyte SX5. Positive cells were quantified from 9 random fields per well at 20x magnification.

### Generation and Culture of Precision Cut Lung Slices

To generate metastatic lung slices for co-culture, lungs were harvested 5 weeks post implantation of Cer2 OVA PyMT tumor cells through tail vein injection into mice. Mice were positioned with the dorsal side facing downward and the chest was sterilized with 70% ethanol. A lateral incision from the lower end of the rib cage to the upper end of the throat was performed. The chest cavity was opened along the midline from abdomen to chin and the front ribs removed to generate sufficient space for lung inflation. Lungs were perfused through the right cardiac ventricle with 10 mL ice cold PBS using 10 mL syringe fitted with 25 G needle (#305122, BD). Trachea was exposed by careful resection of muscle tissue and the trachea was gently lifted with forceps and a loose knot positioned with a 5-0 braided silk thread around it. A 21 G x 1 in (#305165, BD) needle was laterally inserted into trachea and needle secured by tightening the knot. A syringe with 2% preheated LMP Agarose (#A4108, Sigma) was connected and lungs inflated with approximately 1 mL LMP Agarose. Lung was transferred into ice-cold PBS on ice. Lung lobes were separated and 300 μm slices carried out using a vibratome or compresstome.

### Intravital Imaging of CD8^+^ T cells and metastatic lung slice co-culture

Precision-cut lung slices (PCLS) were prepared from tumor-bearing C57BL/6J mice. Slices were transferred to 24-well plates containing 1 mL complete DMEM and recovered overnight at 37 °C, 5% CO₂. For imaging, individual slices were moved to glass-bottom 24-well plates and secured beneath a custom stainless-steel window (8 mm) sealed peripherally with 2% agarose to prevent drift. Fluorescently labelled CD8⁺ T cells (100 μL; applied to the slice surface) were allowed to infiltrate for 15 min at 37 °C, after which 900 μL phenol-red-free complete DMEM (supplemented with 20 mM HEPES, 1% non-essential amino acids, 1% L-glutamine, 5 μM β-mercaptoethanol, 1% penicillin-streptomycin) containing 40 IU/mL IL-2 ± low-dose anti-CD3 (2 μg/mL) and anti-CD28 (5 μg/mL) and test agents was added. Where indicated, nuclei and apoptosis were labelled with NucSpot Live 650 (1 μM) and Apotracker Red (150 nM), respectively. Time-lapse imaging was performed on a confocal or two-photon microscope maintained at 37 °C, 5% CO₂ using a 25x/1.05 NA water-immersion objective; Z-stacks (∼50 μm in 3-5 μm steps) were acquired just below the slice surface. T-cell motility and CD8⁺ T cell-tumor cell contact dynamics were quantified from registered fields over time using established particle/track-based analyses.

### Surgery for implantation of the window for high resolution of the lung (WHIRL)

Surgical implantation of the WHIRL including window passivation and surgical procedure and after care has previously been described ^83^. In brief, lung windows were passivated acid-alkaline-acid (A-A-A) method adopted from standards released by the American Society for Testing and Materials (ASTM A967-13). Prior to surgery, mice were anaesthetized with 5% isoflurane (#029405, Henry Schein) and chest hair was depilated. Mice were intubated using a 20 G catheter (#26746, Excelint international) that was then secured around the snout with a 2-0 silk suture (#LA556 Perma-Hand Silk, Ethicon, Inc). Mice were placed in the right lateral decubitus position on a heated surgical platform and the limbs secured cranially and caudally with paper tape. A catheter was connected to mechanical ventilator (MouseVent for PhysioSuite, Kent Scientific, Inc.) and the mouse ventilated throughout surgery. Anesthesia was reduced to 2.5-3% isoflurane and the surgical field was disinfected with 70% chlorhexidine gluconate (ChloraPrep, Becton Dickenson, Inc.). A pre-surgical analgesic (buprenorphine) was administered and the skin and muscle resected between ribs 2 and 3 on the upper left chest region, ∼7 mm lateral to the sternum and superior to the last floating rib. The thoracic cavity was opened with a 5 mm circular incision using a biopsy punch, exposing lung tissue. A purse-string suture (774B, Ethicon,Inc.) was run 1 mm from the edge of the circular hole, intercalating between the ribs. The stainless steel window frame was inserted so that the ribs and suture fit snugly into the 1.25 mm groove. Lung tissue was adhered with cyanoacrylate (LOCI362589, Henkel Adhesive) adhesive dispensed on the underside of the window frame while applying positive end-expiratory pressure (PEEP). Next, a 5 mm diameter #1.5 circular coverslip (72296-05, Electron Microscopy Sciences) was coated with a thin layer of adhesive and used to seal the aperture of the window frame while adhering the exposed tissue to the glass by increasing PEEP briefly. Excess air in the thoracic cavity was evacuated using an insulin syringe placed through the diaphragm just below the xiphoid process and angled cranially toward the left shoulder. At the end of surgical procedure, mice were extubated and allowed to recover from anesthesia in a heated chamber, buprenorphine (NDC 0409-2012-32, Hospira) was administered subcutaneously daily for pain control.

### Intravital Imaging of CD8^+^ T Cell and Cancer Cell Interaction

Mice were anaesthetized using 2% isoflurane and retroorbitally injected with 50 μL 20 mg/mL 155 kDa TMR-Dextran (#T1287-50MG, Sigma) to visualize blood flow ^84^. For long time-lapse imaging, either a tail-vein catheter was inserted or PBS was administered through peritoneal injection for periodic hydration. A fixturing plate was placed between the skin and the window frame. The mouse was inverted and placed on the microscope stage, fixturing plate was taped to the stage using paper tape. Physiological temperatures were maintained through environmental enclosure and heated control, vitals were monitored using pulse oximeter (MouseStat for PhysioSuite, Kent Scientific). Imaging was performed on a previously described, custom-built, two-laser multiphoton ^85^. All images were acquired in 16-bit using 25x 1.05 NA objective lens and acquired with two frame averages.

### Quantification of CD8^+^ T Cell Movement

Image analysis was performed in ImageJ/Fiji as previously described ^84^. All presented images represent raw microscope data with minimal linear adjustments to brightness and contrast and processed using our previously described blood-averaging algorithm to suppress erythrocyte-derived serum signal fluctuations ^83^. Time-lapse datasets acquired 1 h after CD8⁺ T cell injection were assembled into hyperstacks and residual x-y drift was corrected using the HyperStackReg plugin (https://github.com/ved-sharma/HyperStackReg). Three-dimensional renderings were generated in Imaris (v9.9). CD8⁺ T cell-tumor cell interactions were classified as “no contact,” “contact,” or “long-duration contact” (>20 min). Channels were separated, and the CD8⁺ T cell channel was thresholded to segment individual cells. Regions of interest (ROIs) encompassing motile CD8⁺ T cells were manually defined and imported into the ROI_Tracker plugin in Fiji for automated tracking^85^. Track data were exported for quantification of velocity, turning behavior, and interaction dynamics.

### Immunofluorescence of Metastatic Breast Cancer Lungs

After paraffin embedding, tissues were sectioned with 3 μm thickness. Tissue sections were deparaffinized and rehydrated with twice 100% xylene for 5 min, twice 100% ethanol for 5 min and 3 min each in 70% and 50% ethanol. Tissues were transferred into distilled water prior to antigen retrieval. Sodium citrate buffer, pH 6 was then used for antigen retrieval and antigen retrieval was carried out for 5 min in the pressure cooker. Sections were then allowed to slowly transfer back to room temperature prior to three washes in PBS containing 0.05% Tween 20 (PBS-T) for 5 min. PAP pen (DAKO) was used to generate hydrophobic barrier around tissue and slides were blocked for 1 h at room temperature in PBS containing 0.05% Tween 20, 5% bovine serum albumin and either 3% donkey or goat serum depending on species of secondary antibody. Sections were incubated overnight at 4 °C with primary antibodies diluted in blocking buffer: Ki-67 (rat anti-mouse, eBioscience, Cat#14-5648-82, 1:100), CD8 (rabbit anti-mouse, Abcam, Cat#ab237723, 1:200), CD45 (goat anti-mouse, R&D Systems, AF114, 1:500), GFP (chicken anti-mouse, 1:100, Novus Biologicals, #NB100-1614), CD4 (rat anti-mouse, eBioscience, Cat#14-9766-82, 1:200), and FoxP3 (rabbit anti-mouse, clone D608R, Thermo Fisher, Cat#14-5773-12, 1:200). After three PBS-T washes, sections were incubated with secondary antibodies for 2 h at room temperature, washed again in PBS-T, and counterstained with Hoechst 33342 (1 μg/mL, 20 min). Slides were washed three times in PBS-T and mounted with ProLong Glass Antifade (Invitrogen). Images were acquired on a Zeiss AxioScan with a 20x objective, and quantification was performed using QuPath.

### Immunohistochemistry for ECM Components

Formalin-fixed, paraffin-embedded tissue sections (4 μm) were deparaffinized in xylene (2 x 5 min), rehydrated through graded ethanol (100%, 95%, 70%, 50%, 2 min each) and rinsed in distilled water. Antigen retrieval was performed in 0.1 M sodium citrate buffer (pH 6.0) using a pressure cooker (15 min at ∼95 °C), followed by cooling at room temperature for 30 min. Sections were washed in TBST (0.5% Tween 20 in TBS, 2 x 5 min), endogenous peroxidase activity was quenched with DAKO Peroxidase Blocking Solution (5 min), and non-specific binding was minimized with protein block (10 min, RT). Slides were incubated overnight at 4 °C with primary antibodies against laminin α5 (polyclonal, Bioss, Cat# BS-1086R, 1:500) or collagen VIIIα1 (rabbit polyclonal, Abcam, Cat# ab236653, 1:300) diluted in antibody diluent. After washing in TBST (3 x 5 min), sections were incubated with DAKO EnVision+ labelled polymer anti-rabbit secondary antibody (2 h, room temperature), washed, and developed with DAB chromogen (1:50 in substrate buffer, 10 min). Nuclei were counterstained with Mayer’s haematoxylin (2 min), blued in Scott’s tap water (30 s), dehydrated in ascending ethanol (70%, 95%, 100%; 2 min each), cleared in xylene (2 x 5 min), and mounted in DPX mounting medium.

### Quantification of Metastatic Burden and Immune Cells

After completion of treatments, lungs were harvested and immersed in 10% formalin in a volume ratio of tissue to formalin of 1:10. The tissues were processed for histological examination by paraffin embedding. Sections were obtained at 0, 200, 400 μm intervals and stained for H&E. Metastatic foci were counted in the tissue section of the entire lung in the three slides and averaged for the number of metastatic lesions. For metastatic tumor burden, the area of metastatic foci was calculated and a ratio of area of metastatic foci to area of the whole lung was determined. Metastatic lesions were confirmed through comparison with immunofluorescence staining at first interval for anti-GFP and anti-P53. Staining was used to validate the stratification of clusters of cells as metastatic lesions.

### Flow Cytometry Analysis of Metastatic Lungs

CD8⁺ T cell activation and surface marker expression on tumor cells was characterized by flow cytometry. In brief, adherent cells were trypsinized and resuspended in flow cytometry buffer (PBS without calcium and magnesium containing 0.1% BSA). Suspension cells were centrifuged at 300 g for 5 min and resuspended in flow cytometry buffer. Cells were washed once in PBS and resuspended in PBS. Cells were stained with 1 μM Live/Dead Blue viability dye (BioLegend, Cat #423102) for 30 min at room temperature in the dark. Following viability staining, cells were washed, resuspended in flow cytometry buffer, and stained for surface marker expression. For intracellular staining, cells were fixed and permeabilized with the FOXP3/Transcription Factor Staining Buffer Set (Thermo Fisher Scientific, Cat #00-5523-00), starting with fixation/permeabilization buffer for 45 min at 4 °C in the dark. Cells were centrifuged at 300g for 5 min at 4 °C and resuspended in permeabilization buffer. Intracellular antibodies were added and incubated for 45 min at 4 °C in the dark. Cells were then washed once in permeabilization buffer and once in flow cytometry buffer before acquisition. For characterization of immune cell populations in metastatic lungs, lungs were harvested and transferred into RPMI buffer. Two lobes were cut into small pieces and transferred into digestion buffer consisting of RPMI supplemented with collagenase/hyaluronidase (Stemcell Technologies, Cat #07912). Lung tissue was digested for 30 min at 37 °C with gentle agitation, filtered through 40 μm strainers (Corning, Cat #352340), and washed in PBS containing 1% BSA. For splenic samples, spleens were dissociated through a 40 μm strainer into PBS with 1% BSA. Blood, lung cell suspensions, and splenocytes were treated with ACK lysis buffer (Thermo Fisher Scientific, Cat #A1049201) for 5 min at 4 °C to lyse erythrocytes and washed in PBS with 1% BSA. Cell suspensions were counted, and 1x10^6^ cells per sample were used for staining. Two immune cell panels were employed to characterize lymphoid and myeloid subsets. For lymphoid subsets, cells were stained with Live/Dead Blue, and for myeloid subsets, cells were stained with Live/Dead Near-IR (Thermo Fisher Scientific, Cat #L34975). The lymphoid panel included CD3, CD4, CD8, CD25, PD-1, LAG3, TIM-3, FoxP3, IFN-γ, TNF-α, and Granzyme B. The myeloid panel included CD11b, Ly6C, Ly6G, F4/80, CD11c, and MHC-II, as well as extended markers including CCR2, CD206, CX3CR1, and CD103. Antibody clones, fluorophores, and catalogue numbers are listed in **Supplementary Table 3**. Samples were acquired on a BD LSRFortessa cell analyzer equipped with five lasers, and compensation was applied using UltraComp eBeads (Thermo Fisher Scientific, Cat #01-2222-42). FlowJo v10.10.0 (BD Biosciences) was used for data analysis, with gating strategies guided by fluorescence minus one (FMO) controls.

### Flow cytometry for *in vitro* activation expression

For ECM-based activation assays, 96-well plates were coated with laminin 511 and collagen VIIIα1 together with anti-CD3 (5 μg/mL) and anti-CD28 (5 μg/mL) at 4 °C for 24 h, washed, and seeded with 5 x 10⁴ CD8⁺ T cells/well in complete medium (RPMI-1640 with 10% FBS, 80 U/mL IL-2, 14 μM β-mercaptoethanol (Sigma), 1% L-glutamine, 1% penicillin-streptomycin) for 48 h with or without VS-4718 (300 nM). At 24, 48, and 72 h, cells were harvested, washed with PBS (DPBS without Ca²⁺/Mg²⁺), and stained with Zombie UV viability dye (1:1000 in PBS, 30 min, RT, protected from light). Cells were then washed with FACS buffer (DPBS + 1% BSA + 0.1% sodium azide) and stained for extracellular markers (CD3, CD8, PD-1, LAG3, TIM-3, PD-L1, CD25, and CD305) for 30 min at 4 °C in the dark. After washing, cells were fixed and permeabilized using commercial fix/perm buffer (prepared 1:3 concentrate:diluent; 45 min, RT, dark), washed in perm buffer (1:10 dilution in dH₂O), and stained for intracellular markers (IFN-γ, TNF-α, Granzyme B) for 45 min at 4 °C. Finally, cells were washed once in perm buffer, once in FACS buffer, and resuspended in 300 μL FACS buffer before acquisition on a BD LSR Fortessa.

### Western Blotting

Total protein from tumor cells was obtained by lysis of cells in 1x RIPA buffer (#20-188, Sigma) containing 1x protease inhibitor cocktail (#P8340, Sigma), 1x phosphatase inhibitor cocktail (#524624, Sigma) and 1 mmol/L PMSF (#52332, Sigma). In brief, cell media was aspirated and remaining protein removed with ice-cold PBS, RIPA supplemented with inhibitor cocktail was added and incubated for 15 min on ice. Cell scrapers were used to lift cells from plate and transferred into Eppendorf tubes. After centrifugation for 15 min, 4 **°**C, 15000 x g, protein concentration of supernatant was determined with Pierce^TM^ Bradford Protein Assay (Cat #23200, ThermoFisher Scientific). 15 μg protein per sample was separated on 8-16% protein gel (Cat #4568106, Biorad) and transferred onto a nitrocellulose membrane. Membrane blocking was carried out for 1 h, room temperature in 5% BSA in 0.05% TBS-T, subsequent antibody incubations were all performed in blocking solution. The primary antibodies were used at 1:1000 monoclonal rabbit anti-GAPDH (Cat #5174S, Cell Signaling); rabbit anti-FAK (Cat #3285S, Cell Signaling) and rabbit anti-pFAK (Cat #3283S, Cell Signaling). Secondary antibody mouse anti-rabbit HRP (Cat #7074S, Cell Signaling) was used at 1:5000 for 1 h, room temperature. Immunodetection was performed with Pierce ECL Western Blotting Substrate (Cat #32106, ThermoFisher Scientific).

### Cytokine Array

Cer2-OVA MMTV-PyMT cells and FAK^-/-^ Cer2-OVA MMTV-PyMT cells were cultured for 24 h with or without VS-4718 (1 μM). At 48 h, conditioned supernatant was collected for cytokine analysis. All conditions were analyzed in biological triplicates (n = 3). A cytokine assay was carried out in microarray format using validated capture/detection antibody pairs (R&D Systems). For each sample, a selected panel of 64 capture antibodies was printed as four replicate sub-array sets on a single nitrocellulose-coated glass slide (Supernova Grace Biolabs) using a Quanterix-2470-microarrayer and 185 μm printing pins. After carrying out a four-step dilution series for each sample (25%, 12.5%, 6.25% and 3.125% in phosphate-buffered saline), each dilution was incubated with an identical sub-array in order to generate four readout values per sample. After blocking, sample incubation and repeated wash steps, arrays were incubated with a specific detection antibody, which was biotin labelled. A final incubation with fluorescently labelled streptavidin generated the signal for all samples and this was quantified using an InnoScan710IR scanner (Innopsys) and analyzed using Mapix software (Innopsys).

### Proteomics

Cer2-OVA MMTV-PyMT cells were cultured for 24 h or 48 h with or without VS-4718 (1 μM). At 24 h, cells were harvested, and at 48 h, conditioned supernatant was collected for proteomic analysis. All conditions were analyzed in biological triplicates (n = 3). Protein from supernatant was extracted using TCA precipitation. Cells were lysed in PAC buffer (5% SDS, 100 mM Tris-HCl pH 8.5, 1 mg ml⁻¹ chloroacetamide, 1.5 mg ml⁻¹ TCEP), heated at 95 °C for 30 min, and sonicated. Lysates were processed for automated digestion (8 h) on the KingFisher Duo (Thermo Fisher Scientific) using MagReSyn HILIC beads (ReSyn Biosciences) and 70% acetonitrile. Washing was performed with 95% acetonitrile and 70% ethanol, and digestion was carried out with 1 µg ml⁻¹ MS-grade trypsin (Promega) in 50 mM triethylammonium bicarbonate. Reactions were stopped with trifluoroacetic acid (TFA; Thermo Scientific, 28901). Peptides were desalted on C18 StageTips and analysed on an Orbitrap Fusion Lumos Tribrid Mass Spectrometer (Thermo Fisher Scientific) coupled to an Ultimate 3000 Nano LC with a C18 packed emitter (Aurora, IonOptiks). Peptides were separated using a 70 linear gradient from 5% to 30% acetonitrile, 0.5% acetic acid wash. The mass spectrometer was operated in DIA mode, acquiring a MS 350-1650 Da at 120k resolution followed by MS/MS on 45 windows with 0.5 Da overlap (200-2000 Da) at 30 k with a NCE setting of 28. Data were analysed using DIA-NN (version 2.3.0 Academia) or FragPipe platform (version 23.1) against the Mus Musculus UniProt reference proteome. Raw data were processed with DIA-NN 2.0 against the Mus musculus UniProt/SwissProt database (reviewed, release 2014_02). Peptides were filtered at a 1% false discovery rate (FDR), LFQ intensities were calculated, and contaminants and reverse hits were excluded. Differential protein abundance was determined using MS-EmpiRe with significance thresholds set at ratio >2 or <0.5 and p<0.05. MS DAP and Perseus was used for statistical analysis. Perseus was used to filter significantly upregulated and downregulated proteins determined by MS-EmpiRe. Heatmaps were generated with Perseus using hierarchically clustered log2-transformed, median-centred protein abundances which were z-scored across rows according to group (cellular and secreted proteome). Protein symbols were converted into gene symbols and analyzed using ShinyGO 0.85 for gene ontology (GO) and KEGG pathway enrichment. STRING analysis was performed via string-db.org, and gene ontology and KEGG plots were generated using www.bioinformatics.com.cn.

### Quantitative Real-time Polymerase Chain Reaction (qPCR)

TNBC lines were grown under normal conditions and treated for 3 h with 1 μM VS-4718 in serum-free DMEM. Total RNA was isolated from cells by using RNeasy Plus Mini Kit (Cat #74134, Qiagen) and cDNA was synthesized with Maxima First Strand cDNA Synthesis Kit (#K1671, ThermoFisher Scientific). Fast SYBR Green Master Mix (Cat #4385612, Thermo Fisher Scientific) was used for qPCR and analyzed using a QuantStudio 3 real-time PCR instrument (Applied Biosystems, Thermo Fisher Scientific). The following primers were used and purchased from IDT: mouse GAPDH 5’-CATCACTGCCACCCAGAAGACTG -3’, 5’ ATGCCAGTGAGCTTCCCGTTCAG -3’; mouse CCL2 5’- GCTACAAGAGGATCACCAGCAG -3’, 5’- GTCTGGACCCATTCCTTCTTGG -3’; mouse CXCL7 5’- CTGATCCTTGTTGCGCTGGCTC -3’, 5’- GCCTGTACACATTCACAAGGGAG -3’; mouse CCL5 5’- CCTGCTGCTTTGCCTACCTCTC -3’; mouse LIX 5’- CCGCTGGCATTTCTGTTGCTGT -3’, 5’- CAGGGATCACCTCCAAATTAGCG -3’; mouse MCSF 5’- TGAGACACCTCTCCAGTTGCTG -3’, 5’- GCAATCAGGCTTGGTCACCACA -3’. The mean cycle threshold (Ct) values were then used to analyze relative expression. GAPDH was used as housekeeping gene for normalization, and 2^−ΔΔCT^ was used to determine the relative abundance of the target transcripts. Expression was compared to Cer2-OVA PyMT cells.

### SCAN-B clinical cohort analysis

Bulk RNA-sequencing data were obtained from the Sweden Cancerome Analysis Network – Breast (SCAN-B) clinical cohort^62^. Clinical annotations included tumor grade and Genius score. The Genius score is a genomic risk classifier that categorizes tumors into three discrete risk levels based on multigene expression patterns: 0 (low genomic risk), 0.5 (intermediate risk), and 1 (high genomic risk). This score reflects tumour aggressiveness and recurrence potential.

### ECM, metastatic burden and cytotoxic signatures

The ECM signature score was calculated using the first principal component (PC1) of 53 VS-4718-regulated proteins. The metastatic burden signature (MBS) was defined as a set of 49 genes previously reported to be overexpressed in high metastatic burden breast cancer PDX models^86^. Signature scores for both ECM and MBS were computed using PC1 of their respective gene sets. Cytotoxic and CD8 T cell scores were estimated using the consensusTME package (v0.0.1.9000)^87^ in R v4.1.0. Correlations between ECM gene expression and these signatures were assessed using Spearman’s rank correlation across TNBC, ER-positive, and all samples.

### Correlation analysis

Associations between ECM signature, COL8A1, LAMA5, PTK2, and clinical variables (tumor grade, Genius score) were evaluated in three patient subsets: all breast cancer cases, ER-positive tumors, and TNBC. Correlation with continuous variables was assessed using Spearman’s rank correlation, while comparisons across Genius categories (0, 0.5, 1) and tumor grades were tested using ANOVA in stats v4.1.0 package.

### Survival analysis

Overall survival (OS) was analyzed over a 10-year follow-up period. For gene signatures, expression was summarized using the first principal component (PC1). Patients were stratified into high and low expression groups based on tertile cut-offs. Within each gene-expression group, patients were further classified by CD8 T-cell score (or cytotoxic score) into high and low categories, resulting in four subgroups. Kaplan–Meier curves were generated using the survdiff function in survival v3.2.13, and hazard ratios (HR) were estimated using Cox proportional hazards models (coxph). Statistical significance was assessed using the log-rank test.

### I-SPY 2 clinical trial analysis

I-SPY 2 is a phase 3 randomized clinical trial (Clinical trial number: NCT01042379) involving 987 women with BC, including 363 patients with TNBC, randomly assigned to 13 arms with various combinations of new agents of neoadjuvant chemotherapy^88^. Expression of COL8A1 and LAMA5 was compared between responders (pathological complete response, pCR) and non-responders (residual disease, RD) in TNBC patients enrolled in two treatment arms: (i) Paclitaxel + ABT-888 + Carboplatin and (ii) Paclitaxel + Pembrolizumab. Differential expression analysis was performed using the limma package v3.50.3 in R v4.1.0.

### Statistical Analysis

All statistical analyses were performed using GraphPad Prism (v9.0). Unless otherwise stated, results are presented as mean ± SEM from at least three independent biological replicates. Normality of datasets was assessed using the Shapiro-Wilk test. For comparisons between two groups, unpaired or paired two-tailed Student’s *t*-tests were applied as indicated in figure legends. Where data failed normality testing, non-parametric tests (Mann–Whitney or Kruskal–Wallis with Dunn’s correction) were used as specified in individual figure legends. For multiple group comparisons, one-way or two-way ANOVA followed by Tukey’s or Sidak’s post-hoc correction for multiple testing was used. Outliers were identified and excluded using the ROUT method (*Q* = 1%). For ordinal clinical variables (e.g., tumor grade and GENIUS score categories), non-parametric Kruskal–Wallis tests were used unless otherwise stated. For intravital imaging, CD8⁺ T cell motility parameters (velocity, turning frequency, displacement) were extracted from time-lapse movies and compared across groups using ANOVA with post-hoc correction. For cytokine arrays and proteomic datasets, differential abundance was determined using multi-sample *t*-tests with false discovery rate (FDR) correction (Benjamini-Hochberg, FDR < 0.05). Volcano plots and hierarchical clustering were generated from log₂-transformed, median-centered, z-scored LFQ values. Significance thresholds for proteomic analyses were set at fold change >2 or <0.5 with adjusted *p* < 0.05. For flow cytometry, frequencies of immune subsets were compared using one-way ANOVA with Tukey’s post-hoc test. Sample sizes were chosen based on prior experience with metastatic breast cancer models and expected effect sizes, rather than formal power calculations. No animals or samples were excluded unless pre-established humane endpoints were reached. Randomization was performed at the level of treatment allocation, and investigators were blinded to group identity during image analysis and quantification. All tests were two-sided, and *p* values < 0.05 were considered statistically significant.

### Data Availability

The proteomic and secretomic mass spectrometry raw datasets have been uploaded to ProteomeXchange, the dataset is currently private and will be publicly available after peer review. Bulk and Single-cell RNA sequencing datasets analyzed in this study are publicly available from the Human Cell Atlas (https://www.humancellatlas.org/) and from McGinnis *et al.* (2021) as cited in the manuscript. All other data supporting the findings of this study are available from the corresponding author upon reasonable request.

## Acknowledgements

This work was supported by the Cancer Research UK Scotland Centre (CTRQQR-2021\100006), the Sir Henry Wellcome Trust grant (221647/Z/20/Z) and the Institute of Genetics & Cancer, University of Edinburgh Langmuir Talent Development Fellowship.

We would like to thank the following service providers at the Institute of Genetics & Cancer, University of Edinburgh; Histology Research Service (Helen Caldwell) for mouse tissue sample preparation, the Flow Cytometry Facility (Elisabeth Freyer) for running samples, the Mass Spectrometry Facility (Alex von Kriegsheim, Chinmayi Pednekar, Agata Markar) for running proteomic samples and the Host and Tumour profiling unit (HTPU) (Alison Munro) for running cytokine array. We thank the Gruss-Lipper Biophotonics Center, the Integrated Imaging Program, the Integrated Imaging Program for Cancer Research and The Evelyn Gruss-Lipper Charitable Foundation.

We thank Damir Vareslija and group for providing a log2 batch-corrected of Perou’s bulk RNA-seq data (PMID: 35082299). We thank Chartinum Chutoe and Alex von Kriegsheim (The University of Edinburgh) for providing 4T1 cells. We thank Marc Vendrell and Lorena Mendive-Tapia (Centre for Inflammation Research, The University of Edinburgh, UK) for providing Apotracker Red to label apoptosis *in vivo*.

## Contributions

N.D.B. conceptualized the study, designed and performed the experiments, analyzed the data and wrote the paper. M.P. and N.D.B. performed the proteomics analysis. N.D.B and F.A.M. analyzed human bulk RNA-seq public datasets and clinical samples with supervision and critical input from R.N. and S.H. and V.B.. C.D. helped with generating the cells and provided critical input into experimental design. B.P., C.M., M.P., E.R.W. and J.C.D. performed experiments. M.W. F.L. and Y.J. provided technical assistance. M.W. and F.L. supported animal studies and helped randomization. X.H. and M.E. provided cells and critical input into the study. A.S., R.N., S.H., M.O. provided critical input into the study. V.B. and D.E. supervised the study, provided guidance in experimental design and data interpretation. N.D.B. and V.B. wrote the paper, with input and revision by D.E.. All authors reviewed and approved the manuscript.

## SUPPLEMENTARY FIGURE LEGENDS

**Supplementary Figure 1:**
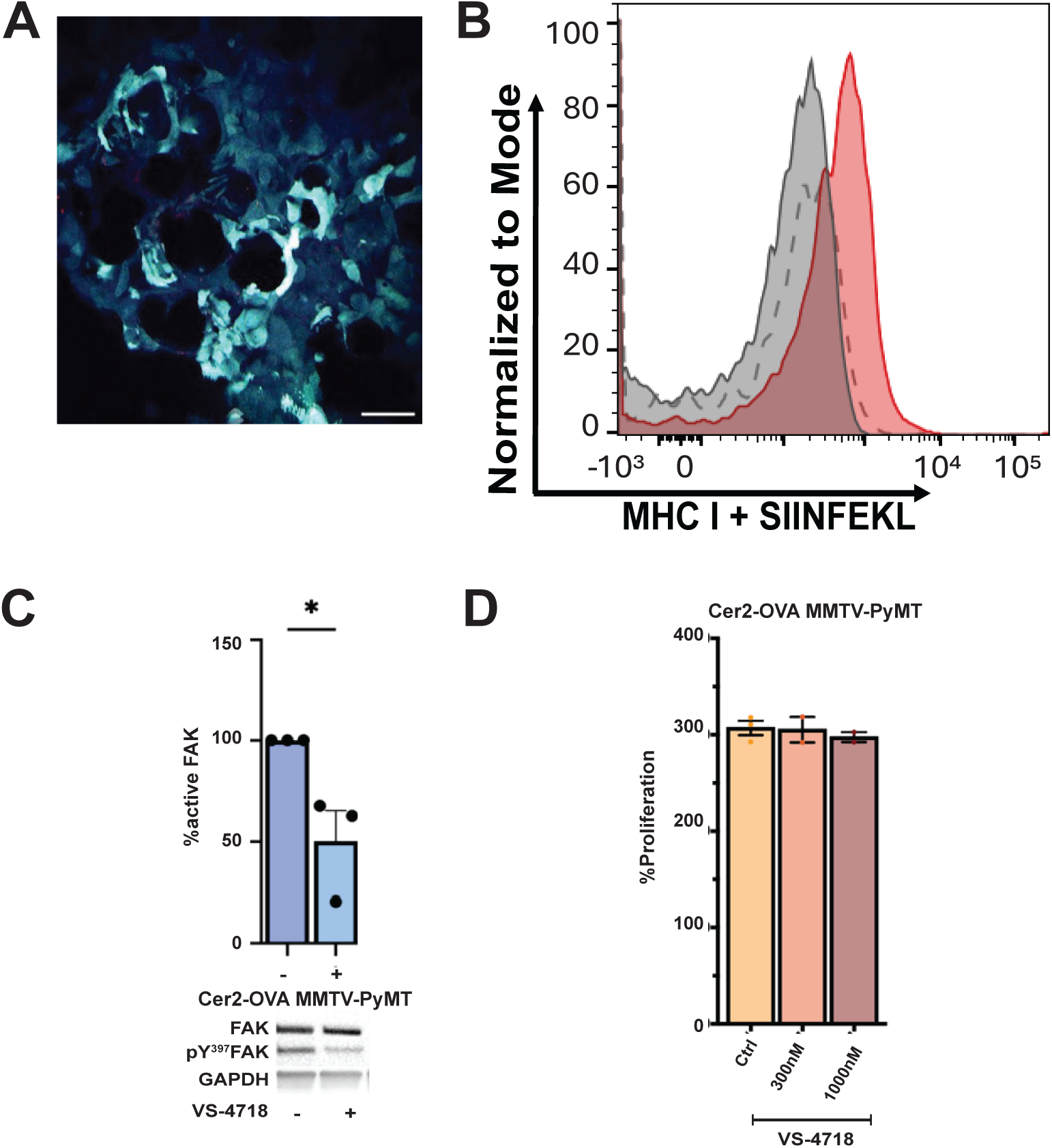
Fluorescent and SIINFEKL-presenting Cer2-OVA MMTV-PyMT cell line. **(A)** Representative 2-photon image of Cerulean2-OVA MMTV-PyMT cells (λ_exc_: 405, 860 nm; λ_em_: 450-510 nm) of *ex vivo* lung slices. **(B)** Surface SIINFEKL-MHC I expression on Cer2-OVA (red) vs. Cer2 (grey) MMTV-PyMT cells analyzed by flow cytometry (representative, n = 3). Scale bar = 50 µm. (**C)** Phospho-FAK levels (Y397) relative to total FAK following 24 h of VS-4718 (1 μM) treatment in Cer2-OVA MMTV-PyMT cells. n ≥ 3 independent experiments, Mean ± SEM; * = p < 0.05. (**D**) No direct effect of VS-4718 on proliferation of breast cancer cell lines in vitro. Quantification of images at 24 h upon addition of VS-4718 at indicated concentrations compared to 0 h expressed as percentual increase of cells (n = 3). Mean ± SEM, Kruskal-Wallis Test, ns < 0.05.

**Supplementary Figure 2:**
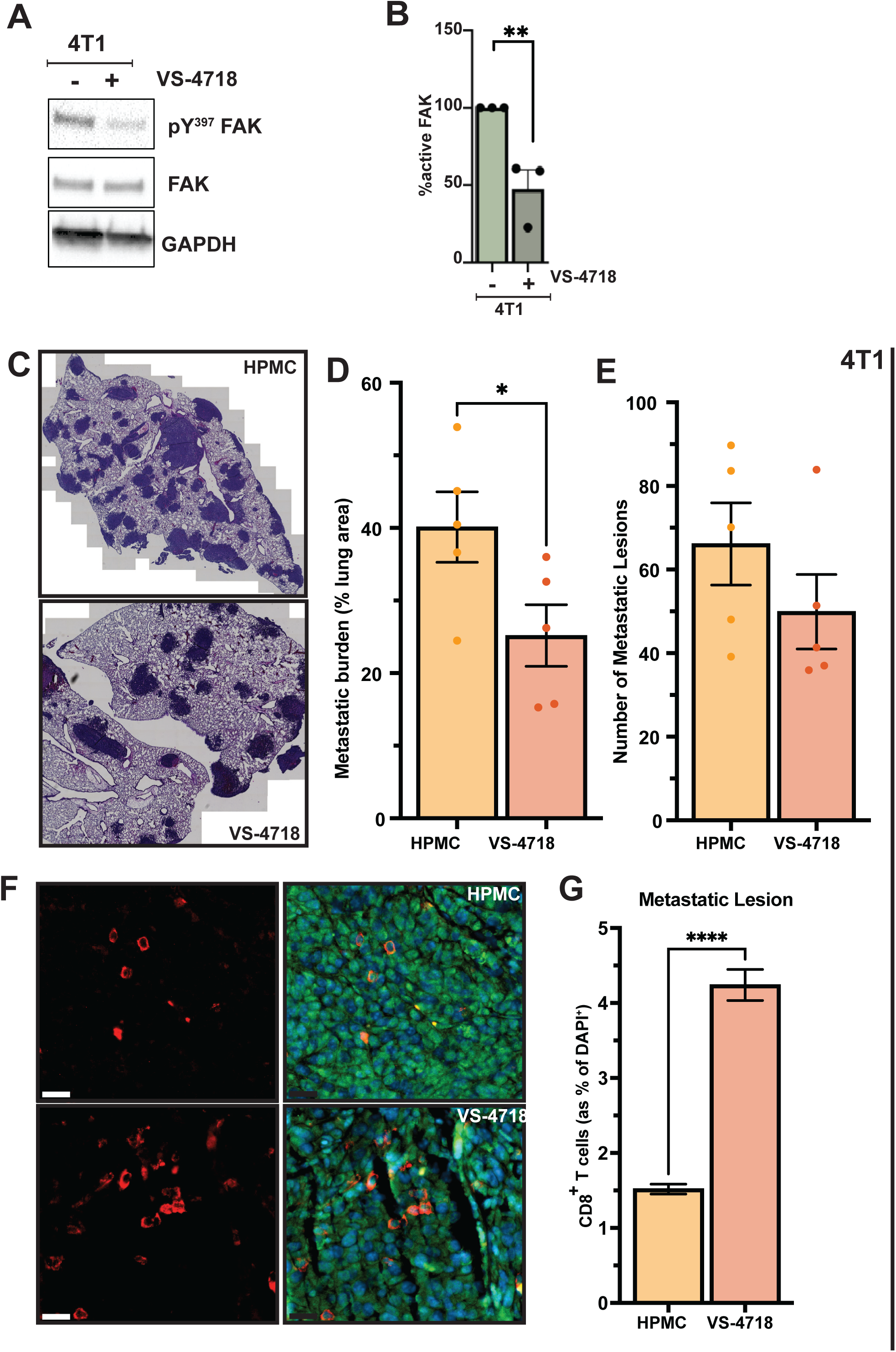
VS-4718 modulates lung metastatic burden and CD8⁺ T cell infiltration in the 4T1 breast cancer model. (A &. **B)** 24 h treatment with 1 µM VS-4718 inhibits FAK phosphorylation at Y397. **(B)** Quantification of phosphorylated versus total FAK (n = 3). **(C-E)** VS-4718 reduces metastatic burden but not lesion number *in vivo*. BALB/c mice were implanted with 4T1 cells and treated with vehicle or VS-4718 for 2 weeks starting day 5 post-implantation. **(C)** Representative H&E images of lung metastases (scale bar, 200 µm). **(D)** Quantification of metastatic burden (% metastatic area/total lung area). **(E)** Average number of metastatic lesions (n = 5 mice). **(F-G)** Representative immunofluorescence images and quantification of CD8⁺ T cells (red) infiltrating metastatic lesions (4T1, anti-p53 = green; nuclei, DAPI = blue; scale bar = 25 µm, n = 5 mice/group). Mean ± SEM; * = p < 0.05, ** = p< 0.01, *** = p < 0.001, **** = p < 0.0001.

**Supplementary Figure 3:**
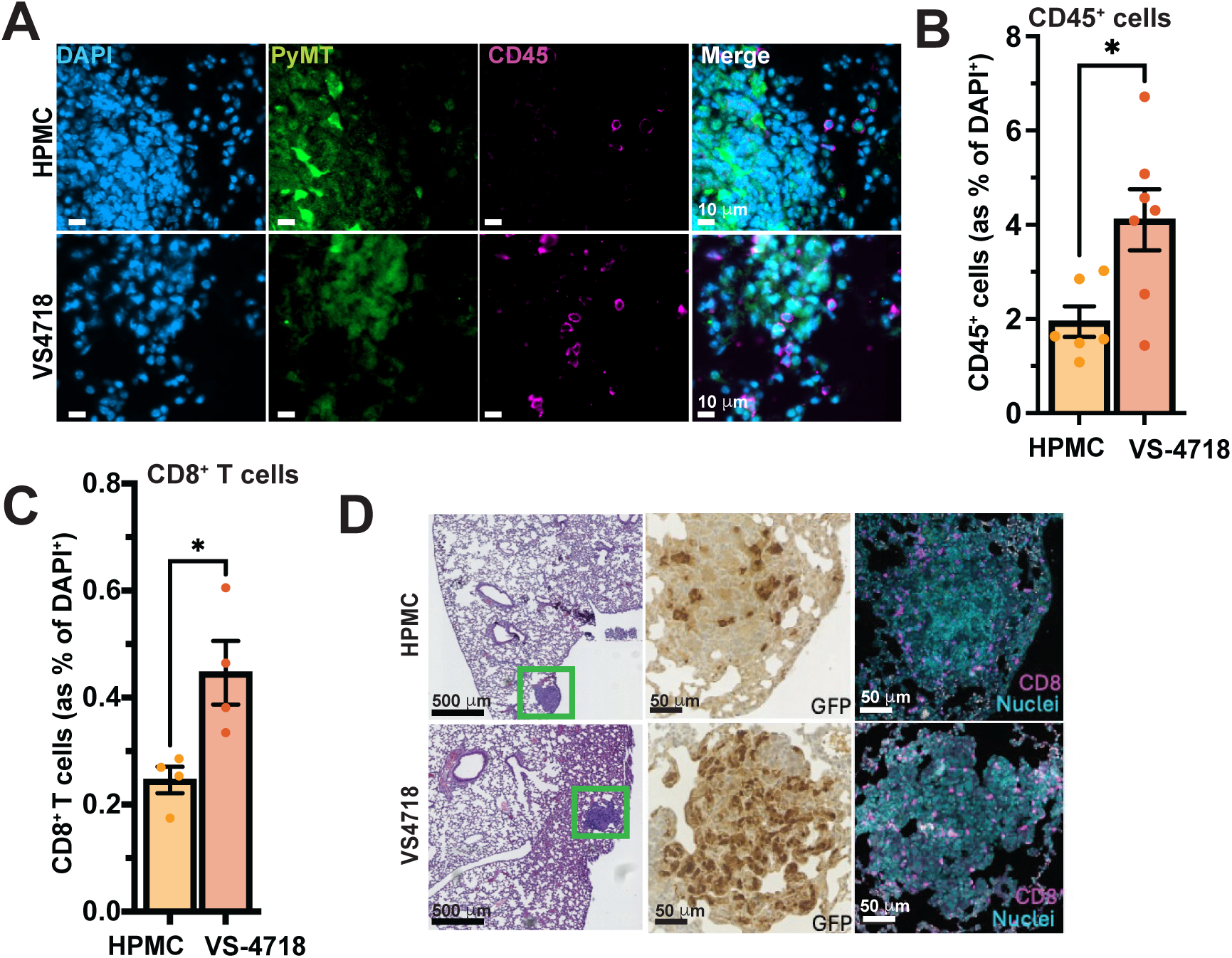
Increased immune cell infiltration in lungs of VS-4718-treated mice. Cer2-OVA MMTV-PyMT cells were injected i.v.. Mice received twice daily treatment with vehicle (0.5% HPMC) or VS-4718 (75 mg/kg) from week 3 to 5 post-implantation. **(A)** Representative immunofluorescence images of CD45⁺ leukocytes and Cer2-OVA cells (anti-GFP, green) in lungs after 2 weeks of treatment. **(B)** Quantification of CD45⁺ cells in whole lungs (n = 7-8 mice/group). **(C)** Increased CD8⁺ T cells in lungs following VS-4718 treatment (n = 4 mice/group). **(D)** Representative H&E staining (left), anti-GFP IHC (middle), and CD8⁺ immunofluorescence (magenta, right) of lung sections (n = 4 mice/group). Mean ± SEM; * = p < 0.05, ** = p < 0.01, *** = p < 0.001, **** = p < 0.0001.

**Supplementary Figure 4:**
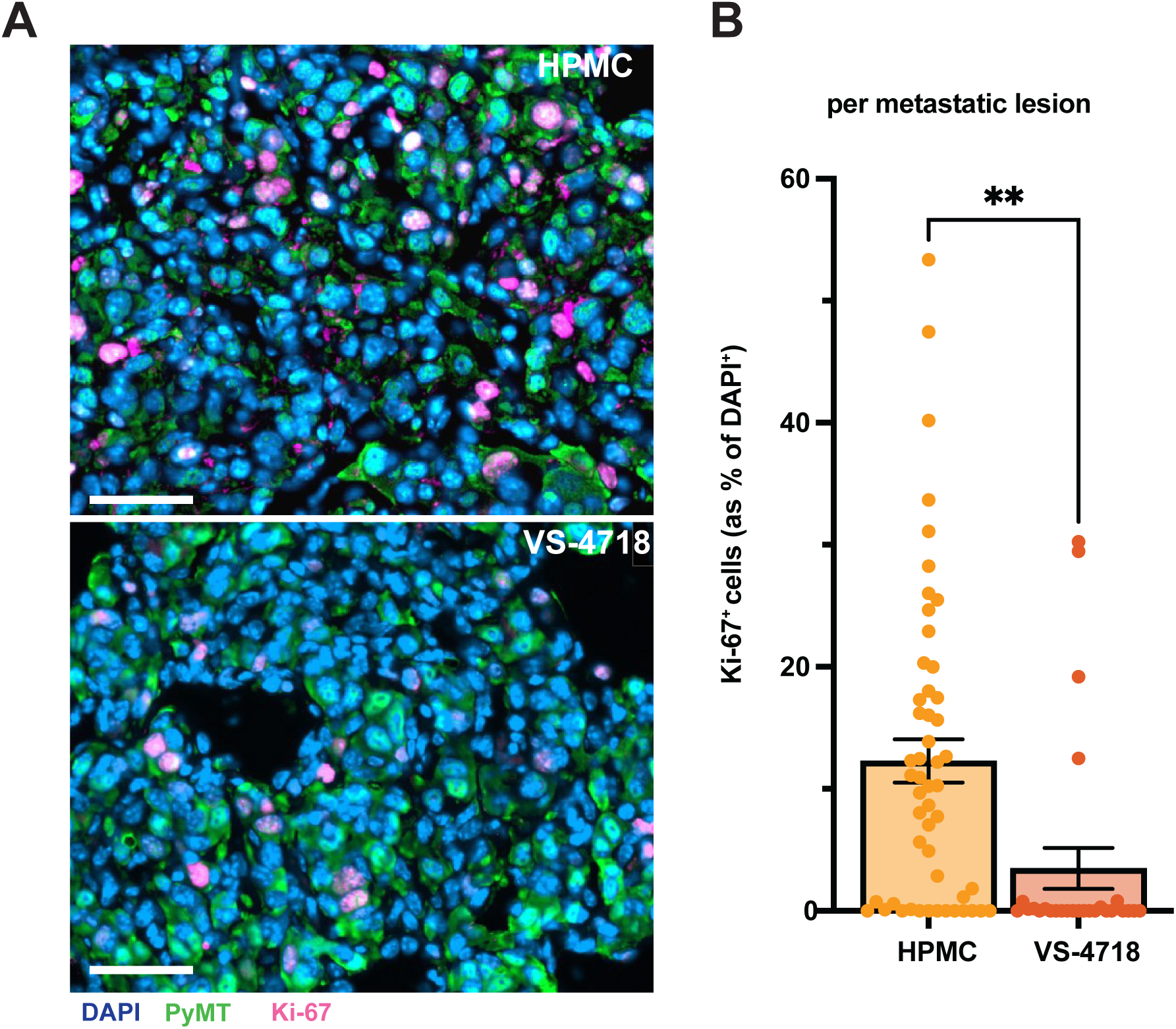
VS-4718-treatment decreases the number of Ki-67^+^ cells in pulmonary metastatic lesions in Nude mice. **(A)** Representative images of Ki-67^+^ cells in metastatic lesions from vehicle (HPMC) and VS-4718 treated mice at 5 weeks. Ki-67 = magenta; Cer2-OVA MMTV-PyMT, anti-GFP = green. Scale bar = 50 μm. **(B)** Quantification of Ki-67^+^ cells per metastatic lesion. Each data point represents one metastatic lesion. n = 5 mice/group. Mean ± SEM; ** = p < 0.01.

**Supplementary Figure 5:**
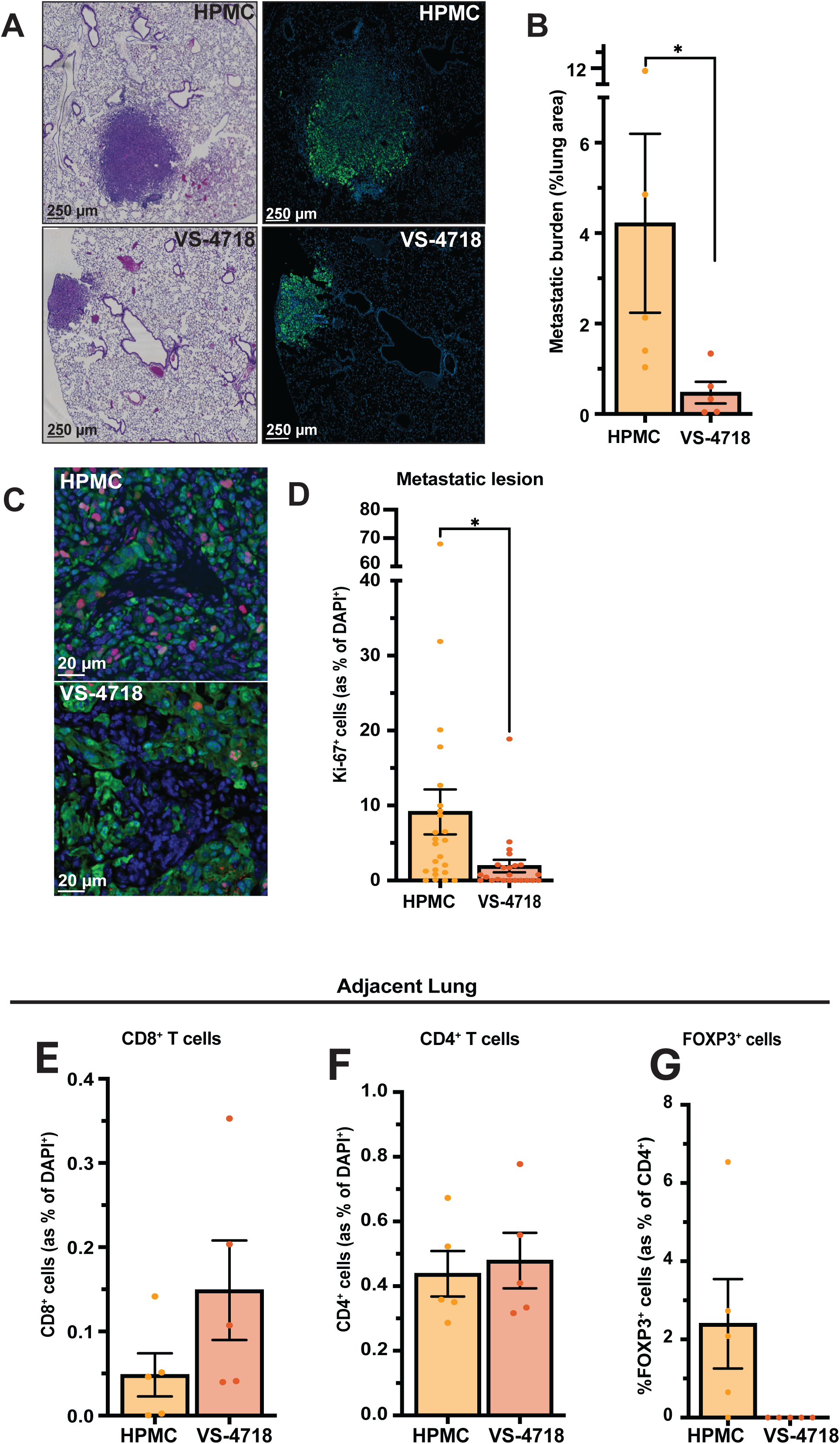
VS-4718 reduces metastatic burden and alters immune cell composition in a pulmonary breast cancer model. Cer2-OVA MMTV-PyMT cells were injected i.v., and mice were treated with vehicle (0.5% HPMC) or 75 mg/kg VS-4718 p.o. from 4-6 weeks post-implantation. **(A)** Representative histology and immunofluorescence of GFP⁺ metastatic lesions (anti-GFP = green; DAPI, nuclei - blue. Scale bar = 250 µm. **(B)** Quantification of metastatic burden (% metastatic area relative to total lung, measured in three sections at 200 µm apart; n = 5 mice/group). Mann-Withney test was performed to compare metastatic burden. **(C)** Representative images of proliferating Ki-67⁺ cells in GFP⁺ metastases. Ki-67 = red; anti-GFP = green; DAPI = blue. Scale bar = 20 µm. **(D)** Quantification of Ki-67⁺ cells in metastatic lesions. n = 5 mice/group. **(E-G)** Quantification of CD8⁺, CD4⁺ (as % of DAPI⁺ cells), and FOXP3⁺ Tregs (as % of CD4⁺ cells) in adjacent lung at 6 weeks (n = 5 mice/group). Mean ± SEM; * = p < 0.05, ** = p < 0.01, *** = p < 0.001, **** = p < 0.0001.

**Supplementary Figure 6:**
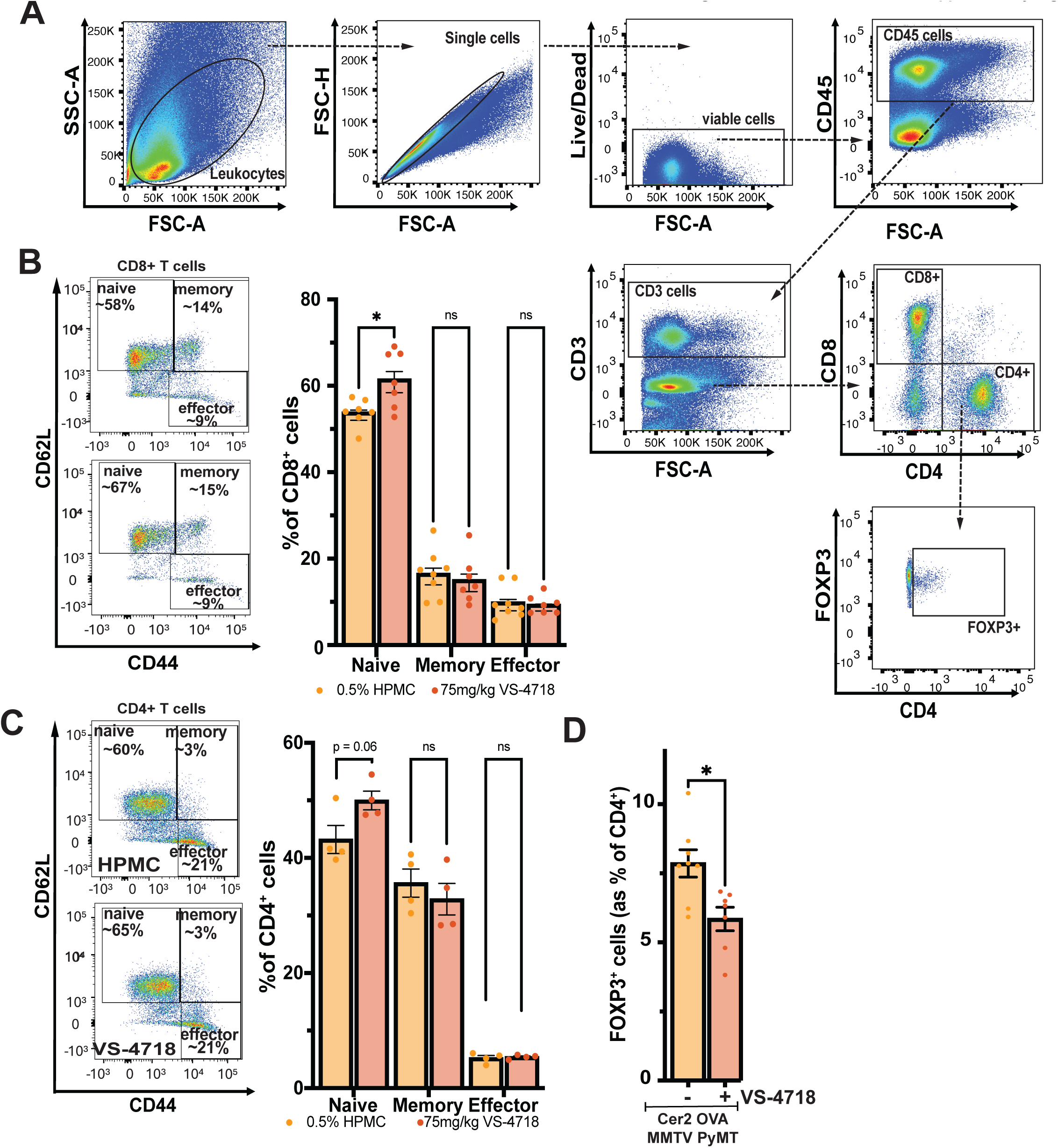
Flow cytometric analysis of CD4 and CD8 T cell subsets 2 weeks post VS-4718 treatment in mice with Cer2-OVA MMTV-PyMT pulmonary metastases. **(A)** Gating schematic for CD4^+^ and CD8^+^ T cells. **(B, Left)** Pre-gating on CD45^+^CD8^+^ cells, naïve CD8^+^ T cells (CD62L^hi^CD44^neg^), effector CD8^+^ T cells (CD62L^lo^CD44^hi^) and memory CD8^+^ T cells (CD62L^hi^CD44^hi^). **(B, Right)** VS-4718 treatment increases the percentage of naïve CD8^+^ T cells in Cer2 OVA PyMT metastatic lungs compared to vehicle. n = 4-8 mice/group. **(C, Left)** Pre-gating on CD45^+^CD4^+^ cells, naïve CD4^+^ T cells (CD62L^hi^CD44^neg^), effector CD4^+^ T cells (CD62L^lo^CD44^hi^) and memory CD4^+^ T cells (CD62L^hi^CD44^hi^). **(C, Right)** VS-4718 treatment increases the percentage of naïve CD4^+^ T cells in Cer2 OVA PyMT pulmonary metastases compared to vehicle. n = 5 mice/group. **(D)** VS-4718 treatment decreases the percentage of regulatory T cells in Cer2 OVA PyMT pulmonary metastases compared to vehicle. n = 4-8 mice/group. Mean ± SEM; * = p < 0.05, ** = p < 0.01, *** = p < 0.001, **** = p < 0.0001.

**Supplementary Figure 7:**
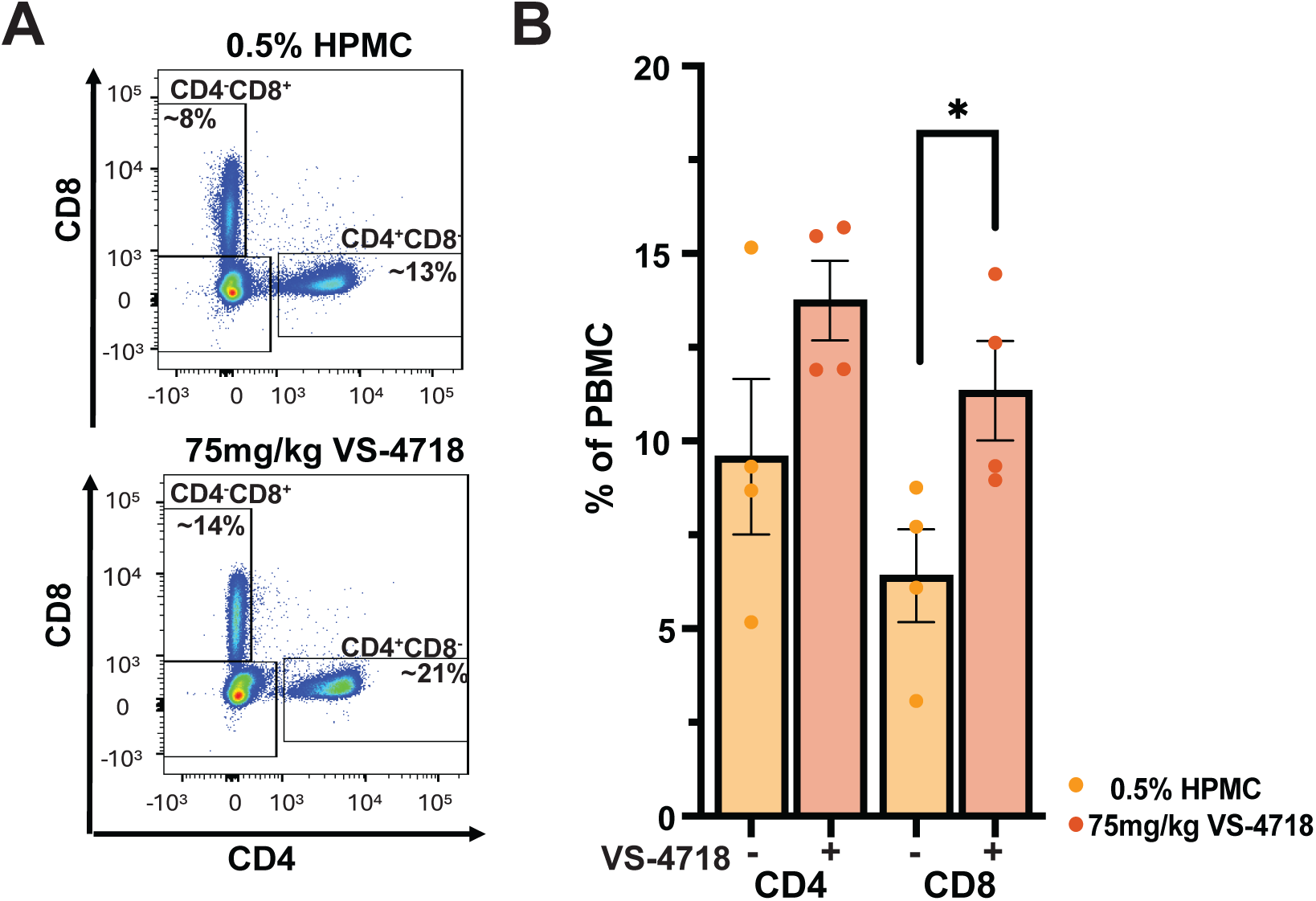
VS-4718 increases circulating CD4⁺ and CD8⁺ T cells. Mice bearing Cer2-OVA MMTV-PyMT tumors were treated twice daily with vehicle (0.5% HPMC) or 75 mg/kg VS- 4718 p.o. for 2 weeks starting 3 weeks post i.v. injection of 1x10⁶ Cer2-OVA MMTV-PyMT cells. n = 4 mice/group. **(A)** Representative flow cytometry pseudo-color plots of CD4⁺ and CD8⁺ T cells in blood (pre-gated on single viable cells). **(B)** Quantification of circulating CD4⁺ and CD8⁺ T cells after 2 weeks of treatment. n = 4 mice/group. Mean ± SEM. *\* =* p < 0.05.

**Supplementary Figure 8:**
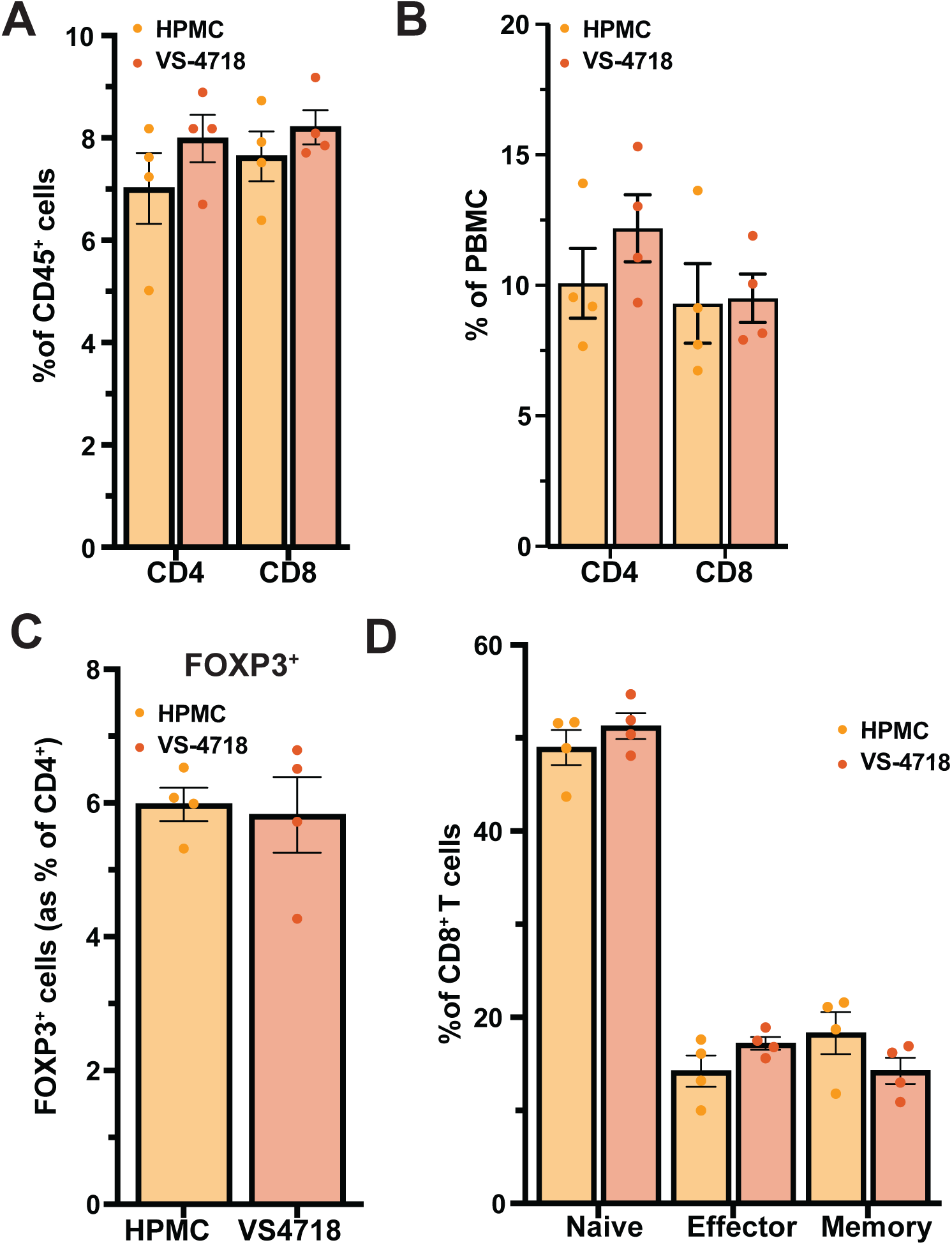
VS-4718 does not alter the immune cell landscape in non-tumor bearing mice. 4 weeks post-PBS implantation, mice were treated with either vehicle (0.5% HPMC) or 75 mg/kg VS-4718 *p.o.* for 2 weeks. **(A)** Quantification of CD4⁺ and CD8⁺ T cells after 2 weeks of treatment in the lung with vehicle control (HPMC) or VS-4718. n = 4 mice/group. Mean ± SEM. **(B)** Quantification of circulating CD4⁺ and CD8⁺ T cells in the blood after 2 weeks with vehicle control (HPMC) or VS-4718 treatment. n = 4 mice/group. Mean ± SEM. **(C)** Compared to vehicle, VS-4718 treatment does not alter the percentage of regulatory T cells in the naïve lung. n = 4 mice/group. Mean ± SEM. **(D)** Compared to vehicle, VS-4718 treatment does not alter the percentage of CD8 T cell subsets in the naïve lung. n = 4 mice/group. Mean ± SEM.

**Supplementary Figure 9:**
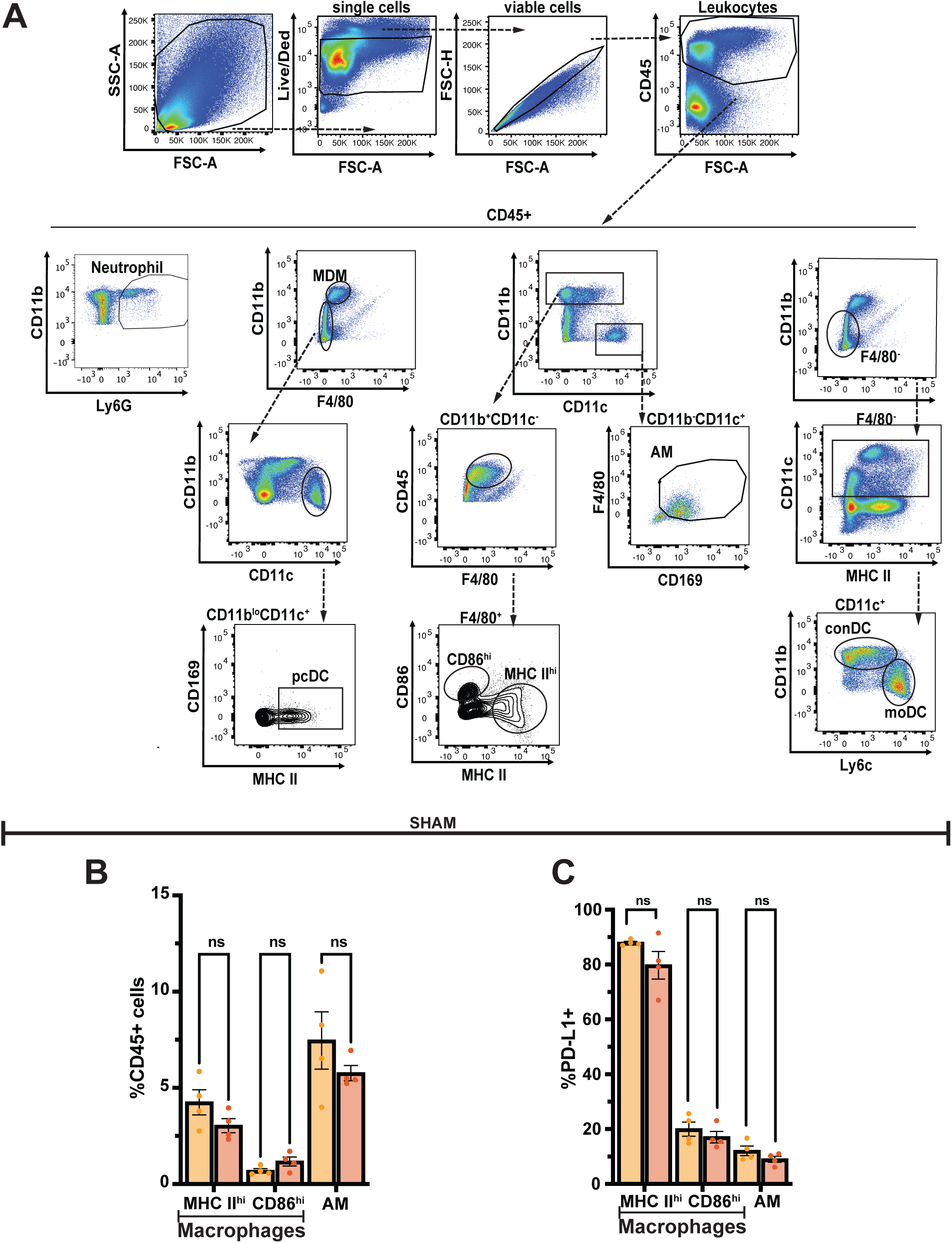
VS-4718 treatment does not influence myeloid subsets in the lung. **(A)** Gating of myeloid populations *in vivo*. **(B)** Percentage bar chart of myeloid populations in the lungs of naïve mice at 2 weeks treatment with vehicle control (HPMC) or VS-4718. Alveolar macrophages (AM) = CD45^+^CD11b^-^CD11c^+^F4/80^+^CD169^+^); MHC II^hi^ macrophages = CD45^+^CD11b^+^CD11c^-^ F4/80^+^CD86^lo^MHC II^+^; CD86^hi^ macrophages = CD45^+^CD11b^+^F4/80^+^CD11c^-^CD86^hi^MHC II^lo^. 4 mice/group. Numbers are percentage of CD45^+^ cells. 4 mice/group. **(C)** Percentage of PD-L1^+^ cells in CD86^hi^ macrophages, MHC II^hi^ macrophages and AM in naïve lungs after 2 weeks of treatment with 0.5% HPMC or 75 mg/kg VS-4718. 4 mice/group. Mean ± SEM; * = p < 0.05, ** = p < 0.01, *** = p < 0.001, **** = p < 0.0001.

**Supplementary Figure 10:**
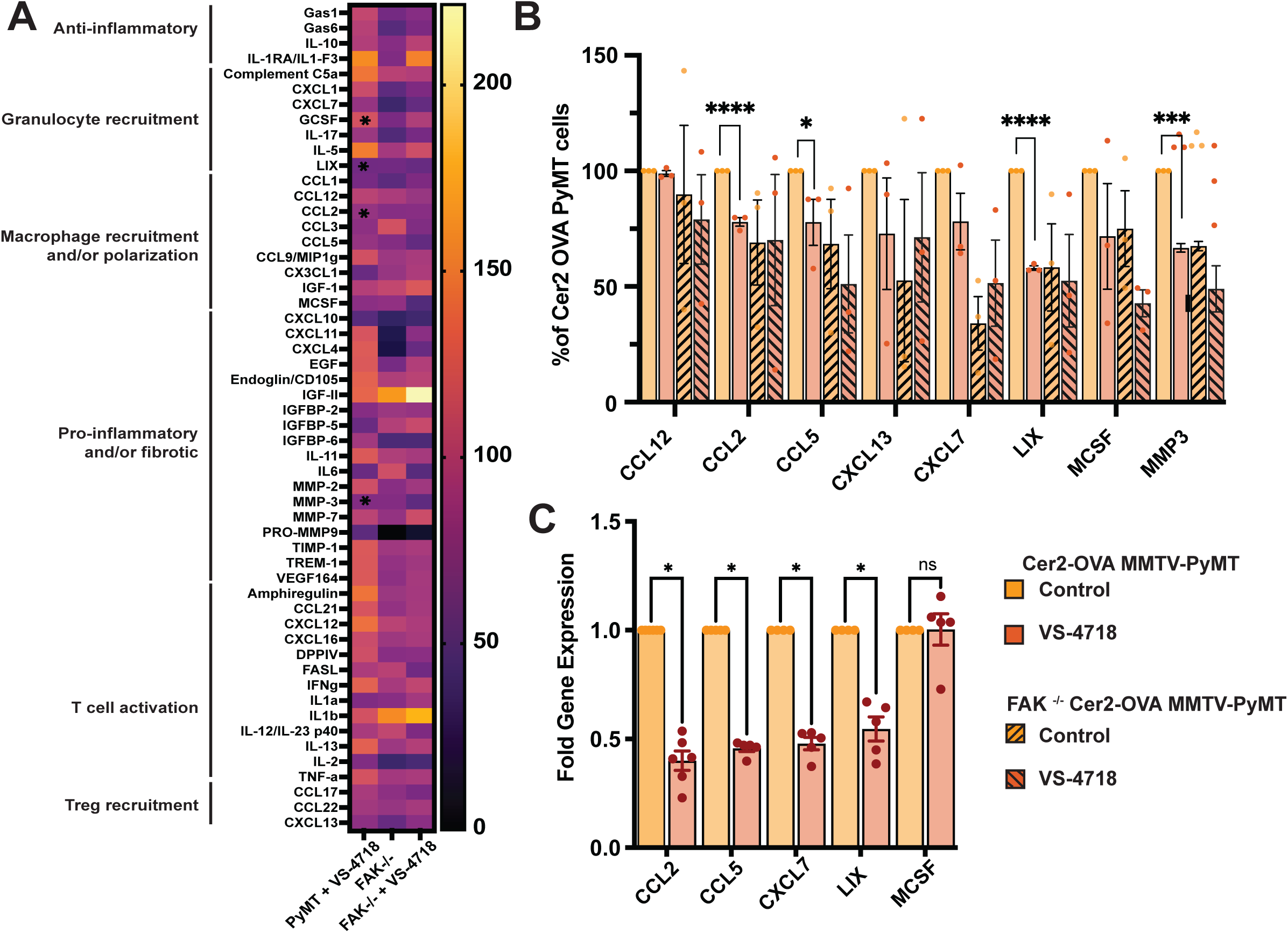
VS-4718 modulates release of cytokines and chemokines by Cer2 OVA MMTV PyMT cells involved in the modulation of the metastatic immune landscape. **(A)** Heatmap of cytokine and chemokine expression in supernatant of control and VS-4718 treated Cer2 OVA MMTV PyMT cells and FAK^-/-^ Cer2 OVA PyMT cells. Cytokine array was performed on supernatant of cells treated for 48h with 1 μM VS-4718. n = 3 independent biological replicates per group. Data is normalized to protein content and displayed as percentage compared to untreated Cer2 OVA MMTV PyMT group. **(B)** Bar graph of significant cytokine and chemokine changes upon VS-4718 treatment with Cer2 OVA MMTV PyMT cells and FAK-/- Cer2 OVA PyMT cells. 3 independent biological replicate per group. **(C)** Bar graph of fold change in gene expression 3 h upon treatment with VS-4718 compared to untreated control. (n = 3-6). Mean ± SEM; * = p < 0.05, ** = p < 0.01, *** = p < 0.001, **** = p < 0.0001.

**Supplementary Figure 11:**
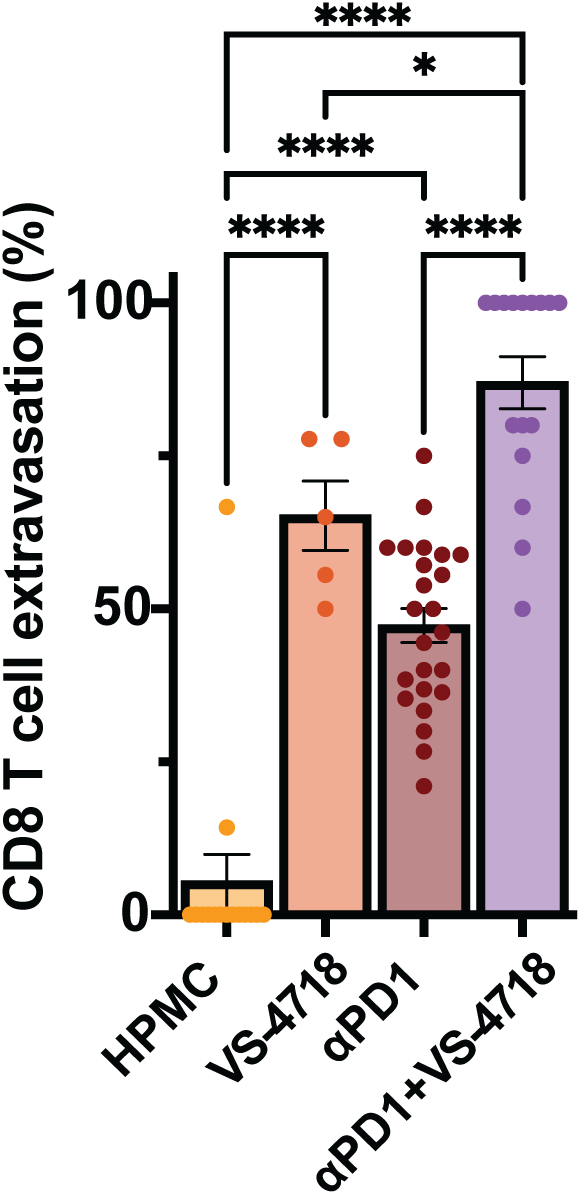
FAK inhibition alone and in combination with anti-PD1 drives CD8^+^ T cell extravasation. Percentage of extravasated cells CD8^+^ T cells. One-way ANOVA. Mean ± SEM; * = p < 0.05, ** = p < 0.01, *** = p < 0.001, **** = p < 0.0001.

**Supplementary Figure 12:**
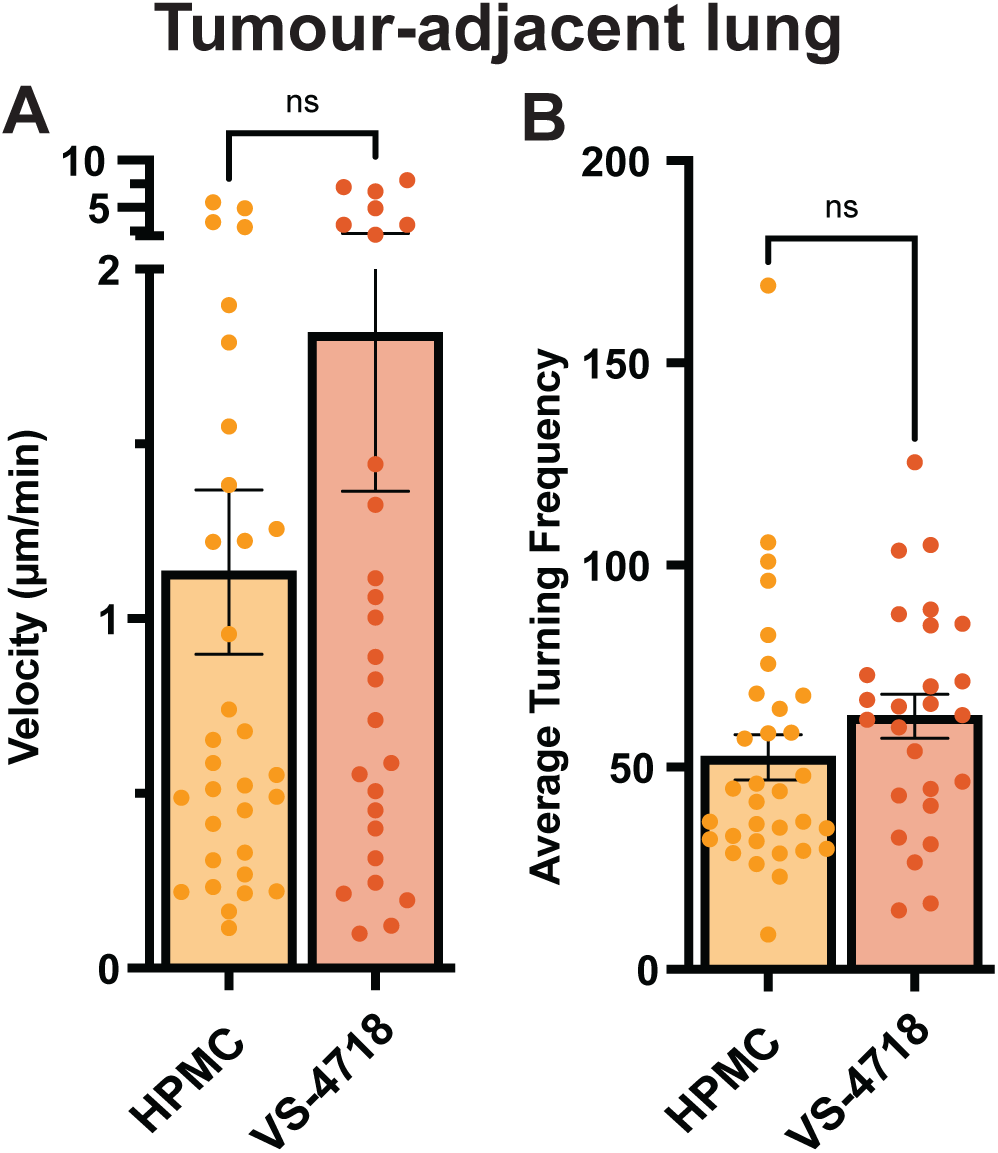
CD8⁺ T cell motility in tumor-adjacent lung tissue. Activated OT-1 CD8⁺ T cells were tracked in precision-cut lung slices from tumor-adjacent regions, and single-cell trajectories were extracted from time-lapse imaging. Quantification shows (A) average velocity and (B) average turning frequency of CD8⁺ T cells. Mean ± SEM; ns = p > 0.05. Unpaired two-tailed t-test.

**Supplementary Figure 13:**
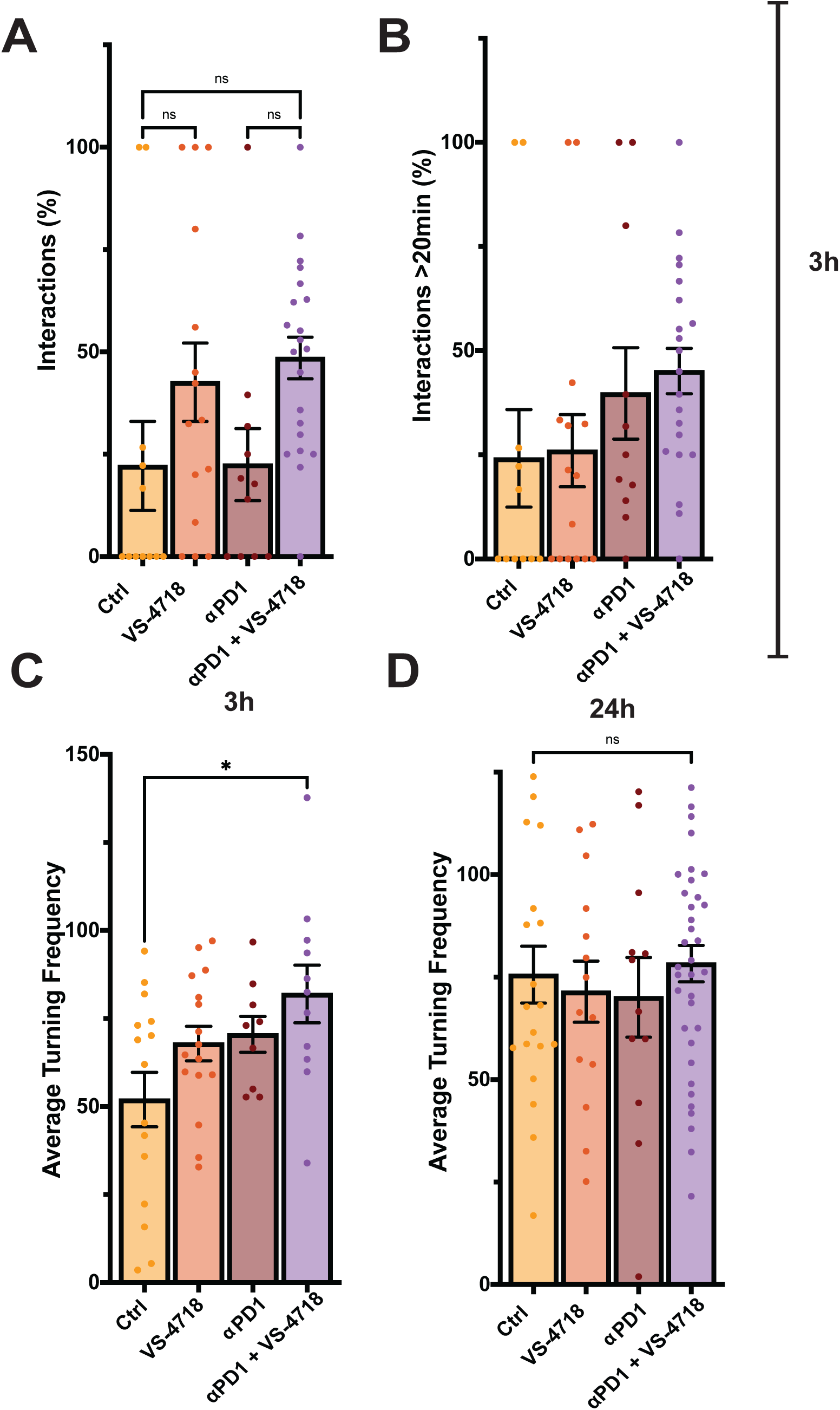
Effect of 3 h treatment on CD8^+^ T cell dynamics in *ex vivo* lung slices. Lung slices (300 μm) from C57BL/6 mice bearing Cer2-OVA MMTV-PyMT metastases (5 weeks post i.v. injection), were pre-treated *ex vivo* for 24 h with vehicle (0.01% DMSO), VS-4718 (300 nM), anti-PD-1 (10 μg/mL), or their combination. CMFDA labelled OT-1 CD8⁺ T cells were added prior to time-lapse multiphoton imaging. Quantification of CD8⁺ T cell-tumor cell dynamics after 3 h of treatment showing: **(A)** interaction frequency, **(B)** stable tethering >20 min, and **(C)** average turning frequency. **(D)** Quantification of average turning frequency after 24 h. n ≥ 3 mice/group using ≥3 slices. Mean ± SEM; * = p < 0.05. One-way ANOVA for normal distributed and Kruskal-Wallis test and Dunn’s multiple comparison for not normal distributed.

**Supplementary Figure 14:**
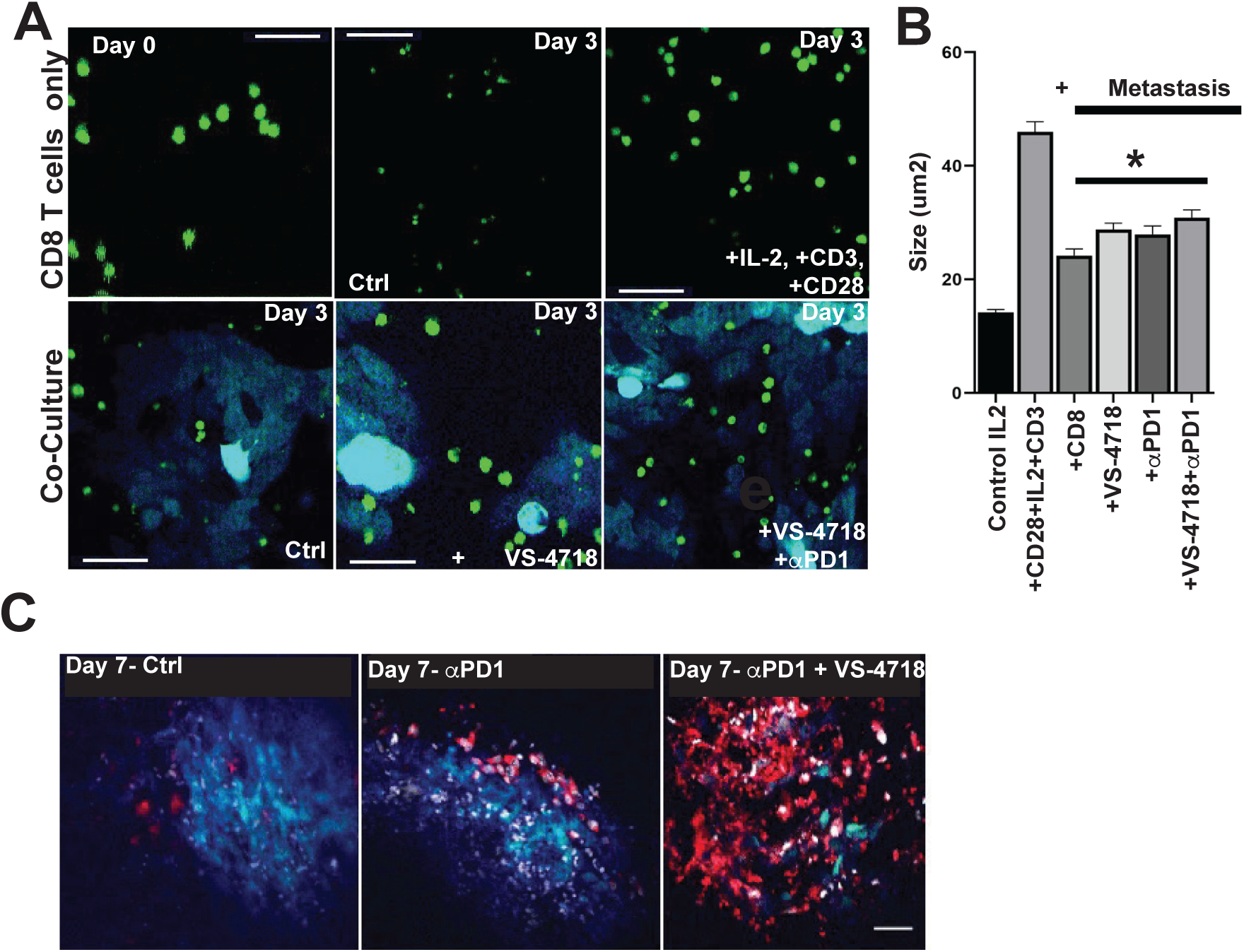
Combination therapy promotes CD8^+^ T cell activity and induces cancer cell cytotoxicity on metastatic lung slices. **(A)** Representative graphs of CD8^+^ T cells alone on agarose or in co-culture with metastatic lung slices. To determine the impact of activation of CD8^+^ T cells cultured alone, cells were treated with either IL-2 (80 IU/mL) alone as a control, or the combination of IL-2 (80 IU/mL), α-CD3 (2 μg/mL), and α-CD28 (5 μg/mL) for 3 days. Cells cultured on lung slices were treated with IL-2 (80 IU/mL) alone or combination of IL-2 (80 IU/mL) and VS-4718 (300 nM), αPD1 (10 μg/mL) or their combination. CD8^+^ T cells = green; Cer2-OVA MMTV-PyMT = cyan. **(B)** Quantification of CD8^+^ T cell size after 3 days culture on agarose or metastatic lung slices. 3 independent experiments per group using 3 fields of view each. Mean ± SEM; * = p < 0.05. **(C)** Representative images of OT-1 CD8^+^ T cells co-cultured on metastatic lung slices for 7-days with control, α-PD1, or combined α-PD1 + VS-4718 treatments. Cer2-OVA MMTV-PyMT = blue; ApoRed (indicating apoptosis) = red; nucleus = white.

**Supplementary Figure 15:**
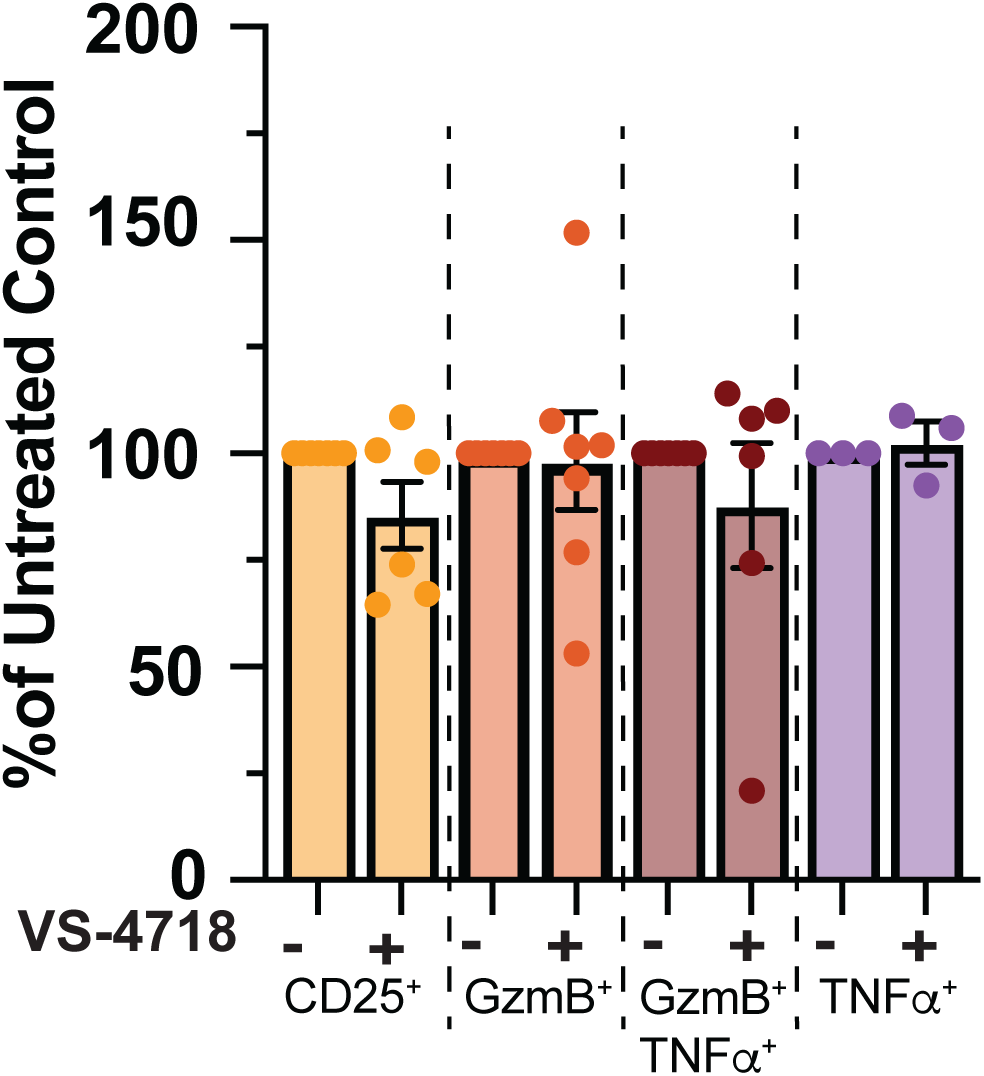
VS-4718 treatment does not affect activation of CD8^+^ T cells *in vitro*. Quantification of the percentage of activity markers relative to untreated control. CD8^+^ T cells were isolated and stimulated for 48 h ± VS-4718. Mean +/- SEM. n = 3-6 independent experiments.

**Supplementary Figure 16:**
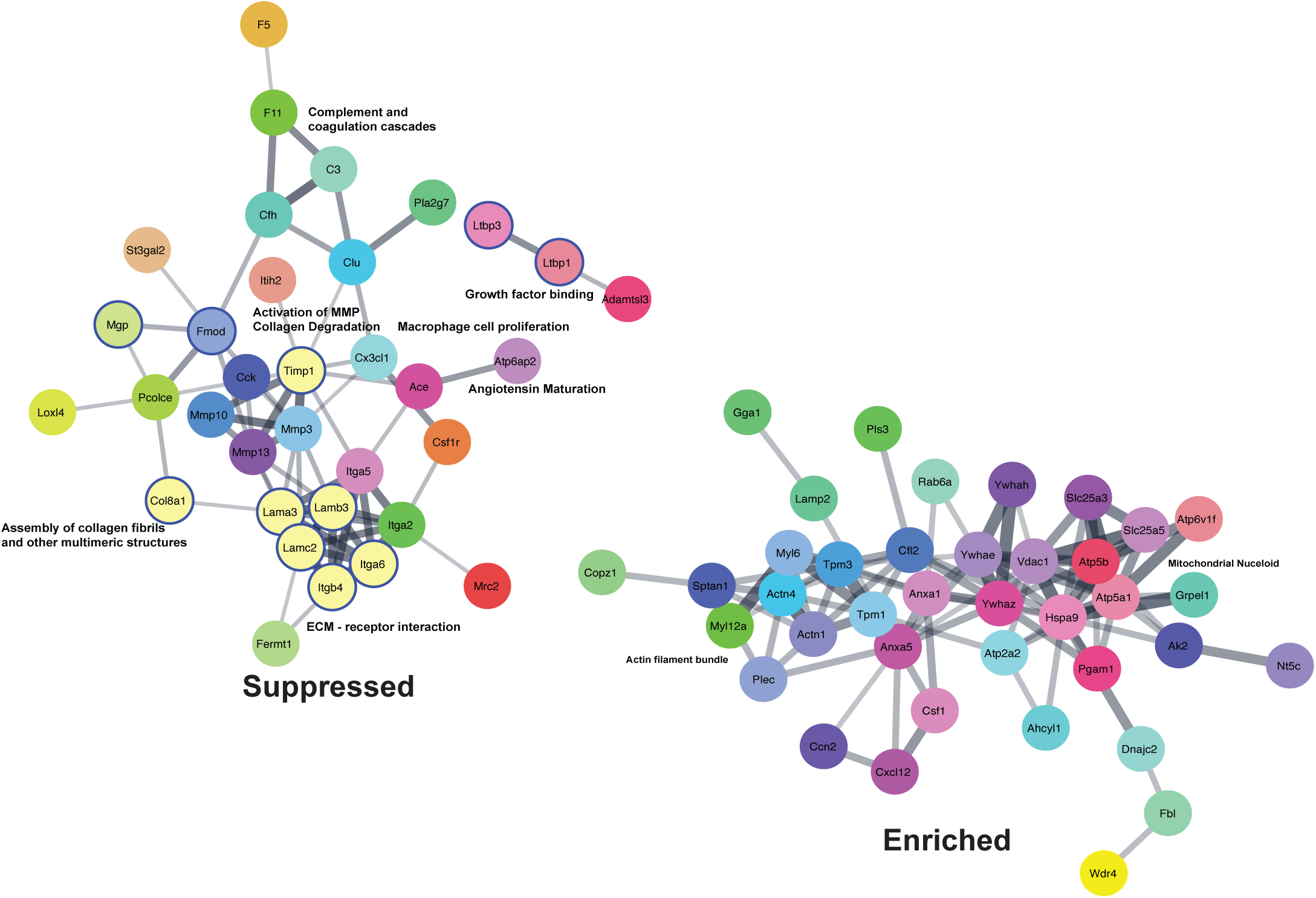
STRING interaction network of significantly altered secreted proteins showing functional clusters visualized in Cytoscape. MCL clustering, inflation parameter+3; |log₂FC| ≥ 0.5; p < 0.05. Pale yellow circles: basement membrane, other colors: random. Blue outline: ECM factors. Edges are weighted by the S score, with edge thickness proportional to the strength of similarity between gene sets.

**Supplementary Figure 17:**
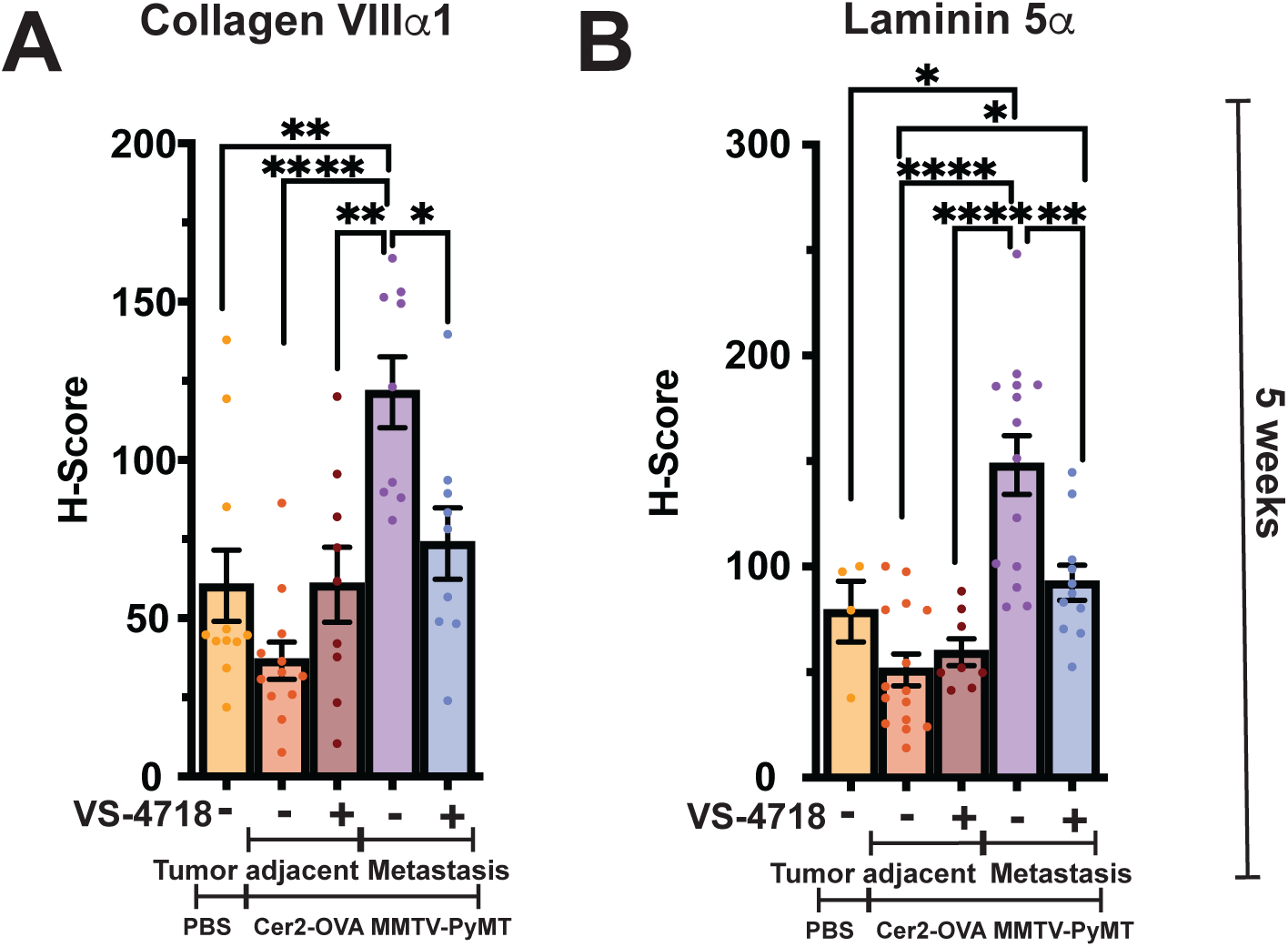
Increased collagen VIIIα1 and laminin 5α in metastatic lesions at 5 weeks post Cer2-OVA MMTV-PyMT injection. Mice carrying Cer2-OVA MMTV-PyMT metastatic lesions or without Cer2-OVA MMTV-PyMT injection were treated for two weeks with either vehicle control (0.5% HPMC) or VS-4718 (75 mg/kg). Quantification (H-score) of concentration of **(A)** Collagen VIIIα1 and **(B)** Laminin 5α in metastatic lesions and tumor adjacent areas at 5 weeks post implantation of Cer2-OVA MMTV-PyMT cells. Each dot represents a metastatic lesion. n = 5 mice/group using whole lung slide scans. One-way ANOVA for normal distributed with Tukey’s multiple comparison test. Mean ± SEM; ** = p < 0.01.

**Supplementary Figure 18:**
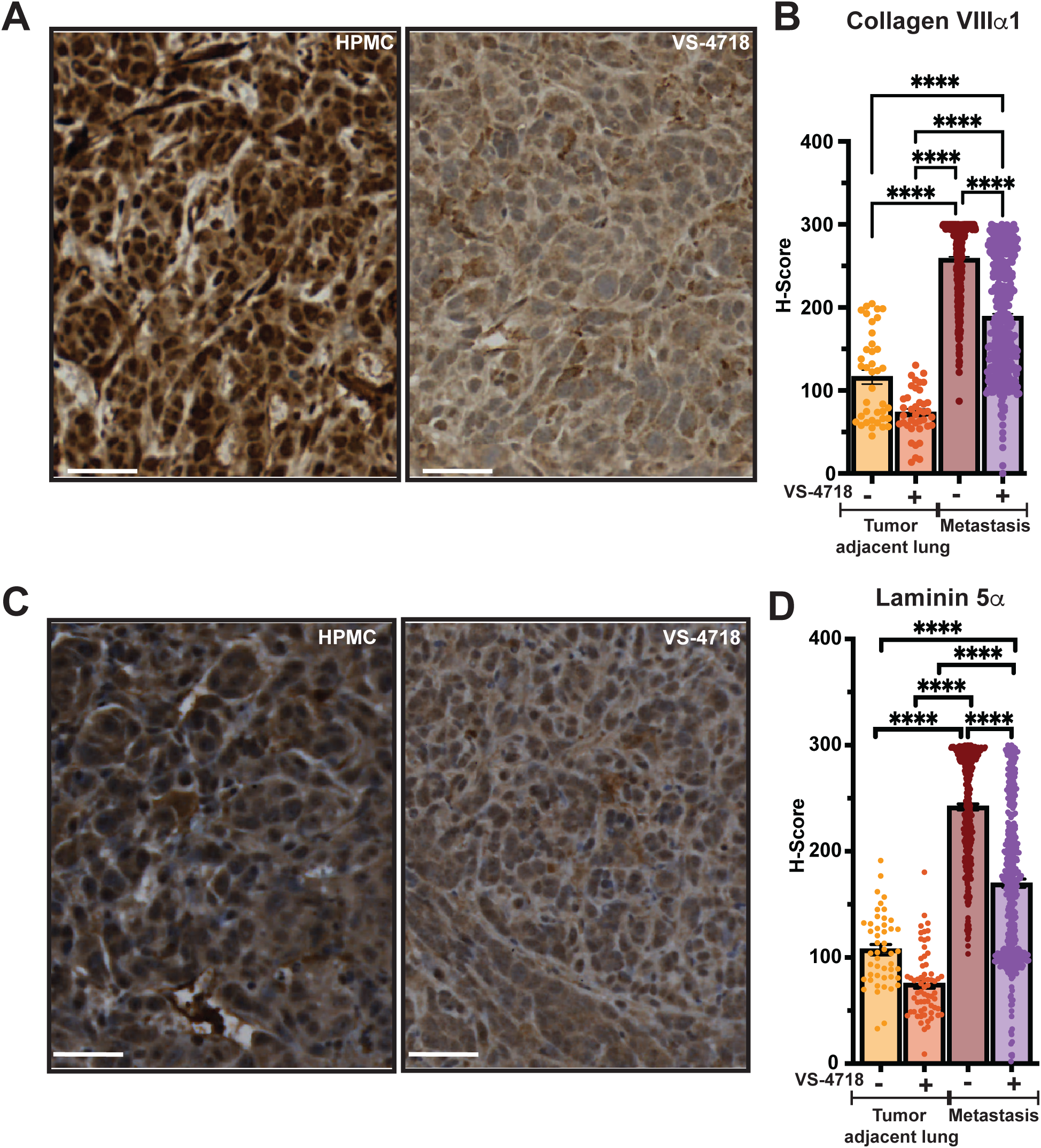
Compared to control treatment, VS-4718-treated mice exhibit decreased concentrations of collagen VIIIα1 and laminin 5 α in pulmonary metastatic lesions from 4T1 cells. **(A)** Representative images of collagen VIIIα1 in metastatic lesions. Scalebar = 50 μm. **(B)** Quantification (H-score) of collagen VIIIα1 in metastatic lesions and tumor adjacent areas. **(C)** Representative images of laminin 5 α in metastatic lesions and tumor adjacent areas. Scalebar = 50 μm. **(D)** Quantification (H-score) of laminin 5 α in metastatic lesions and tumor adjacent areas. Dots represent metastatic lesions. n = 5 mice/group using whole lung slide scans. Kruskal-Wallis test and Dunn’s multiple comparison for not normal distributed. Mean ± SEM; **** = p < 0.0001.

**Supplementary Figure 19:**
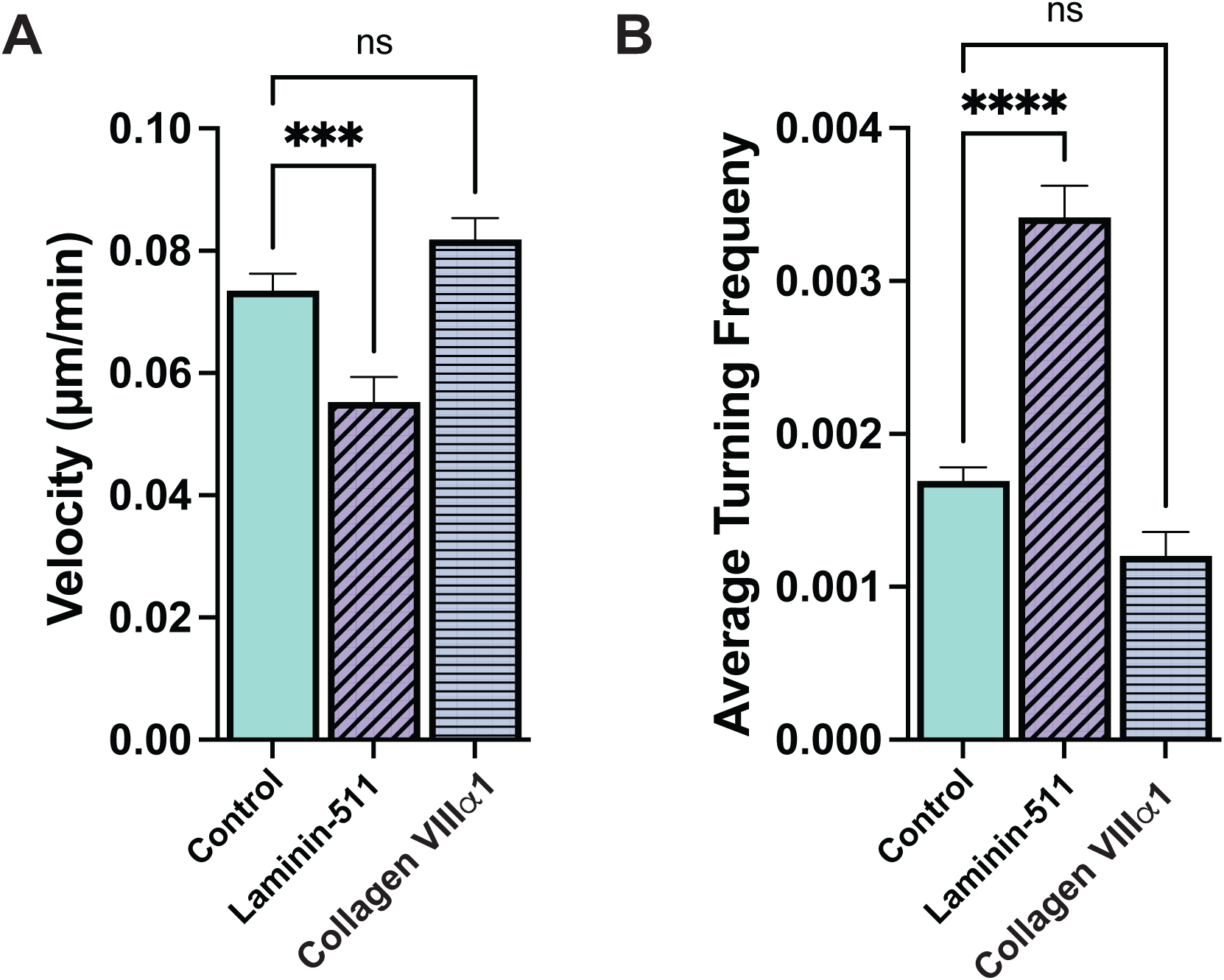
Laminin-511, but not Collagen VIIIα1, alters CD8⁺ T-cell motility. (A) Mean migration velocity and (B) directional change rate/ average turning frequency of activated CD8⁺ T cells migrating on laminin 511 or collagen VIIIα1 (1 µg/cm²). T cells were CellTracker™ Deep Red-labeled and imaged for 6 h at 5-min intervals; tracks with >11 spots were analyzed using TrackMate^89^. Laminin-511 reduced velocity and increased turning frequency, whereas collagen VIIIα1 had no significant effect. n = 3 independent experiments with 3-5 field of views per group. Unpaired t-tests. Mean ± SEM; **** = p < 0.0001.

**Supplementary Figure 20:**
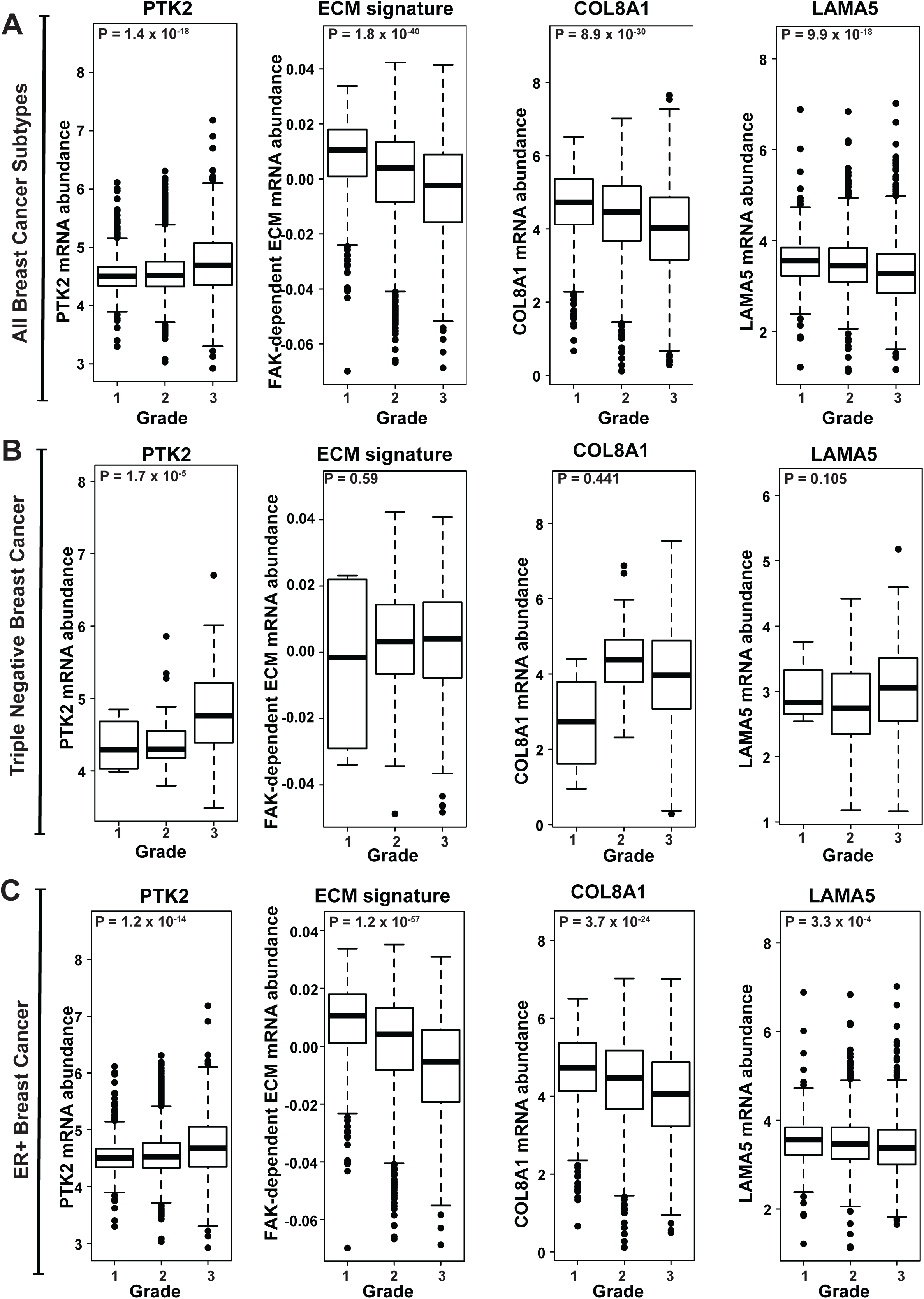
Expression of PTK2, FAK-dependent ECM score, *COL8A1* and *LAMA5* in breast cancer patients of grade 1 to 3. Spearman’s Correlation of Grade correlated with PTK2, FAK-dependent ECM score, COL8A1 and LAMA5 expression in all breast cancer patients (**A**) and separated into triple negative breast cancer (**B**) and ER+ breast cancer (**C**) subtypes of the SCAN-B dataset^62^.

**Supplementary Figure 21:**
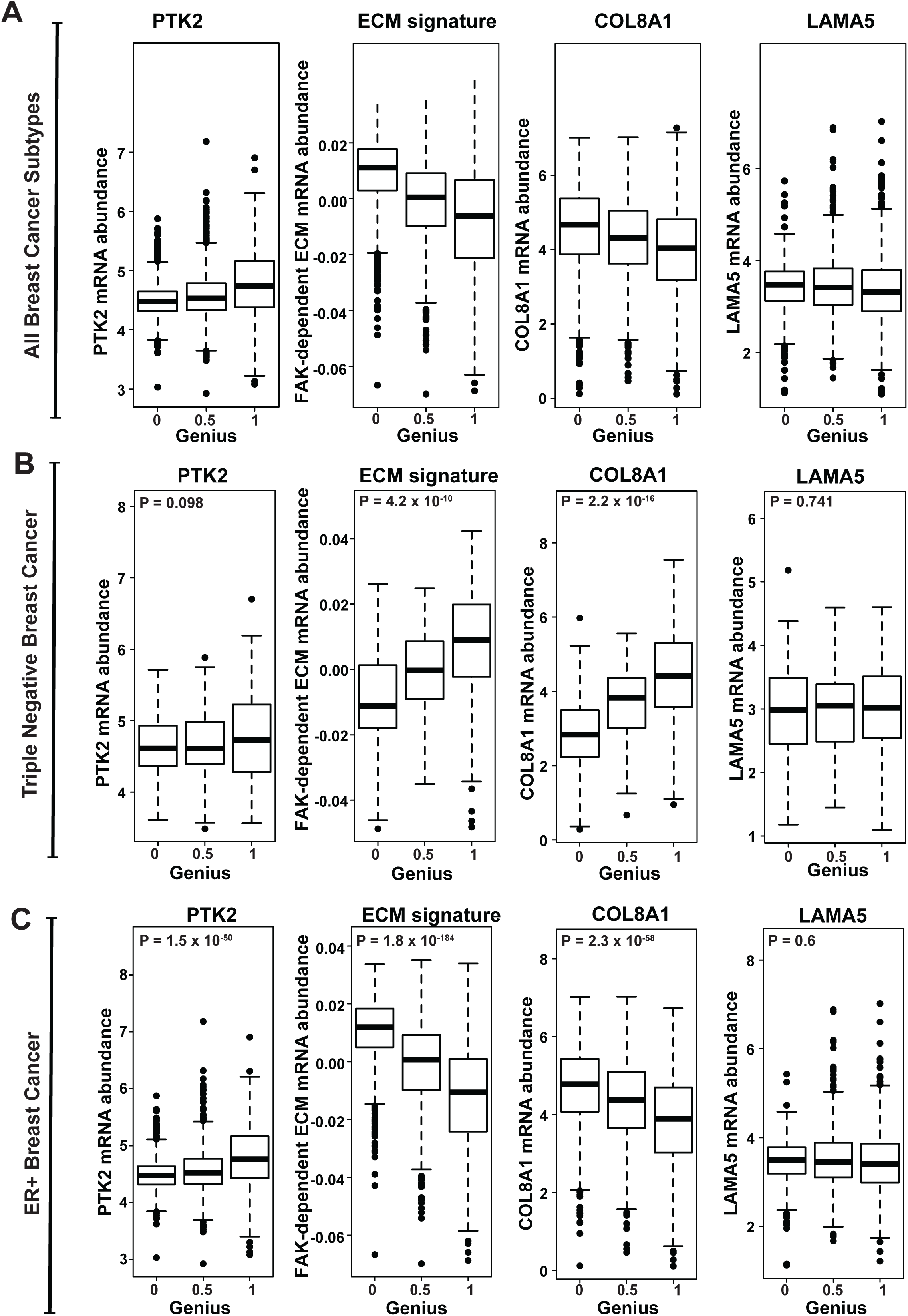
Expression of PTK2, FAK-dependent ECM score, *COL8A1* and *LAMA5* correlated to Genius score. Spearman’s Correlation of Genius score correlated with PTK2, FAK-dependent ECM score, COL8A1 and LAMA5 expression in all breast cancer patients (**A**) and separated into triple negative breast cancer (**B**) and ER+ breast cancer (**C**) subtypes of the SCAN-B dataset^62^.

**Supplementary Figure 22:**
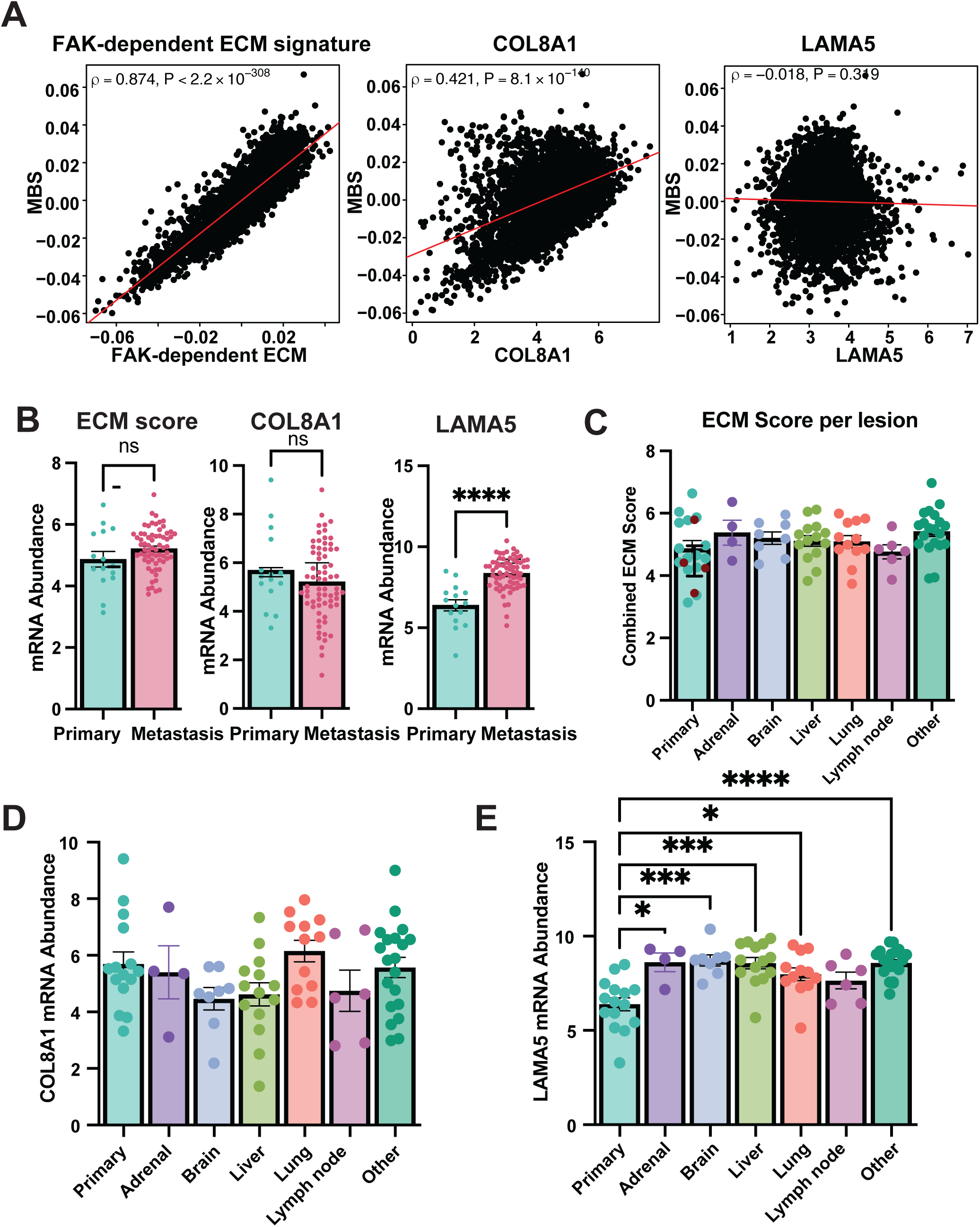
FAK-dependent ECM score, *COL8A1* and *LAMA5* expression in primary site and metastatic lesions. **(A**) Spearman’s Correlation of Metastatic burden signature correlated to FAK-dependent ECM score, COL8A1 and LAMA5 expression in all breast cancer patients of the SCAN-B dataset^62^. (**B**) Analysis of RNAseq data taken from Perou et al. and pre-processed by Damir et al. showing the gene expression levels of FAK-dependent ECM signature (left), *COL8A1* (middle) and *LAMA5* (right) in primary breast tumors and matched metastatic lesions taken from locations indicated. n = 15 primary tumors, n = 69 metastatic lesions. Dots represent expression in individual lesions. Expression of **(C**) FAK-dependent ECM signature, (**D**) COL8A1 and (**E**) LAMA5 expression in primary site and lesions divided into subtypes including adrenal, brain, liver, lung, lymph node and other. Each dot represents a primary or metastatic lesion. Mean ± SEM; *p < 0.05, **p < 0.01, ***p < 0.001, ****p < 0.0001.

**Supplementary Figure 23.**
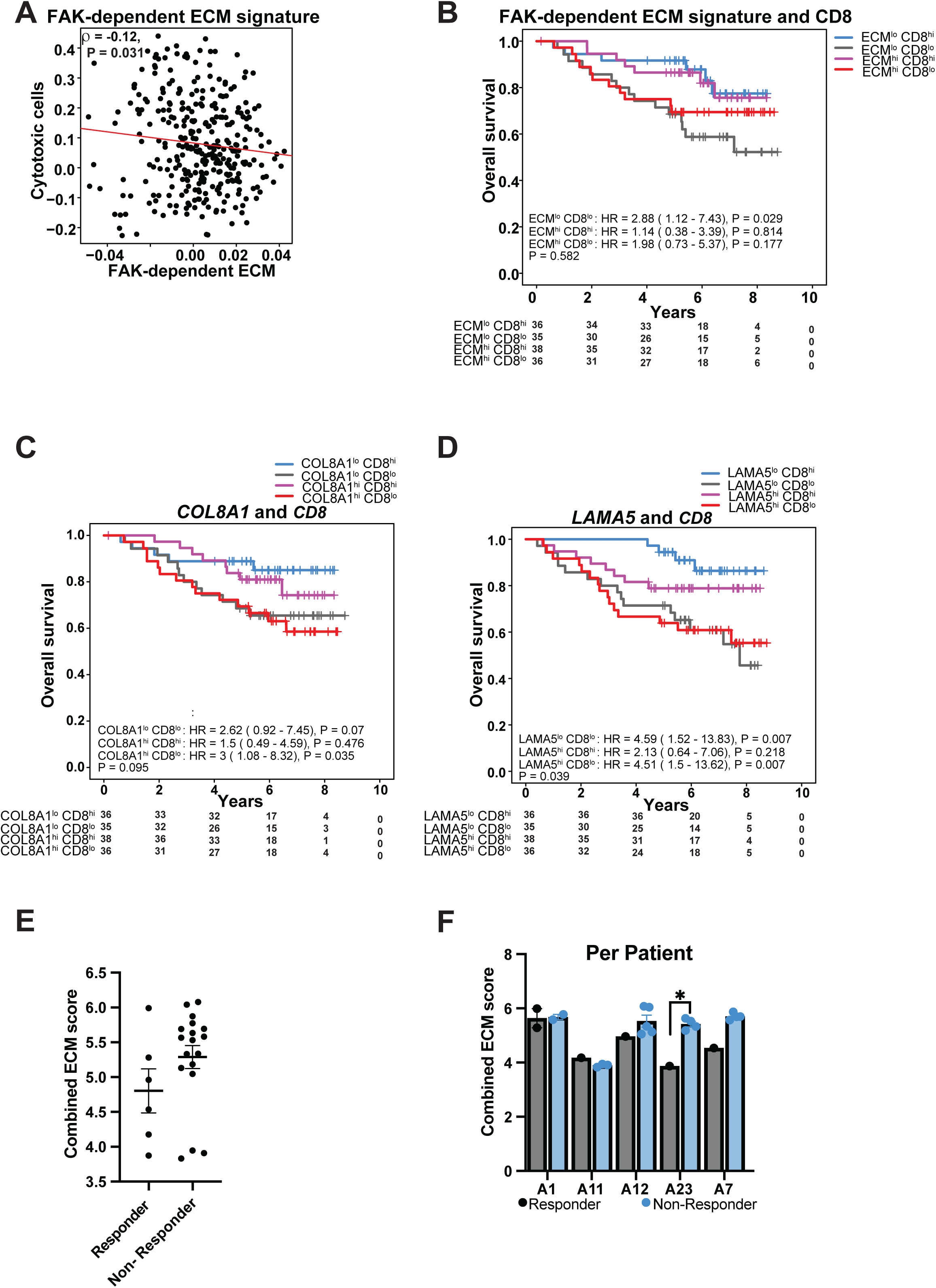
ECM genes correlated to cytotoxic cells and survival of breast cancer patients. **(A**) Spearman’s Correlation of cytotoxic CD8^+^ T cells correlated to FAK-dependent ECM score in in triple negative breast cancer patients of the SCAN-B dataset. (**B-D**) Kaplan Meier Plots of overall survival separated by FAK-dependent ECM score (**B**), COL8A1 (**C**) and LAMA5 (**D**) in triple negative breast cancer patients of SCAN-B dataset. Patients are separated into four group according to expression of gene of interest (GOI) low (lo) or high (hi) and CD8^+^ T cells. Groups were:i) GOI^hi^ CD8^lo^ (red), GOI^lo^ CD8^hi^ (blue), GOI^hi^ CD8^hi^ (purple), GOI^lo^ CD8^lo^ (grey). (**E)** ECM score in responders and non-responders to breast cancer therapy in Perou’s dataset. (**F**) ECM score in responders and non-responders to breast cancer therapy broken out by patient. Detailed information on treatments can be found in Garcia-Recio et al. ^65^. Mean ± SEM; *p < 0.05, **p < 0.01, ***p < 0.001, ****p < 0.0001.

**Supplementary Table 1:** List of quantitative proteomics in matched cell lysate and supernatant. Unsupervised hierarchical clustering of cellular and secreted proteins from Cer2-OVA MMTV-PyMT cells treated +/- VS-4718 for 24 h and 48 h, respectively, revealing three major clusters (C1-C3) based on Euclidean distance. P values were adjusted using Benjamini–Hochberg FDR.

| UniProt Accession | Gene Symbol | Protein Name | Cluster | Regulation pattern |
| --- | --- | --- | --- | --- |
| P56480 | ATP5F1B | ATP synthase subunit beta, mitochondrial | C1 | Down in cells, up in supernatant |
| P46638 | RAB11B | Ras-related protein Rab-11B | C1 | Down in cells, up in supernatant |
| Q9JKF1 | IQGAP1 | Ras GTPase-activating-like protein IQGAP1 | C1 | Down in cells, up in supernatant |
| P07356 | ANXA2 | Annexin A2 | C1 | Down in cells, up in supernatant |
| Q62261-2 | SPTBN1 | Spectrin beta chain, non-erythrocytic 1 | C1 | Down in cells, up in supernatant |
| P16546-2 | SPTAN1 | Spectrin alpha chain, non-erythrocytic 1 | C1 | Down in cells, up in supernatant |
| A0A087WRX8 | SRRM2 | Serine/arginine repetitive matrix protein 2 | C2 | Up in cells and supernatant |
| Q9R0H5 | KRT71 | Keratin, type II cytoskeletal 71 | C2 | Up in cells and supernatant |
| Q8VDD5 | MYH9 | Myosin-9 | C2 | Up in cells and supernatant |
| E9QA15 | CALD1 | Caldesmon 1 | C2 | Up in cells and supernatant |
| Q80X50-5 | UBAP2L | Ubiquitin-associated protein 2-like | C2 | Up in cells and supernatant |
| A0A7N9VR94 | AHNAK2 | AHNAK nucleoprotein 2 | C2 | Up in cells and supernatant |
| Q7TQH0-2 | ATXN2L | Ataxin-2-like protein | C2 | Up in cells and supernatant |
| Q61140-2 | BCAR1 | Breast cancer anti-estrogen resistance protein 1 | C2 | Up in cells and supernatant |
| Q9QZQ1-2 | AFDN | Afadin | C2 | Up in cells and supernatant |
| Q9ER72 | CARS1 | Cysteine-tRNA ligase, cytoplasmic | C2 | Up in cells and supernatant |
| Q62523 | ZYX | Zyxin | C2 | Up in cells and supernatant |
| P14602 | HSPB1 | Heat shock protein beta-1 | C2 | Up in cells and supernatant |
| E9Q3G8 | NUP153 | Nucleoporin 153 | C2 | Up in cells and supernatant |
| Q80YX1 | TNC | Tenascin | C2 | Up in cells and supernatant |
| Q3TJD7 | PDLIM7 | PDZ and LIM domain protein 7 | C2 | Up in cells and supernatant |
| P48036 | ANXA5 | Annexin A5 | C2 | Up in cells and supernatant |
| Q3UTJ2-2 | SORBS2 | Sorbin and SH3 domain-containing protein 2 | C2 | Up in cells and supernatant |
| E0CYH7 | FILIP1L | Filamin A-interacting protein 1-like | C2 | Up in cells and supernatant |
| A1BN54 | ACTN1 | Actinin alpha 1 | C2 | Up in cells and supernatant |
| Q9DBJ1 | PGAM1 | Phosphoglycerate mutase 1 | C2 | Up in cells and supernatant |
| P10107 | ANXA1 | Annexin A1 | C2 | Up in cells and supernatant |
| Q99K51 | PLS3 | Plastin-3 | C2 | Up in cells and supernatant |
| Q8BTM8 | FLNA | Filamin A | C2 | Up in cells and supernatant |
| E9PYF4 | LMO7 | LIM domain only protein 7 | C2 | Up in cells and supernatant |
| Q80X90 | FLNB | Filamin B | C2 | Up in cells and supernatant |
| Q91Y82 | KLK6 | Kallikrein-related peptidase 6 | C3 | Down in cells and supernatant |
| Q8R116 | NOTUM | Palmitoleoyl-protein carboxylesterase NOTUM | C3 | Down in cells and supernatant |
| P01027 | C3 | Complement C3 | C3 | Down in cells and supernatant |
| E9Q600 | LOXL4 | Lysyl oxidase-like 4 | C3 | Down in cells and supernatant |
| Q61739 | ITGA6 | Integrin alpha-6 | C3 | Down in cells and supernatant |
| Q8CG19 | LTBP1 | Latent TGF-beta-binding protein 1 | C3 | Down in cells and supernatant |
| Q62179 | SEMA4B | Semaphorin-4B | C3 | Down in cells and supernatant |
| Q60963 | PLA2G7 | Platelet-activating factor acetylhydrolase | C3 | Down in cells and supernatant |
| Q9ER10 | PRSS22 | Serine protease 22 | C3 | Down in cells and supernatant |
| Q3UUY6 | PROM2 | Prominin-2 | C3 | Down in cells and supernatant |
| Q91ZX7 | LRP1 | LDL receptor-related protein 1 | C3 | Down in cells and supernatant |
| P33435 | MMP13 | Matrix metalloproteinase 13 | C3 | Down in cells and supernatant |
| G5E874 | LAMC2 | Laminin subunit gamma 2 | C3 | Down in cells and supernatant |
| Q61001 | LAMA5 | Laminin subunit alpha 5 | C3 | Down in cells and supernatant |
| P06909 | CFH | Complement factor H | C3 | Down in cells and supernatant |
| E9PZ16 | HSPG2 | Perlecan | C3 | Down in cells and supernatant |
| Q3UQ28 | PXDN | Peroxidasin | C3 | Down in cells and supernatant |
| A2A864 | ITGB4 | Integrin beta-4 | C3 | Down in cells and supernatant |
| Q00780 | COL8A1 | Collagen alpha-1(VIII) chain | C3 | Down in cells and supernatant |

**Supplementary Table 2:**
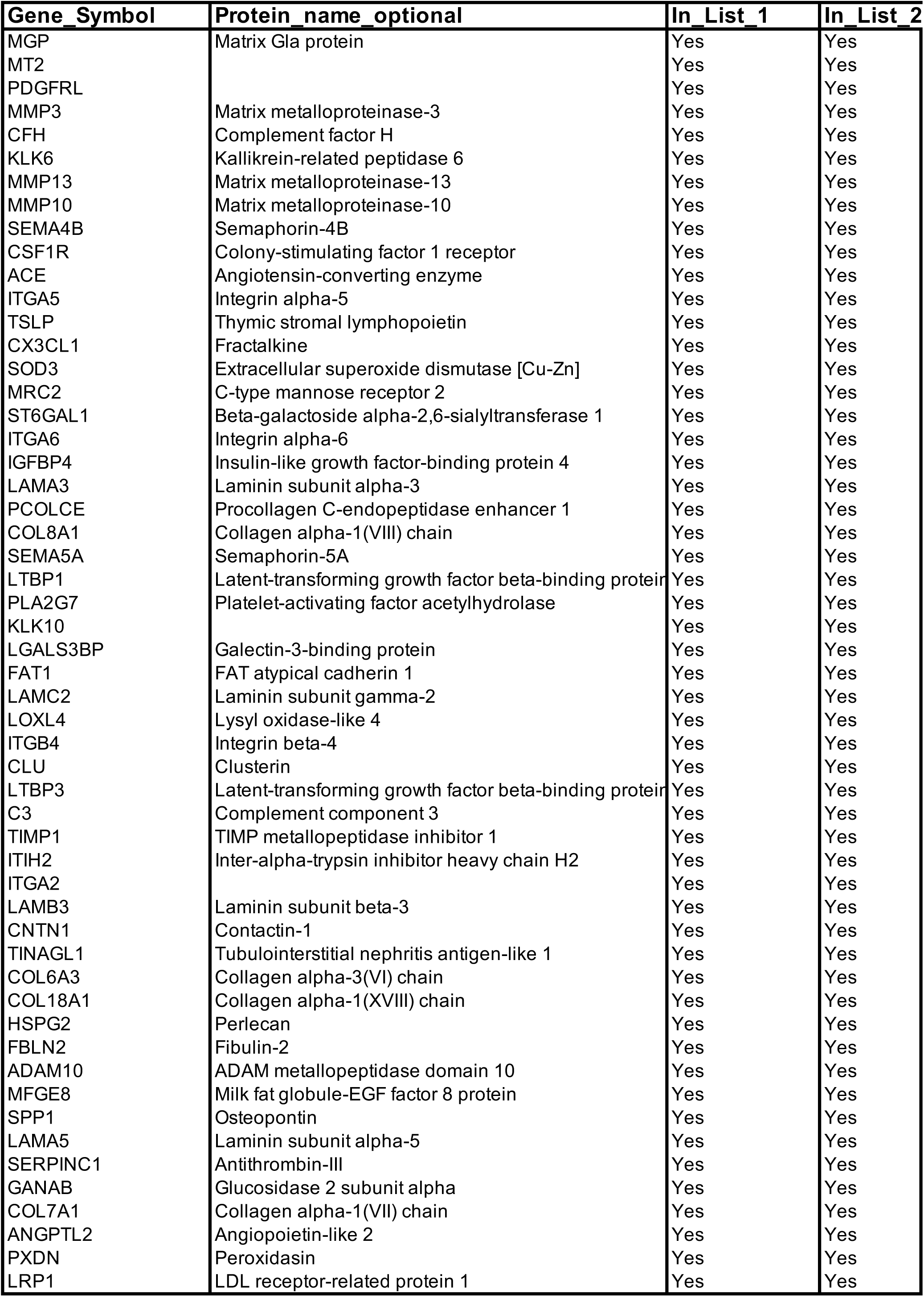
Overlapping genes/proteins (List FAK-dependent genes ∩ List publicly available datasets).

**Supplementary Table 3:**
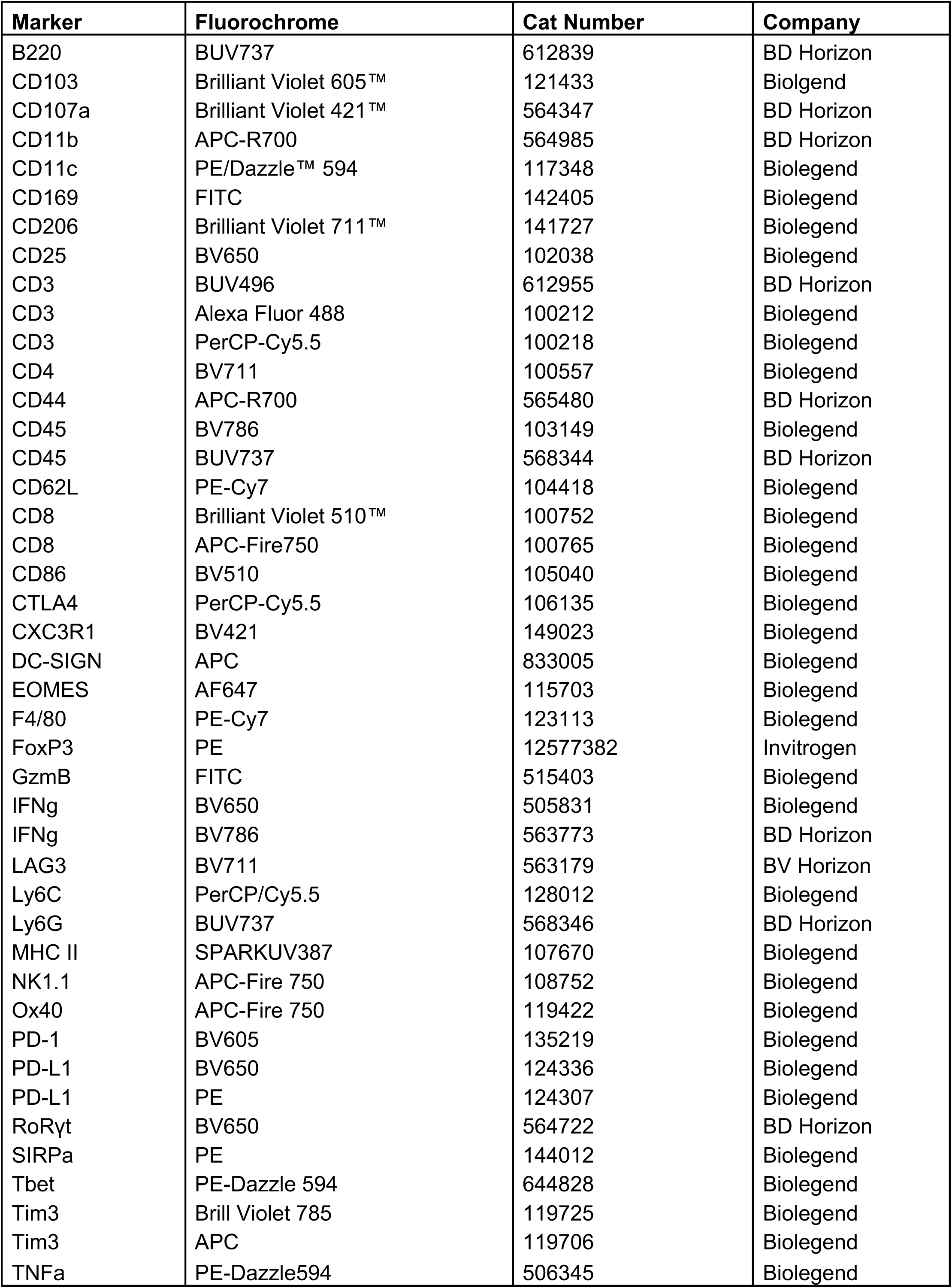
Antibody List for Flow Cytometry staining.

